# A tissue-targeted auxin-inducible degradation toolkit for *C. elegans* embryogenesis

**DOI:** 10.64898/2026.10.07.757239

**Authors:** Neha Varshney, Rebecca Green, Ayla O. Desai, Shabnam Moghareh, Dylan Green, Arshad Desai, Karen Oegema

## Abstract

Many proteins required for embryogenesis are expressed in multiple tissues and across developmental time, often with tissue- and stage-specific roles. To resolve where and when proteins function, we developed a tissue-targeted auxin-inducible degradation toolkit for *C. elegans* embryogenesis. We generated a plasmid set compatible with single-copy insertion or high-copy extrachromosomal array generation that employs tissue-specific promoters (active in the germline, intestine, muscle, pharynx, neuroblasts, neurons, sensory neurons, epidermis, and pan soma) to drive an operon expressing an optimized auxin-dependent plant F-box factor TIR1 and, in most cases, a fluorescent histone. Using a ubiquitous nuclear envelope marker as a test substrate, we identified single-copy insertions and arrays that exhibit efficient auxin-dependent, tissue-specific target degradation. We also establish a workflow that ensures all functionally important isoforms of a target are tagged and that degradation reproduces the loss-of-function phenotype prior to tissue-specific analysis. Applying the toolkit to the broadly expressed Rho kinase LET-502, we find that degradation in the epidermis, but not in other embryonic tissues, disrupts elongation. Unexpectedly, temporal analysis revealed that LET-502 is required for late as well as early elongation.

**SUMMARY STATEMENT:** A toolkit for auxin-mediated tissue- and stage-specific degradation of essential targets during *C. elegans* embryogenesis reveals that the contractility regulator LET-502 is required in the epidermis for late as well as early elongation.

## INTRODUCTION

The *C. elegans* embryo is transparent and develops from a single cell into a 558-cell larva containing all major somatic tissues within an ∼14-hour window (Corsi et al., 2015; Sulston et al., 1983). *C. elegans* offers several technical advantages for studies of metazoan embryogenesis, including ease of genetic manipulation (Brenner, 1974; Corsi et al., 2015), CRISPR/Cas9 for *in situ* gene tagging (Dickinson et al., 2013; Friedland et al., 2013; Paix et al., 2015), and public datasets characterizing its genome, cell lineage, and gene expression (Consortium, 1998; Murray et al., 2006; Packer et al., 2019; Sulston et al., 1983). The experimental tractability of *C. elegans* has enabled studies ranging from elucidating signaling and transcriptional networks that define cell polarity and fate (Guan et al., 2022; Jackson and Eisenmann, 2012; Lam and Phillips, 2017; Murray, 2018; Packer et al., 2019; Priess, 2005; Rose and Gonczy, 2014) to dissecting mechanical and physiological processes underlying development (Carvalho and Broday, 2020; Chisholm and Hsiao, 2012; Goldstein and Nance, 2020; Lamkin and Heiman, 2017; Vuong-Brender et al., 2016).

In addition to genetic approaches, RNA interference (Fire et al., 1998; Timmons and Fire, 1998) has enabled systematic high-content screens that have defined a network of ∼800 *C. elegans* genes required for embryo production and the first two embryonic cell divisions (Gonczy et al., 2000; Green et al., 2011; Piano et al., 2000; Piano et al., 2002; Sonnichsen et al., 2005; Zipperlen et al., 2001). These screens also identified ∼1900 candidates for genes with roles in embryonic development; screening 500 of these revealed clear functions during embryogenesis for ∼40% of the genes that had not previously been implicated in development (Green et al., 2024). Collectively, these systematic functional screens suggest that, including the genes required for embryo production and the early embryonic divisions, more than 1500 genes are required for robust embryogenesis. With the expansion of the catalog of genes required for embryonic development, a major challenge is to resolve where, when, and how the proteins encoded by these genes contribute to embryogenesis. Although gene expression profiles can pinpoint where a gene acts—as they have for lineage and cell type-specific transcription factors (Murray et al., 2012; Packer et al., 2019)—many genes critical for embryogenesis are widely expressed across multiple tissues and developmental stages, making it unclear where and when they perform their essential functions. Moreover, disrupting a gene’s early function can mask the roles it plays later in embryogenesis. Thus, addressing the functions of widely expressed genes in the different aspects of embryogenesis requires the ability to disrupt their function with tissue and temporal specificity.

Systems employing conditional CRISPR/Cas9 (Shen et al., 2014) and tissue-specific RNAi (Qadota et al., 2007) have been used to restrict perturbations to defined cell populations. CRISPR/Cas9 can be rendered tissue specific by driving Cas9 from tissue-specific or inducible promoters, and RNAi by restoring RNAi competence in specific tissues in an otherwise RNAi-deficient background. However, because these methods act at the DNA or RNA levels, pre-existing protein can persist and delay or prevent the detection of phenotypes. Strategies that operate at the protein level include tagging proteins at their endogenous locus with degrons that trigger proteasomal degradation in the presence of a tissue-specific “degrader”. In one of these strategies, proteins are tagged with a 36-amino acid ZF1 degron that is recognized by the E3 ligase substrate-recognition adapter ZIF-1 (DeRenzo et al., 2003). Combining a *zif-1* mutant background, in which ZF1-tagged proteins are stable, with tissue-specific re-expression of ZIF-1 allows selective depletion in chosen lineages (Armenti et al., 2014; Sallee et al., 2018). In a related strategy that does not require a *zif-1* mutant background, an anti-GFP nanobody fused to ZIF-1 is expressed under tissue-specific promoters, promoting the degradation of GFP-tagged proteins where the degrader is expressed (Wang et al., 2017). A drawback of both approaches is that, because the degrader is constitutively active, animals homozygous for the degron-tagged target and carrying the degrader cannot be maintained if degradation is lethal; phenotypes must instead be scored after crossing target and degrader strains in the 25% of F2 progeny that are homozygous for the tagged target. The widely adopted auxin-inducible degron (AID) system circumvents these challenges by using a heterologous, plant-derived substrate-recognition adapter (TIR1) that engages AID degron-tagged target proteins only when the small-molecule auxin is present (Negishi et al., 2022; Nishimura et al., 2009; Yesbolatova et al., 2020; Zhang et al., 2015). Because degradation only occurs when auxin is added, animals carrying TIR1 that are homozygous for the degron-tagged target can be maintained as a viable stock.

In *C. elegans*, the *A. thaliana* F-box protein AtTIR1 has been used to trigger degradation of proteins tagged with a short degron in the presence of auxin (Sepers et al., 2022; Zhang et al., 2015). An improved system, called AID2, uses a bump-and-hole approach that pairs mutant TIR1 (AtTIR1^F79G^) with the synthetic auxin 5-Ph-IAA, enabling degradation at low ligand concentrations and reducing leaky auxin-independent degradation (Hills-Muckey et al., 2022; Negishi et al., 2022; Yesbolatova et al., 2020). An acetoxymethyl ester variant, 5-Ph-IAA-AM, has been shown to penetrate the *C. elegans* eggshell, allowing target degradation to be triggered by compound addition at any desired developmental stage (Negishi et al., 2019; Negishi et al., 2022). A prior tissue-specific *C. elegans* TIR1 toolkit (Ashley et al., 2021) employed the original AID system and was not optimized for degradation in embryonic tissues.

To resolve where and when proteins function in early *C. elegans* embryogenesis, we built on the AID2 system to develop a flexible auxin-inducible degradation toolkit for embryonic tissues designed to drive degradation on timescales relevant to the rapid events of embryogenesis. We generated a set of plasmids compatible with single-copy insertion or high-copy extrachromosomal arrays that uses embryonic tissue-specific promoters to drive an operon expressing TIR1^F79G^ and mCherry::Histone H2B. Using degradation of a ubiquitously-expressed test protein, we identified an extrachromosomal array or single-copy insertion that exhibited efficient tissue-specific degradation. To assess utility of the toolkit, we employed it to analyze the widely expressed Rho kinase LET-502, which has been proposed to act in the epidermis to promote embryo elongation (Diogon et al., 2007; Drewnik et al., 2021; Gally et al., 2009). Degradation in the epidermis, but not in the intestine, pharynx, muscle, neuroblasts, or sensory neurons, reproduced the elongation defect observed following degradation in all somatic tissues. Temporally controlled degradation further revealed that LET-502 is required not only for early (1-2-fold) but also for late (2-3-fold) elongation, a phase when the mechanism of force generation is thought to shift from contractility within the epidermis to the underlying musculature, with cytoskeletal remodeling within the epidermis occurring in response to muscle-driven forces (Carvalho and Broday, 2020; Vuong-Brender et al., 2016). These results suggest that epidermal LET-502 continues to contribute after the muscle-driven phase begins, and more broadly demonstrates the utility of the toolkit for dissecting where and when widely expressed proteins act during *C. elegans* embryogenesis.

## RESULTS

### A plasmid toolkit for expression of a TIR1^F79G^— mCherry::H2B operon as single-copy insertions or extrachromosomal arrays in embryonic tissues

To characterize the tissue-specific functions of proteins required for *C. elegans* embryonic development, we constructed a set of plasmids in which promoters active in different embryonic tissues drive a two-gene operon encoding TIR1^F79G^ and mCherry::H2B joined by an operon linker (hereafter the TIR1^F79G^— mCherry::H2B operon) (**Fig. 1A; Fig. S1A**); the mCherry::H2B reporter provides a visual readout for operon expression. Promoters were selected to enable targeting of AID*-tagged proteins specifically in the germline (P*mex-5*, (Merritt et al., 2008)), intestine (P*elt-2*, (Fukushige et al., 1998)), pharynx (P*cest-17*, also called K07C11.4, (Gaudet et al., 2004; Packer et al., 2019)), muscle (P*hlh-1*, (Krause et al., 1990; Krause et al., 1994)), body wall muscle (P*myo-3*, (Fire and Waterston, 1989; Okkema et al., 1993)), epidermis (P*elt-1* and P*lin-26*, (Labouesse et al., 1996; Landmann et al., 2004; Page et al., 1997)), neuroblasts (P*ztf-11*, (Lee et al., 2019)), neurons (P*cnd-1*, (Hallam et al., 2000)), and sensory neurons (P*dyf-7*, (Heiman and Shaham, 2009)); two pan-somatic promoters were also selected (P*eft-*3, also called P*eef-1A.1*, (Zhang et al., 2015), and P*rps-27* (Adikes et al., 2020)). Because embryogenesis proceeds rapidly and promoters differ substantially in strength, the plasmids were designed to enable single-copy insertion using the MosSCI method (Frokjaer-Jensen et al., 2012; Frokjaer-Jensen et al., 2008; Frokjaer-Jensen et al., 2014) or the generation of higher-expressing multi-copy extrachromosomal arrays by co-injection with the dominant *rol-6* selection marker (**Fig. 1A; Fig. S1A-C**; (Mello et al., 1991)). The two formats allow TIR1^F79G^ levels to be titrated for each promoter to achieve degradation that is both efficient and restricted to the target tissue. For arrays, we generated lines that transmit at high frequency (50-90% roller progeny with mCh::H2B expression in the target tissue; see **Fig. S1C**). Because arrays are not transmitted to all progeny, the batch of embryos from each dissected worm contains both embryos that have and embryos that lack the TIR1^F79G^-expressing array, providing a useful internal control.

**Figure 1.**
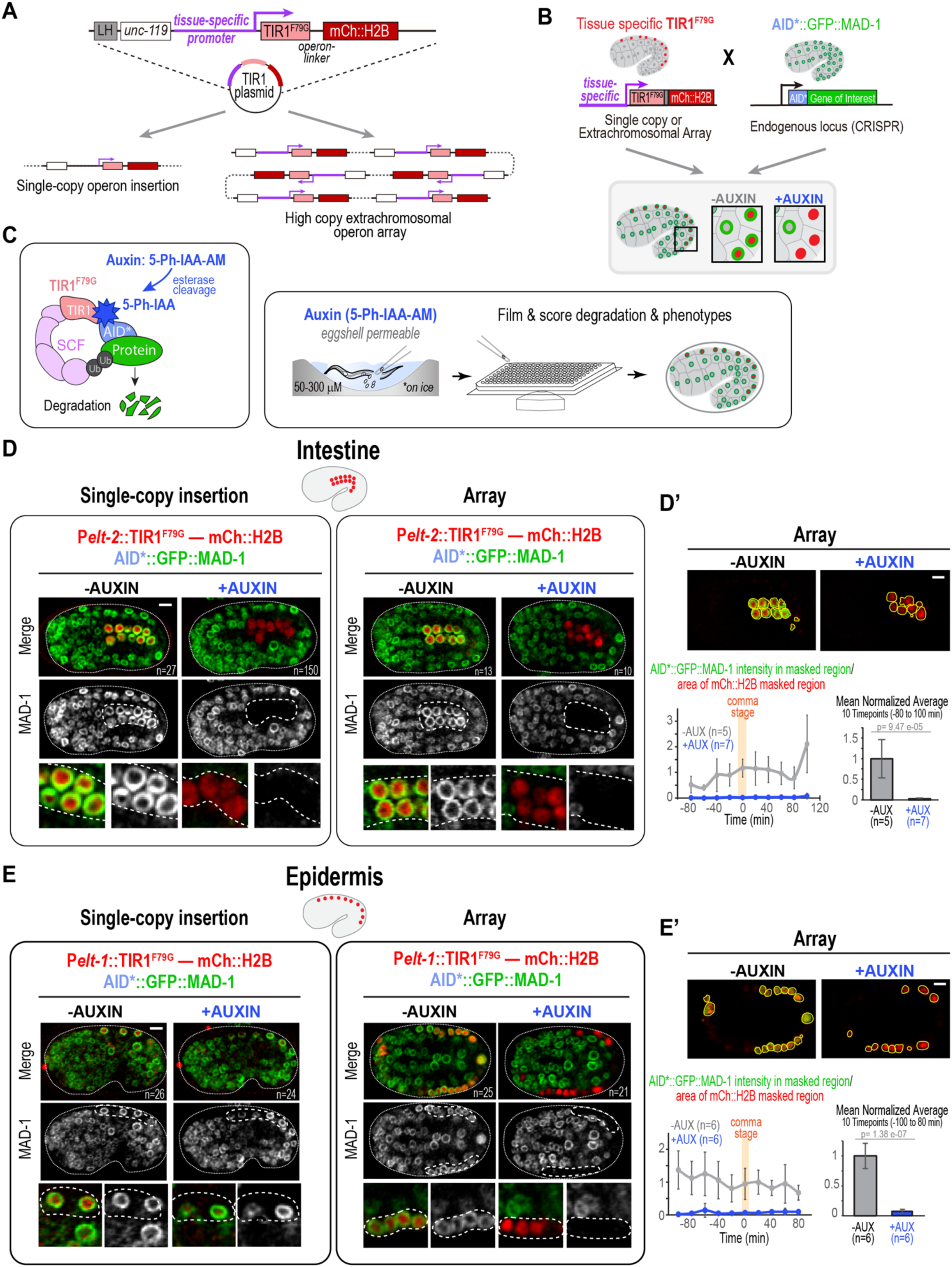
A plasmid toolkit for expression of TIR1^F79G^—mCherry::H2B operons as single-copy insertions or extrachromosomal arrays in embryonic tissues. **(A)** Schematic of the tissue-specific TIR1^F79G^—mCherry::H2B operon expression plasmid, which can be used to generate single-copy insertions or high-copy extrachromosomal arrays. **(B)** To test whether operon expression drives auxin-inducible degradation of AID*::GFP-tagged targets, insertions/arrays were mated into a background containing the broadly expressed nuclear envelope marker AID*::GFP::MAD-1. In operon-expressing cells, mCh::H2B marks the nuclei (*red*) and AID*::GFP::MAD-1 forms a green ring around them; upon auxin addition, the ring disappears if degradation occurs. **(C)** (*left*) Schematic of the auxin-inducible degradation system. The expressed plant F-box protein TIR1^F79G^ assembles with endogenous SCF E3 ligase and engages AID*-tagged targets to promote their ubiquitination and degradation only when bound to the auxin analog 5-Ph-IAA. The eggshell-permeable prodrug 5-Ph-IAA-AM is cleaved by intracellular esterases to release 5-Ph-IAA. (*right*) Assay schematic. Worms are dissected in 50 µM auxin on ice and embryos are transferred by mouth pipette into a 384-well plate. A subset of the embryos is filmed through embryogenesis at 60X, and the whole well is subsequently scanned at 10X to score embryonic lethality and larval phenotypes. **(D-E)** Images compare auxin-mediated degradation of AID*::GFP::MAD-1 for a single-copy insertion (*left*) and an array (*right*) expressing the TIR1^F79G^—mCh::H2B operon under the *elt-2* promoter in the intestine (*D*) or the *elt-1* promoter in the epidermis (*E*). Embryos are shown in the absence (*left*) or presence (*right*) of 50 µM auxin. White dashed lines outline the operon-expressing tissue. Zoomed panels show a higher-magnification view (∼2.5X) of the highlighted region. **(D′, E′)** Quantification of degradation in embryos expressing the TIR1^F79G^—mCh::H2B operon from extrachromosomal arrays in the intestine (D′) or epidermis (E′). For each embryo, total AID*::GFP::MAD-1 intensity was measured within a mask generated by dilating the mCh::H2B signal in a central z-section over the developmental time course. Intensities were normalized by dividing by the mask area at each timepoint and then by the mean of the same ratio in the no-auxin control. Timelapse plots (*left*) show the mean ± SD of the normalized intensity. Times are relative to the comma stage (comma = timepoint 0); gray is −auxin, blue is +auxin. Bar graphs (*right*) show the mean ± SD of the per-embryo averages across a 10-timepoint window (as indicated). *P* values are from two-tailed Welch’s *t*-tests. *n* values in the images in (D and E) are number of imaged embryos and *n* values under the graphs (D’ and E’) are number of embryos analyzed. Scale bars, 5 µm.

To assess tissue-specific degradation, *in situ*-tagged AID*::GFP::MAD-1, a degron tagged nuclear envelope-localized protein expressed in all tissues that is largely dispensable for embryonic development (Rodenas et al., 2012; Wang et al., 2017), was crossed into the promoter-driven TIR1^F79G^— mCh::H2B operon strains (**Fig. 1B**). The AID*::GFP::MAD-1 fluorescence signal enables quantification of target degradation using image segmentation based on nuclear mCh::H2B after addition of eggshell-permeable 5-Ph-IAA-AM ((Negishi et al., 2022); hereafter "auxin"; **Fig. 1C**). Both single copy and extrachromosomal array formats of the TIR1^F79G^— mCh::H2B operon were generated and tested for five representative promoters—P*elt-2* (intestine), P*elt-1* (epidermis), P*cnd-1* (neurons), P*cest-17* (pharynx), and P*hlh-1* (muscle). In each case, auxin-dependent AID*::GFP::MAD-1 degradation was observed only in cells expressing the mCh::H2B operon marker (**Fig. 1D,E; Fig. 2A**). Although three of the promoters (P*elt-2*, P*cest-17*, P*hlh-1*) were effective and specific in both configurations, trade-offs between the formats were apparent for the remaining two (P*elt-1*, P*cnd-1*). Three representative examples illustrate observed outcomes (**Fig. 1D, D’, E, E’; Fig. S1D**). For P*elt-2*, the single copy insertion and array gave similar results: mCh::H2B expression was restricted to the centrally positioned intestinal cells (**Fig. 1D**; (Fukushige et al., 1998)), where AID*::GFP::MAD-1 was robustly and specifically degraded upon auxin addition (**Fig. 1D, D’, Movie S1**). For P*elt-1*, the single-copy insertion produced dim mCh::H2B signal and failed to support efficient AID*::GFP::MAD-1 degradation, whereas the multi-copy extrachromosomal array drove bright mCh::H2B expression in peripheral nuclei, consistent with epidermal expression, and robust AID*::GFP::MAD-1 degradation in the presence of auxin (**Fig. 1E, E’, Movie S1**). In contrast, the single-copy insertion of P*cnd-1* drove bright, neuron-specific mCh::H2B expression and auxin-dependent degradation, whereas the array resulted in non-specific expression and degradation across the majority of the embryo (**Fig. S1D**). Overall, arrays provide robust expression and degradation, which is especially useful for weaker promoters such as P*elt-1*, whereas single-copy insertions may better preserve the tissue-specificity of stronger promoters such as P*cnd-1*.

**Figure 2.**
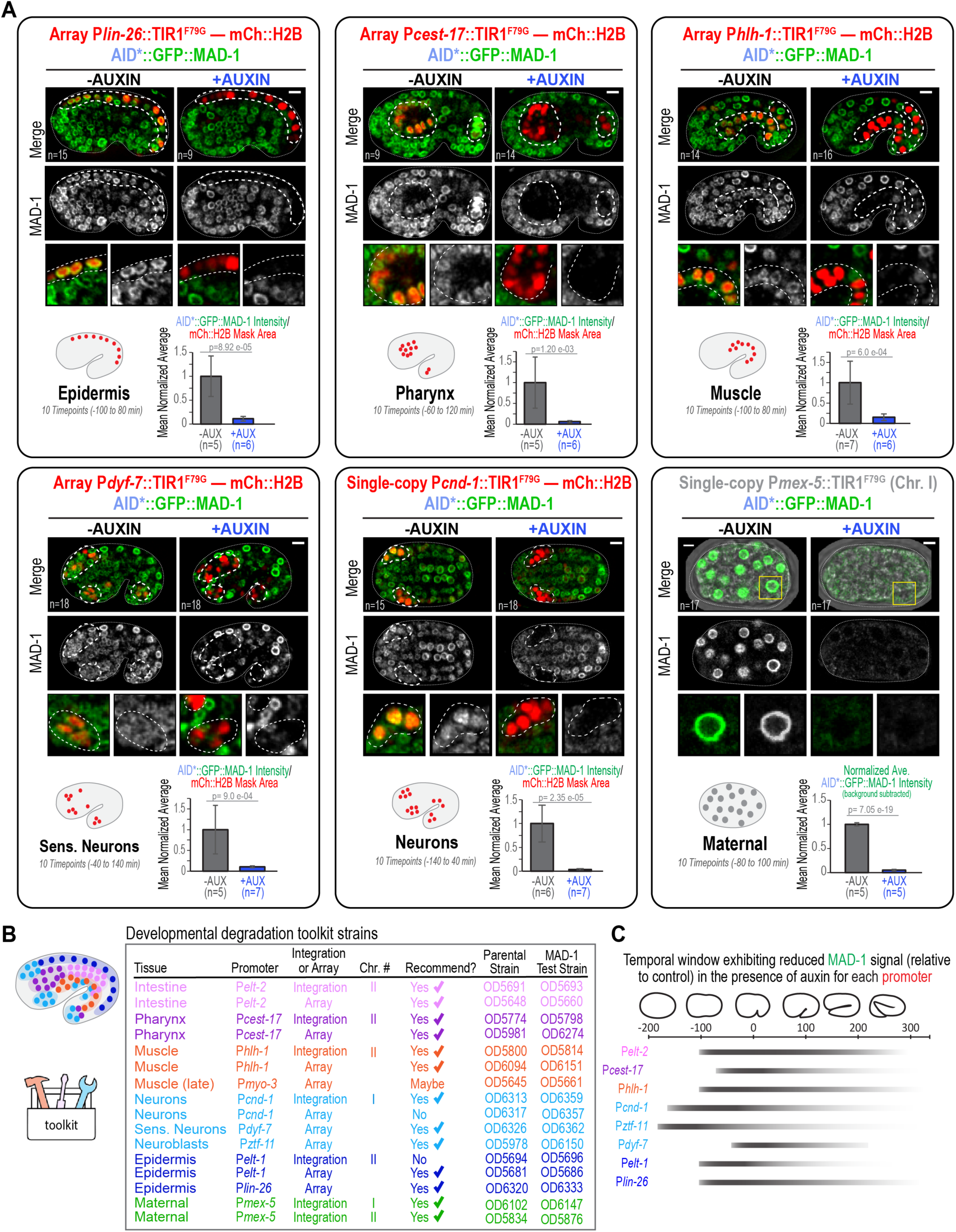
A tissue-specific degradation toolkit spans the major embryonic tissues. **(A)** Representative images of auxin-mediated AID*::GFP::MAD-1 degradation for array strains targeting the epidermis (P*lin-26*), pharynx (P*cest-17*), muscle (P*hlh-1*), and sensory neurons (P*dyf-7*), and for the single-copy insertion strains targeting neurons (P*cnd-1*) and the germline (P*mex-5*). Images in the absence (*left*) or presence (*right*) of 50 μM auxin are shown. Schematics indicate the positions of the TIR1^F79G^— mCh::H2B operon-expressing cells. (*top rows*) Merge with AID*::GFP::MAD-1 in green and the mCh::H2B operon reporter in red—or, for the maternal P*mex-5* insertion that lacks mCh::H2B, a brightfield overlay. (*middle rows*) Grayscale images of AID*::GFP::MAD-1. White dashed lines outline the operon-expressing tissue. (*bottom rows*) Zoomed panels show a higher-magnification view (∼2.5X) of the highlighted region. Bar graphs in the lower right corners show the mean ± SD of the per-embryo average of the normalized AID*::GFP::MAD-1 intensity across the indicated 10-timepoint window, quantified as described for *Fig. 1D’,E*’; or, for the P*mex-5* operon the average background-subtracted intensity (see *Fig. S2D*). *n* values in images are number of imaged embryos and *n* values under the graphs are number of embryos analyzed; at least two independent experiments were performed per condition. *P* values are from two-tailed Welch’s t-tests. Scale bars, 5μm. **(B)** Table summarizing toolkit strains. (*left*) Schematic shows the positions of tissues targeted by toolkit promoters. Tissue colors match color-coded text in the table: intestine (*pink*), pharynx (*purple*), muscle (*orange*), neurons (*sky blue*), epidermis (*navy blue*), maternal/germline (*green*). Table lists target tissue, promoter, whether the strain is a single-copy integration or array, and strain numbers for the parental strain and its AID*::GFP::MAD-1-containing derivative, which can be used for optimizing degradation conditions. Check marks indicate recommended strains. The *Pcnd-1* array and *Pelt-1* single-copy insertion are not recommended because they respectively drive non-specific degradation outside neurons (*Fig. S1D*) and express TIR1^F79G^ at levels too low for efficient degradation (Fig. 1E). **(C)** Schematic approximating time windows when mCh::H2B is visible and AID*::GFP::MAD-1 is degraded (see *Fig. 1D’&E’ and Fig. S2C*). In some cases, the onset of AID*::GFP::MAD-1 degradation starts 20–40 minutes before mCh::H2B becomes visible, which is not captured in this schematic, likely because mCherry matures more slowly than GFP. The tapering of each degradation window is specific to MAD-1 dynamics and may not generalize to other targets, whose turnover rates and expression levels may differ.

### Building a tissue-specific degradation toolkit that spans the major embryonic tissues

Next, we identified optimal extrachromosomal arrays and/or single-copy insertions that directed tissue-specific operon expression and AID*::GFP::MAD-1 degradation for 10 promoters spanning the major embryonic tissues and germline. In addition to the previously highlighted intestinal (P*elt-2*), early epidermal (P*elt-1*), neuronal (P*cnd-1*), pharyngeal (P*cest-17*), and muscle (P*hlh-1*) promoters, extrachromosomal arrays were generated for the pan-epidermal (P*lin-26*), body wall muscle (P*myo-3*), sensory neuron (P*dyf-7*), and neuroblast (P*ztf-11*) promoters; with the exception of P*myo-3*, we could confirm that each drove strong, tissue-specific mCh::H2B expression and efficient AID*::GFP::MAD-1 degradation in the presence of auxin (**Fig. 2A; Fig. S2A–C**). The P*myo-3* operon may be useful, but it turned on relatively late and its degradation did not appear to be as efficient as for the other promoters (**Fig. S2B**).

Because extrachromosomal arrays are silenced in the germline (Kelly et al., 1997) two independent single-copy insertions of the P*mex-5* germline promoter-driven TIR1^F79G^— mCh::H2B operon were generated at the ttTi5605 (chromosome II) and oxTi185 (universal “Uni I” chromosome I) MosSCI sites that both drove efficient AID*::GFP::MAD-1 degradation of maternally loaded proteins in the presence of auxin (**Fig. 2A**, **Fig. S2D**); to maximize compatibility with fluorescent markers these insertions do not direct mCh::H2B expression. Because the P*ztf-11* promoter is expressed in neuroblasts, which are progenitors rather than differentiated neurons, degradation occurred across much of the early embryo. Phenotypes obtained with this strain may therefore reflect a requirement for the protein in neuroblast proliferation rather than a tissue-specific function. The full panel of tissue-specific single-copy insertion and array strains is summarized in **Figure 2B**, which lists the target tissue, promoter, and whether the single-copy insertion and/or array are recommended (also see **Movie S2;** for pan-tissue strains see **Fig. S3A)**.

Quantification of the AID*::GFP::MAD-1 signal over time in the targeted tissues defined a temporal window when robust AID*::GFP::MAD-1 degradation was observed in the array strains for each promoter (P*dyf-7*, P*cest-17*, P*elt-2*, P*hlh-1*, P*lin-26*, P*elt-1*, and P*ztf-11*), as well as for the *Pcnd-1*-driven single-copy insertion (**Fig. 1D′,E′; Fig. S2A-D**). As a guideline, we generated a qualitative summary of the temporal windows when cells expressed mCherry and exhibited robust reduction of AID*::GFP::MAD-1 signal in the strains for each promoter (**Fig. 2C**). When using this as a reference, note that in some cases the onset of AID*::GFP::MAD-1 degradation became apparent ∼20–40 minutes before mCh::H2B became visible, likely because mCherry matures more slowly than GFP. Also note that the tapering of the degradation windows reflects the dynamics of MAD-1 and may not generalize to other targets with different expression kinetics or stability. The two epidermal promoters, P*elt-1* and P*lin-26*, provide an interesting example of how kinetic information is useful. *elt-1* encodes a GATA factor that initiates epidermal specification (Gilleard and McGhee, 2001; Page et al., 1997), whereas *lin-26*, which is activated downstream of *elt-1* (Landmann et al., 2004), is expressed throughout development and maintains epidermal fate (Labouesse et al., 1996; Labouesse et al., 1994; Landmann et al., 2004; Packer et al., 2019). Consistent with this, both arrays began robust degradation ∼100 min before the comma stage, but P*lin-26* supported sustained degradation to ∼250 min after comma, versus only ∼100 min after comma for P*elt-1* (**Fig. 2C**).

Collectively, the toolkit strains provide optimized tissue-specific expression of a TIR1^F79G^— mCh::H2B operon in the major embryonic tissues and germline. The strains are easy to maintain and enable dissection of tissue- and stage-specific protein function during *C. elegans* embryogenesis.

### A workflow for assessing the tissue-specific functions of developmentally essential target genes

Next, we developed a workflow for using the toolkit to analyze the tissue-specific function of targets essential for embryogenesis (**Fig. 3A**). The first step is to generate two *in situ*-tagged versions of the gene of interest, tagged at either the N- or C-terminus: one fused to AID*::GFP (or GFP::AID*) and a second fused to GFP alone (a different fluorescent protein can be substituted for GFP). Upon auxin treatment, the AID*::GFP-tagged protein will be degraded whereas the GFP-tagged protein will not, making it straightforward to distinguish phenotypes resulting from target degradation from those caused by tagging itself or the auxin treatment protocol. Target genes can have multiple splice isoforms and/or use different translational start sites, which may not be properly annotated. Thus, the second step is to confirm that all of the functionally important isoforms are tagged. For this purpose, we recommend using RNA interference (RNAi) to target either the tag (e.g. GFP) or a region of the gene body common to all isoforms and comparing the observed phenotypes. If the tag and gene body RNAi result in similar phenotypes, it is likely that the major functionally important isoforms are tagged and the fusions are suitable for auxin-mediated degradation. If RNAi of the tag results in a weaker phenotype or no phenotype compared to targeting the gene body, one or more functionally important isoforms may lack the tag (see example in **Fig. 3A**). This outcome requires trying a different tagging strategy (for example, at the opposite end of the gene) followed by re-testing by RNAi. Once the tag/gene RNAi comparison indicates that all relevant isoforms have been tagged, the third step is to cross the tagged gene into a strain expressing TIR1^F79G^ under a pan-tissue promoter (**Fig. 3A**; described below). Degrading the encoded protein across all somatic tissues provides a useful check that auxin-mediated degradation can reproduce the RNAi or genetic loss-of-function phenotype. Following these three steps, the tagged gene can be crossed into desired toolkit strains to assess the phenotypic consequences of degradation in specific tissues and/or at specific developmental stages (**Fig. 3A**). This workflow ensures that conclusions derived from use of the toolkit are rigorous. For strains built using arrays, the ability to compare embryos containing versus lacking the array, dissected from the same parent, provides an additional confirmation that any observed effects are related to target degradation.

**Figure 3.**
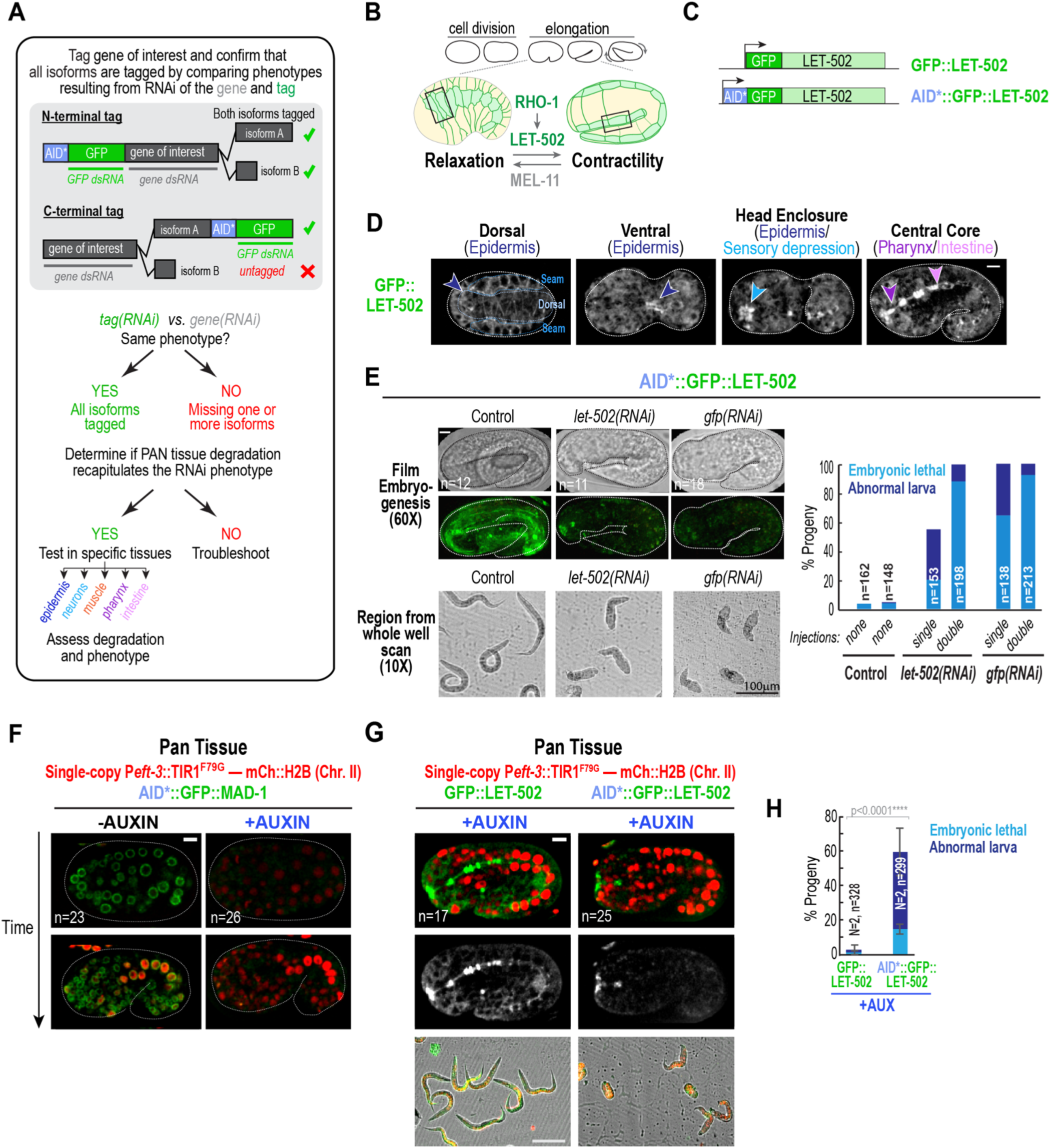
A workflow for assessing the tissue-specific functions of developmentally essential target genes. **(A)** Flowchart outlines a protocol for assessing the tissue-specific functions of an essential gene of interest. The gene is tagged at its N- or C-terminus with an AID*::GFP tag, and phenotypes resulting from RNAi targeting the gene and tag are compared. A weaker phenotype for RNAi targeting the tag compared to the gene suggests an untagged isoform. After all isoforms are tagged, the tagged gene is crossed into a pan-tissue TIR1^F79G^—mCh::H2B operon strain to confirm that auxin-induced degradation across all tissues can recapitulate the RNAi phenotype. Finally, the tagged gene is introduced into tissue-specific toolkit strains to determine where the protein functions. **(B)** Schematic highlights the proposed role of the Rho kinase, LET-502, in embryo elongation. **(C)** Schematic shows the test gene, *let-502*, tagged at its N-terminus with AID*::GFP, or GFP as a negative control. **(D)** Z-sections from GFP::LET-502-expressing embryos showing localization in dorsal and ventral epidermis (navy arrows, presumptive epidermal junctions), the sensory depression (sky-blue arrow), and the central core of the embryo (pharynx, purple arrow; intestine, pink arrow). **(E)** Comparison using RNAi to target the gene or tag in a strain expressing AID*::GFP::LET-502. (*top left*) Paired 60X brightfield and fluorescence images show AID*::GFP::LET-502 depletion in embryos resulting from the single-injection protocol. (*bottom left*) Region from whole well 10X scans show normal hatched larvae in the control and abnormal short larvae and dead embryos in the *let-502(RNAi)* and *gfp(RNAi)* single-injection protocol conditions. *(right)* Graph plots percentage of embryos resulting from the *let-502(RNAi)* and *gfp(RNAi)* single and double-injection protocol conditions that were scored as embryonic-lethal (sky blue) or abnormal larvae (short or abnormally shaped; navy blue). **(F)** Images showing AID*::GFP::MAD-1 (*green*) and the mCh::H2B operon reporter (*red*) at early and mid-developmental timepoints in the absence or presence of 50 µM auxin for the P*eft-3* operon insertion on Chr. II (pan-tissue). The Chr. II insertion drives early, sustained degradation across all tissues, including the germline. **(G)** GFP::LET-502 or AID*::GFP::LET-502 were imaged in the background of the pan-tissue P*eft-3* operon on Chr. II in the presence of 50 µM auxin. (*top*) Color overlay and (*middle*) tagged LET-502 alone, showing a single central plane (60X); (*bottom*) region from 10X whole-well scans. **(H)** Graph plots the mean percentage of embryonic lethality (*sky blue*) and abnormal larvae (*navy blue*). Error bars, SD. *P* value, chi-square test. *n* is the number of progeny analyzed and *N* is the number of replicate experiments. Scale bars, 5 µm (60X images) or 100 µm (10X images in E and G).

### Pan-tissue degradation of the Rho kinase LET-502 leads to an embryo elongation defect

To examine the utility of the toolkit, we focused on LET-502, the *C. elegans* ortholog of Rho kinase, which is broadly expressed and essential for embryo elongation. LET-502 promotes actomyosin contractility by phosphorylating the myosin regulatory light chain MLC-4 and antagonizing the MEL-11 phosphatase that removes this phosphorylation (**Fig. 3B**; (Amano et al., 2010; Gally et al., 2009; Piekny and Mains, 2002; Piekny et al., 2000; Shelton et al., 1999; Wissmann et al., 1999; Wissmann et al., 1997). LET-502 loss reduces the rate of contractile ring constriction during the early embryonic cell divisions (Maddox et al., 2007; Piekny and Mains, 2002; Piekny et al., 2000) and perturbs embryo elongation (Diogon et al., 2007; Gally et al., 2009; Piekny et al., 2000; Wissmann et al., 1997). Although LET-502 is broadly expressed in embryos (**Fig. 3D**), it has been proposed to function primarily in the epidermis (Diogon et al., 2007; Drewnik et al., 2021; Gally et al., 2009).

Following the workflow above, we generated strains expressing GFP::LET-502 and AID*::GFP::LET-502 (**Fig. 3C**). Both fusions localized to the dorsal/ventral and lateral seam cells in the epidermis where they concentrated at cell-cell boundaries (**Fig. 3D**; (Piekny et al., 2003)). GFP fusions with LET-502 were also enriched in the pharynx/intestine region, and in the sensory depression (the future mouth; **Fig 3D**) where neuronal dendrite tips and their associated sensilla pores converge during head enclosure (Grimbert et al., 2021; Kunz et al., 2021). Thus, LET-502 localizes to multiple tissues during embryogenesis. To determine whether all functionally important LET-502 isoforms were tagged, we compared phenotypes resulting from RNAi targeting the GFP coding region versus targeting the *let-502* gene body in the AID*::GFP::LET-502 strain background (**Fig. 3E**). For this, we employed either a single-injection or a more stringent double-injection protocol. In the double-injection protocol, both the maternally loaded and zygotically expressed proteins are strongly depleted, whereas in the single injection protocol zygotic expression is strongly inhibited, but a larger fraction of the maternally-loaded protein remains, compared to the double-injection protocol. Both *let-502(RNAi)* and *gfp(RNAi)* resulted in comparable depletion of visible AID*::GFP::LET-502. Embryos from the single-injection protocol for either dsRNA gave a mixture of embryos arresting at or before the 1.5–2-fold stage or hatching as markedly shortened or misshapen larvae. Embryos from the double-injection protocol led to the majority of embryos arresting at or before the 1.5–2-fold stage (embryonic lethality; **Fig. 3E**, **Movie S3**). Thus, RNAi targeting the GFP tag produced a phenotype at least as severe as targeting the *let-502* gene, indicating that the functionally important isoforms are tagged.

The next step in the workflow is to determine whether auxin-inducible degradation across all somatic tissues recapitulates the RNAi phenotype. To test this, we generated single-copy insertions on Chr. I and Chr. II that express the TIR1^F79G^ — mCherry::H2B operon under the control of the P*eft-3* promoter (*eft-3* is a broadly expressed translational elongation factor; **Fig. 3F; Fig. S3A-C;** (Zhang et al., 2015)). In tests of AID*::GFP::MAD-1 degradation, the P*eft-3* single copy operon insertion on Chr. II exhibited robust target degradation in both the germline and the soma (**Fig. 3F**), whereas the P*eft-3* transgene insertion on Chr. I supported degradation only in the soma (**Fig. S3A-C**), likely due to germline silencing. Next, we generated strains homozygous for the P*eft-3*-driven operon on Chr. II and the endogenous GFP::LET-502 or AID*::GFP::LET-502 loci, which are on Chr. I, and imaged embryos in the presence of 50 µM auxin (**Fig. 3G**). As anticipated, levels of AID*::GFP::LET-502 were markedly reduced in the presence of auxin, whereas levels of GFP::LET-502 were not (**Fig. 3G**). Auxin-treated embryos from the AID*::GFP::LET-502 strain exhibited elongation defects similar to reducing LET-502 levels by RNAi using the single injection protocol (compare **Fig. 3G,H** with **Fig. 3E**). Levels of embryonic lethality were not as high as for the double injection protocol, which may be because LET-502 loss also slows the rate of cytokinesis (Maddox et al., 2007; Piekny and Mains, 2002) which might synergize with direct effects on elongation; these issues may be mitigated by the slightly later depletion in auxin-treated embryos compared to double-injection RNAi embryos. A small population of AID*::GFP::LET-502 remained in the sensory depression region in the presence of auxin, suggesting that LET-502 in this region was not fully degraded by the P*eft-*3-driven operon. A second pan-tissue strain, driven by the P*rps-27* promoter (Adikes et al., 2020), is also available as a toolkit reagent but is expressed at lower levels and did sustain degradation of LET-502 efficiently enough to cause an elongation defect (for details see **Fig. S3A, D, E**).

Overall, employing the workflow indicated that AID*::GFP::LET-502 is tagged such that isoforms important for embryo elongation are degraded in the presence of TIR1^F79G^ and auxin, validating its use for embryonic tissue-specific degradation.

### LET-502 is specifically required in the epidermis to drive embryo elongation

Next, we probed the tissue-specific requirements for LET-502 in elongation by crossing the *in situ*-tagged loci expressing GFP::LET-502 and AID*::GFP-tagged LET-502 into toolkit strains expressing the TIR1^F79G^—mCh::H2B operon under different tissue-specific promoters. Embryos from each strain were imaged in the presence of 50 µM auxin, and embryonic lethality and the percentage of abnormal L1 larvae (visually assessed as misshapen or short) were scored for each condition (**Fig. 4A-D**); larval lengths were also measured directly for the epidermal degradation conditions (**Fig. 4E**). In each embryonic tissue, degradation of AID*::GFP::LET-502 but not GFP::LET-502 was observed in cells with mCh::H2B signal. Auxin treatment of embryos expressing P*mex-5*-driven TIR1^F79G^ in the germline largely eliminated AID*::GFP::LET-502 in early embryos until the initiation of zygotic expression; however, this zygotic pool was sufficient to support embryo elongation and viability (**Fig. 4B,D**; (Piekny et al., 2000; Wissmann et al., 1997)). Degradation of LET-502 in the intestine (*Pelt-2*), pharynx (*Pcest-17*), muscle (*Phlh-1*), neuroblasts (*Pztf-11*), or sensory neurons (*Pdyf-7)* did not result in elevated embryonic lethality or larvae with abnormal morphology (**Fig. 4D**). In the case of the P*ztf-11* operon, AID*::GFP::LET-502 expression was substantially reduced throughout much of the embryo early in development (**Fig. 4D**); however, as for the P*mex-5* and Prps-27 driven operons (**Fig. 4B**), LET-502 levels came back up later (*not shown*) and were sufficient to support elongation and viability (**Fig. 4D**). By contrast, LET-502 degradation in the epidermis (P*elt-1* or P*lin-26*) led to elevated rates of embryonic lethality and hatched larvae with short and/or abnormal morphology similar to what was observed following pan-tissue LET-502 degradation or RNAi-mediated depletion (**Fig. 4D,E; Fig. 3E, G, H**). Two controls ensure that this conclusion is rigorous. First, the effects of auxin treatment on embryonic lethality and the percentage of abnormal larvae were not observed in the strains expressing GFP::LET-502, which lack the AID* degron (**Fig. 4D**). Second, because arrays were employed for P*elt-1* and P*lin-26*, we could assess embryos dissected from the same cohort that lacked the array (no epidermal mCh::H2B), which served as in-well auxin-treated controls. Embryos lacking arrays did not exhibit shortened body lengths in the presence of auxin even when they expressed AID*::GFP::LET-502 (**Fig. 4E**). Collectively, these results indicate that LET-502 degradation specifically in the epidermis significantly impairs elongation leading to embryonic arrest prior to hatching or hatched short larvae (**Fig. 4D,E**).

**Figure 4.**
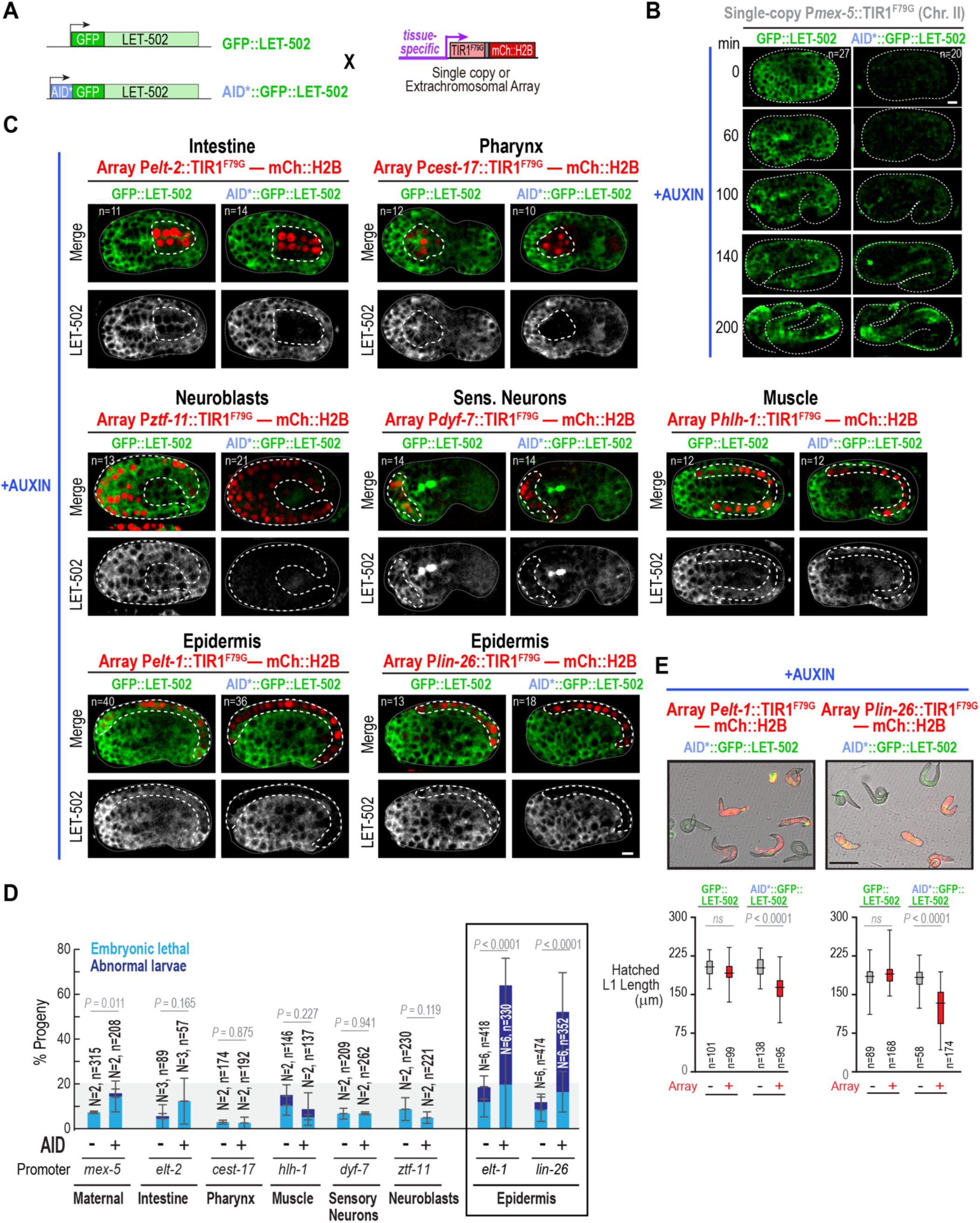
LET-502 is specifically required in epidermal cells to drive embryo elongation. **(A)** Mating scheme used to cross the tagged test (AID*::GFP::LET-502) and control (GFP::LET-502) genes into the tissue-specific TIR1^F79G^—mCh::H2B operon backgrounds. **(B)** Timelapse images showing control GFP::LET-502 (*left*) or AID*::GFP::LET-502 (*right*) expressing embryos in the P*mex-5*::TIR1^F79G^ (Chr.II) background, imaged in the presence of 50 µM auxin (times in minutes). Maternally-loaded AID*::GFP::LET-502 is degraded, but the zygotically transcribed pool turns on midway through development and supports elongation and viability. **(C)** Representative images of GFP::LET-502 (control) and AID*::GFP::LET-502 (test) in the presence of 50 µM auxin, for arrays driving TIR1^F79G^—mCh::H2B operon expression in the intestine (P*elt-2*), pharynx (P*cest-17*), neuroblasts (P*ztf-11*), sensory neurons (P*dyf-7*), muscle (P*hlh-1*), and epidermis (P*elt-1* and P*lin-26*). For each, the top row shows merges (mCh::H2B operon marker, *red*; GFP, *green*) and the bottom row shows GFP in grayscale. White dashed lines outline the operon-expressing tissue. **(D)** Graph shows the mean percentage of progeny scored as embryonic-lethal (sky blue) or abnormal larvae (navy blue) in the presence of 50 µM auxin for GFP::LET-502 (AID−) versus AID*::GFP::LET-502 (AID+) in backgrounds expressing the indicated tissue-specific TIR1^F79G^ — mCh::H2B operons. All operons were in the array format, except for P*mex-5*, which was the single-copy insertion on Chr. II. The shaded band marks the baseline range of lethality/abnormal larvae in the dissect-and-film assay which holds the embryos on ice during setup. Only degradation in the epidermis (P*elt-1*, P*lin-26*; boxed) elevated the phenotype above this baseline. Error bars, SD across independent experiments (weighted by the number of progeny scored per experiment). *P* values, chi-square test (or Fisher’s exact test where a categorical count was <5). **(E)** (*top*) Representative images from 10X whole-well scans of embryos from AID*::GFP::LET-502 expressing mothers carrying the P*elt-1* (*left*) or P*lin-26*-driven (*right*) TIR1^F79G^—mCh::H2B operon arrays in the presence of 50 µM auxin. Larvae lacking the array (*green*) appear normal length, whereas larvae carrying the array (*red*) are shorter and abnormal. (*bottom*) Plots show hatched L1 length (traced along the long axis) for GFP::LET-502 and AID*::GFP::LET-502 expressing embryos lacking (−) or carrying (+) the array, for each promoter. Box and whiskers (min-max) plots show the median and interquartile range. *P* values, Kruskal-Wallis test with Dunn’s multiple-comparisons test. *n* is the number of progeny analyzed; *N* is number of replicates (at least two per condition). Scale bars, 5μm (60X images in B, C) or 100μm (10X image in E).

### The penetrance of epidermal LET-502 degradation phenotypes is elevated at higher concentrations of embryo-permeable auxin

Epidermal LET-502 degradation induced by addition of 50 µM embryo-permeable auxin analog (5-Ph-IAA-AM; referred to here as “auxin”) caused robust but incompletely penetrant elongation phenotypes (**Fig. 4D,E**). Although temperature-sensitive alleles suggest some intrinsic variability in the elongation effects of LET-502 loss (Drewnik et al., 2021; Martin et al., 2014; Vanneste et al., 2013), variability could also arise from partial degradation due to less-than-optimal auxin levels. The acetoxymethyl (AM) modification allows 5-Ph-IAA-AM to pass through the eggshell and enter the embryo where endogenous esterases remove this group to generate 5-Ph-IAA, the compound that bridges TIR1^F79G^ to the AID degron (**Fig. 1C**; (Negishi et al., 2019; Negishi et al., 2022)). The concentration of 5-Ph-IAA inside the embryo therefore depends on the rate at which the compound crosses the eggshell; although 50 µM 5-Ph-IAA-AM is well-tolerated (Negishi et al., 2022) and sufficient for complete degradation of AID*::GFP::MAD-1, the optimal concentration may need to be determined empirically for other targets. To perform a dose-response analysis, we varied the concentration of auxin added to the medium (0, 50, 150, and 300 µM; **Fig. 5A-D; Fig. S3F**), with 300 µM representing an upper limit imposed by compound solubility. To control for effects independent of target degradation, we analyzed GFP::LET-502 and AID*::GFP::LET-502 embryos in parallel and compared embryos that contained or had lost the P*lin-26* array. We observed a dose-dependent increase in the penetrance of embryonic lethality and abnormal larval morphology in embryos expressing AID*::GFP::LET-502 and harboring the P*lin-26* array, but not in the controls (**Fig. 5B**). The length of hatched L1 larvae similarly exhibited a dose-dependent decrease (**Fig. 5C**). High resolution (60X) imaging of embryonic development in AID*::GFP::LET-502-expressing P*lin-26* array-containing embryos in the presence (300 µM) or absence (0 µM) of auxin was consistent with these results. No embryos arrested in the absence of auxin (n=12), while at 300 µM auxin, 50% (6/11) arrested without hatching at stages between comma and 2-fold (**Fig. 5D**, **Movie S4**).

**Figure 5.**
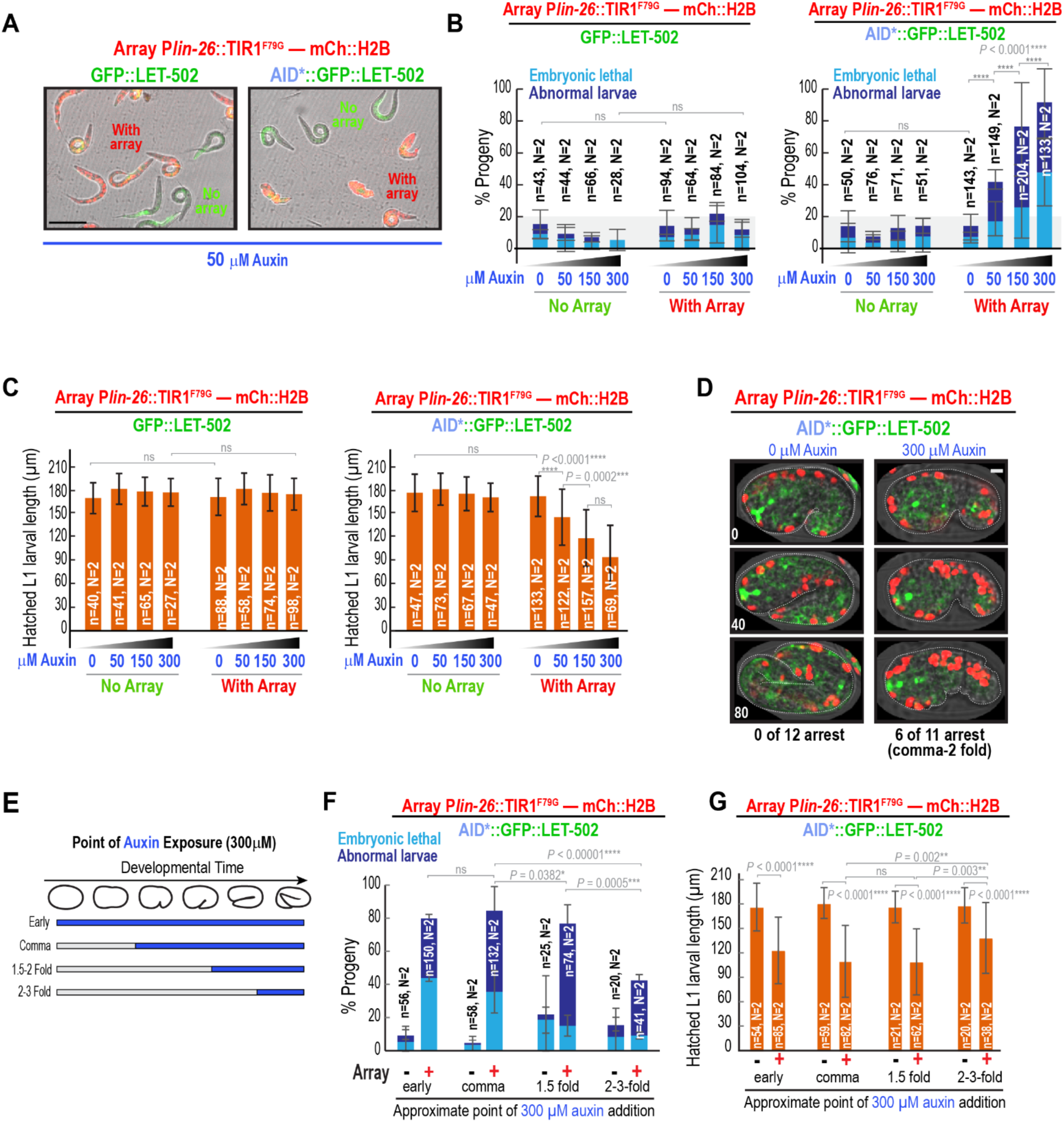
Higher 5-Ph-IAA-AM concentrations increase the penetrance of epidermal LET-502 degradation phenotypes. **(A)** Representative 10X images from whole-well scans of GFP::LET-502 (*left*) and AID*::GFP::LET-502 (*right*) expressing embryos carrying (*red*) or lacking (*green*) the P*lin-26*::TIR1^F79G^—mCh::H2B array after treatment with 50 µM auxin. Progeny that inherit the array have red nuclei (mCh::H2B), whereas progeny lacking the array appear green and serve as an in-well control; this illustrates how the "No array" and "With array" categories in (B, C) were scored. The AID*::GFP::LET-502 image is reproduced from Fig. 4E for comparison. **(B,C)** Graphs show the mean percentage of progeny scored as embryonic-lethal (sky blue) or abnormal larvae (navy blue) (B) and mean hatched L1 larval length (C) for GFP::LET-502 (*left graphs*) or AID*::GFP::LET-502 (*right graphs*) expressing embryos carrying or lacking the P*lin-26*-driven TIR1^F79G^—mCh::H2B operon array, after treatment with a range of auxin concentrations (0, 50, 150, and 300 µM). The shaded band in (B) marks the baseline range of lethality/abnormal larvae observed in this assay. **(D)** Representative merged images (mCh::H2B operon reporter, *red*; AID*::GFP::LET-502, *green*) from 60X confocal time-lapse sequences of AID*::GFP::LET-502-expressing embryos carrying the P*lin-26*::TIR1^F79G^—mCh::H2B array, in the absence (0 µM, left) or presence (300 µM, right) of high-dose auxin (times in minutes relative to the comma stage). The number of filmed embryos that arrested between the comma and 2-fold stages is indicated. (**E**) Scheme of the timed auxin-addition experiment: 300 μM auxin was added to a cohort of staged embryos either immediately after dissection (early) or after they had developed to the comma, 1.5–2-fold, or 2–3-fold stage, to determine when LET-502 is required in the epidermis for elongation. (**F,G**) Graphs show the mean percentage of progeny scored as embryonic-lethal (sky blue) or abnormal larvae (navy blue) (F) and mean hatched L1 larval length (G) for AID*::GFP::LET-502-expressing embryos carrying or lacking the P*lin-26*-driven TIR1^F79G^—mCh::H2B operon array, following the addition of 300 μM auxin at the indicated developmental stages (early, comma, 1.5–2-fold, or 2–3-fold). Error bars for the progeny-outcome graphs (B, F) are SD across independent experiments (weighted by the number of progeny scored per experiment); *P* values, chi-square or Fisher’s exact test. Error bars for the length graphs (C, G), SD of the pooled length measurements; *P* values, Kruskal-Wallis test with Dunn’s multiple-comparisons test. *n* is the number of progeny analyzed; *N* is the number of independent experiments. Scale bars, 5 µm (60X image in D) or 100 µm (10X image in A).

These results suggest that, in addition to TIR1^F79G^ expression, auxin ligand availability impacts degradation efficiency. Notably, increasing the concentration of embryo-permeable auxin in the medium enhanced the phenotypes observed following epidermal LET-502 degradation without off-target effects.

### LET-502 is required in the epidermis for late as well as early elongation

In addition to identifying the tissues where proteins like LET-502 function, the auxin-mediated degradation toolkit also allows investigation of their temporal requirements, which has typically only been possible if a fast-acting temperature-sensitive mutant allele is available. Having established that 300 μM auxin maximizes the penetrance of epidermal LET-502 degradation phenotypes, we used this concentration to determine when epidermal LET-502 is required. For LET-502, prior work with the temperature-sensitive *let-502(sb118ts)* allele monitored embryonic and larval lethality and reported the highest impact when *let-502* was inactivated at the comma stage (∼90% total lethality), a mild effect following inactivation at the 1.5 to 2-fold-stage (∼10% lethality), and no effect after the 2-fold stage (Diogon et al., 2007). However, although viability was assessed, effects on elongation were not directly monitored. It is therefore possible that LET-502 contributes to late elongation, but loss of this late function is insufficient to trigger embryonic lethality.

To determine when the epidermal population of LET-502 functions and whether it contributes to or is dispensable for late elongation, P*lin-26* array embryos with AID*::GFP::LET-502 were allowed to develop to specific stages before auxin was added and developmental outcomes were compared between in-well embryos containing or lacking the array (**Fig. 5E-G).** Auxin addition at the earliest stages of embryogenesis (immediately following dissection) resulted in the expected high rates of embryonic lethality (44%) and abnormal hatched larvae (36%) (**Fig. 5F).** Adding auxin at the comma stage, after the majority of early cell divisions were complete but prior to elongation, resulted in a similarly severe phenotype (36% embryonic lethality and 49% abnormal larvae). Consistent with the *let-502(sb118ts)* data, adding auxin after elongation was already underway at the 1.5-fold or 2-3-fold stages led to levels of embryonic lethality that were similar to array-lacking controls (**Fig. 5F).** However, both of these late addition populations still exhibited elevated levels of abnormal hatched progeny (62% and 33%, respectively), suggesting that lethality cannot be used as a readout for the success of late elongation.

To directly examine elongation, we measured the length of hatched progeny. Under our conditions, 2-fold stage embryos are ∼85-90 µm long after early elongation (prior to twitching) and reach a final average length of ∼175-180 µm after late elongation (**Fig. 5G).** When auxin was added during the early elongation window (comma or 1.5-fold stages), the array-carrying embryos that hatched attained an average length of only ∼109 µm, compared to ∼175-180 µm for the non-array-containing progeny in the same wells (**Fig. 5G),** suggesting failure to elongate significantly beyond the ∼2-fold stage, consistent with a role for LET-502 in early elongation. However, when auxin was added at the 2-3-fold stage, after embryos had already elongated to ∼85-130 µm, the average final length of the hatched larvae was only 138 µm compared to ∼175-180 µm for controls (**Fig. 5G).** Given that these embryos were already 85-130 µm when auxin was added, this result suggests that relatively little elongation occurred beyond the small amount that might be expected during the interval required for LET-502 degradation. Thus, these results suggest that late elongation also depends critically on epidermal LET-502 and highlight the utility of temporal perturbations in a specific tissue in addressing the functions of widely expressed factors at different developmental stages.

## DISCUSSION

Here we describe a toolkit of plasmids, strains, and protocols that enable the tissue-specific degradation of essential proteins during *C. elegans* embryogenesis. Our system is optimized for early embryogenesis and is built around the AID2 system to take advantage of its improved sensitivity and specificity (Hills-Muckey et al., 2022; Negishi et al., 2022; Yesbolatova et al., 2020). We combine the use of TIR1^F79G^ with the eggshell permeable auxin derivative 5-Ph-IAA-AM, which allows rapid target degradation by compound addition throughout embryogenesis (Negishi et al., 2019; Negishi et al., 2022) and provide a set of plasmids that direct expression of a two-gene operon that co-expresses TIR1^F79G^ with an mCh::H2B reporter under a collection of promoters specific for different embryonic tissues. The mCh::H2B reporter allows monitoring of tissue distribution and level of operon expression in analyzed embryos. Our plasmids are compatible with both single-copy insertion and the generation of high-copy extrachromosomal arrays, which enables promoter-dependent tuning of TIR1^F79G^ expression. Because tissue-specific degradation is difficult to quantify when embryos remain fluorescent outside of the target tissue, we also developed a quantitative assay based on the broadly expressed, largely dispensable nuclear envelope marker AID*::GFP::MAD-1 to test the specificity and efficiency of operon-directed degradation. Finally, we establish a workflow that outlines the steps required to maintain rigor and avoid common pitfalls when establishing auxin-dependent degradation systems for new target proteins.

### Titrating TIR1^F79G^ levels and the 5-Ph-IAA-AM concentration to optimize degradation

The *C. elegans* embryo develops from a single cell into a 558-cell larva containing all major somatic tissues within an ∼14-hour window (Corsi et al., 2015; Sulston et al., 1983). This means that new gene expression programs are established in tissues as cell fate is specified and morphogenesis rapidly shapes tissues and elongates the larval body. To effectively counteract these rapid protein expression programs, our results suggest that it is important to titrate TIR1^F79G^ levels so that proteins can be degraded on a comparable timescale. For example, although single-copy insertions of operons driven by the P*eft-3* and P*rps-27* promoters both led to clear pan-tissue degradation of AID*::GFP::MAD-1 and AID*::GFP::LET-502, only the stronger, sustained P*eft-3* promoter degraded AID*::GFP::LET-502 to an extent sufficient to cause an elongation defect (**Fig. 3G,H; Fig. S3E**). Similarly, for the *elt-1* promoter, a single-copy TIR1^F79G^ — mCh::H2B operon insertion did not drive meaningful degradation, whereas a high-copy operon array drove TIR1^F79G^ expression at a level sufficient to degrade epidermal AID*::GFP::LET-502 and trigger an elongation defect (**Fig. 1E,E’; Fig. 4C-E**). We also encountered examples in which TIR1^F79G^ expression had to be titrated down; for example, operon expression driven by the *cnd-1* promoter drove potent tissue-specific degradation as a single-copy transgene but lost tissue specificity in the high-copy format (**Fig. 2A; Fig. S1D**).

In addition to directing high levels of expression, the arrays, which confer a dominant roller phenotype and are not on a specific chromosome, are easy to introduce into even complex genetic backgrounds by backcrossing with males from that background. The arrays we have generated are transmitted between generations at rates between 50 and 90%. With respect to the question of whether array-based expression could lead to mosaic operon expression within the target tissues, we have not noticed significant within tissue mosaicism in the expression of the mCherry histone operon marker for any of the toolkit arrays. This may suggest that transmission to the gametes may be subject to more variability than transmission during the mitotic cell divisions within a tissue.

Our results also suggest that it is important to titrate the concentration of the eggshell permeable 5-Ph-IAA-AM added to the media. Consistent with prior work showing that 50 µM 5-Ph-IAA-AM is effective and well-tolerated (Negishi et al., 2022), we observed clear degradation and elongation phenotypes when embryos expressing the pan-epidermal P*lin-26*::TIR1^F79G^—mCh::H2B operon were treated using this concentration. However, the severity and penetrance of the elongation defects increased as we titrated the 5-Ph-IAA-AM concentration from 50 µM up to 300 µM (**Fig. 5B,C**), which we found to be close to its solubility limit. In our experiments, these higher concentrations were also well-tolerated, and phenotypes were not observed in treated control embryos expressing non-degradable GFP::LET-502 or lacking the pan-epidermal TIR1^F79G^ operon (**Fig. 5B,C**).

### Tissue- and stage-specific degradation reveals where and when LET-502 is required for elongation

As a test case, we used the toolkit to dissect the role of the Rho kinase LET-502 in embryo elongation, where it promotes actomyosin contractility by phosphorylating the myosin regulatory light chain MLC-4 and antagonizing the MEL-11 phosphatase (Gally et al., 2009; Piekny et al., 2000; Wissmann et al., 1999; Wissmann et al., 1997). Although LET-502 is broadly expressed (**Fig. 3D**; (Piekny et al., 2003)), degradation in the intestine, pharynx, muscle, neuroblasts, or sensory neurons had no effect on viability or larval morphology. By contrast, degradation in the epidermis using either the *elt-1* or *lin-26* promoters reproduced the elongation defects observed following pan-tissue degradation or RNAi-mediated depletion. This provides strong confirmation for the idea that LET-502 in the epidermis is required for elongation, which prior work had inferred from the ability of LET-502 expressed under a promoter active in epidermal seam cells to partially rescue a *let-502* mutant (Drewnik et al., 2021).

The toolkit also allowed us to refine the timing of the LET-502 requirement. Embryo elongation is proposed to occur in two phases, an early phase prior to the 2-fold stage, which is thought to be driven by actomyosin contractility within the seam cells of the epidermis, and a late phase beyond the 2-fold stage, when force is generated by the body wall muscle that underlies the epidermis (Carvalho and Broday, 2020; Vuong-Brender et al., 2016). Time-shift experiments with a thermosensitive *let-502* allele have been interpreted to indicate that LET-502 is required only during the early phase (Diogon et al., 2007; Vuong-Brender et al., 2016). However, those experiments scored viability rather than directly monitoring elongation. By adding auxin at successively later stages and measuring the length of hatched larvae, we found that degrading epidermal LET-502 after the 2-fold stage caused little lethality, in agreement with the earlier work, but nevertheless produced larvae that had lengthened only minimally relative to their length when auxin was added. These results suggest that epidermal LET-502 is required for late as well as early elongation, and that embryonic lethality is not a reliable readout for successful late elongation. Whether LET-502 contributes contractile force during late elongation or has another function such as maintaining the circumferential actin bundles that are remodeled by muscle-generated force (Lardennois et al., 2019), is an important question for future work.

Overall, our results highlight the ability of tissue-specific and temporally controlled degradation to facilitate detailed mechanistic analysis. The tools and workflow we present here also enable a rigorous approach to initiate these studies. The modular design of our plasmids also makes it straightforward to expand the promoter set as needed to extend this strategy to the many broadly expressed essential genes whose spatial and temporal requirements during embryogenesis remain to be defined.

## MATERIALS AND METHODS

### Plasmid construction

All plasmids directing expression of the TIR1^F79G^—mCh::H2B operon under different tissue-specific promoters (listed in **Table S1**) were constructed using Gibson Assembly (Gibson et al., 2009). First, the plasmid pLA024 was constructed by PCR amplifying the TIR1^F79G^ fragment from pTN4 (addgene no. #174212) using a reverse oligo that added a 3xFlag onto the TIR1^F79G^ C-terminus and cloning this into the plasmid framework amplified from pOD2494/pPLG187 (Lara-Gonzalez et al., 2019), which replaced the sequence encoding GFP::PH with the TIR1^F79G^ fragment, placing it downstream of the *mex-5* promoter and upstream of the operon linker::mCherry::his-11::tbb-2 3’UTR cassette. For all of the TIR1^F79G^ operon-containing plasmids except the one driven by the *mex-5* germline promoter, Gibson assembly was used to combine a tissue-specific promoter fragment (see **Table S1** for full sequences), the TIR1^F79G^::3xFlag::operon linker:: mCherry::his-11::tbb-2 3’UTR cassette amplified from pLA024 and the MosSCI targeting vector backbone amplified from pCFJ151 (Frokjaer-Jensen et al., 2008). For the germline-expressed P*mex-5*::TIR1^F79G^::3xFlag construct, fragments containing the *mex-5* promoter and TIR1^F79G^ with its C-terminal FLAG tag were PCR amplified from pLA024 and the MosSCI targeting vector backbone was amplified from pCFJ151. For Gibson assembly, PCR primers were designed to provide at least 20 bp of overlap between adjacent fragments. Equimolar amounts of DNA fragments (5 μl total) were combined with 15 μl of 1.33X homemade Gibson Assembly mix [133 mM Tris-HCl (pH 7.5), 267 μM each dNTP, 1.33 mM NAD, 6.67% PEG 8000, 13.3 mM MgCl₂, 13.3 mM DTT, 5.33 U/ml T5 exonuclease, 33.3 U/ml Phusion High-Fidelity DNA polymerase, and 5330 U/ml Taq DNA ligase] and incubated at 50°C for 1 h. Following assembly, 5–10 μl of the reaction was used for bacterial transformation and positive clones were screened. For the P*lin-26*::TIR1 ^F79G^::mCh::H2B construct, the *lin-26* promoter along with a partial *myo-2* promoter fragment was amplified from OD2018/pSW339 (see **Table S1)** before combining with other individual fragments as described above. All plasmids were midi-prepped using the QIAGEN Plasmid Plus Midi Kit (Qiagen cat. no. 12945) with one additional 0.7 ml Buffer ETR wash (two total) to ensure adequate endotoxin removal before preparing plasmid mixes for injection into worms. All the TIR1^F79G^ plasmid constructs along with complete sequences and maps will be deposited in Addgene by K.O.

### C. elegans strains

#### Single copy insertion strains

*C. elegans* strains (listed in **Table S2**) were maintained at 16-22.5°C on nematode growth medium (NGM) plates seeded with a lawn of OP50 *E. coli* (Brenner, 1974). All single-copy transgenes, except for the single-copy-insertions that express P*eft-3*::TIR1^F79G^, were generated using the MosSCI transposon-based method (Frokjaer-Jensen et al., 2012; Frokjaer-Jensen et al., 2008; Frokjaer-Jensen et al., 2014) to insert them at defined chromosomal sites. Transgenes with TIR1^F79G^ — mCh::H2B expressed under tissue-specific promoters were cloned into pCFJ151 (as described above), which contains the *Cb-unc-119* selection marker and appropriate homology arms. Injection mixes contained the pCFJ151-derived repair plasmid (50-100 ng/μL), transposase plasmid pCFJ601 (Peft-3::Mos1 transposase, 50 ng/µl), and four plasmids for negative selection against chromosomal arrays: pMA122 (P*hsp-16.41::peel-1*, 10 ng/μL), pCFJ90 (P*myo-2*::mCherry, 2.5 ng/μL), pCFJ104 (P*myo-3*::mCherry, 5 ng/μL), and pGH8 (P*rab-3*::mCherry, 10 ng/μL). Mixes were injected into the strain EG6429 for insertion into the *ttTi5605* site on Chr. II or into EG8078 for insertion into the universal *oxTi185* site on Chr. I. After 9-10 days at RT, when the worms had starved out and many small third generation progeny were on the plate, the plates were heat-shocked by placing the box of plates into a 34°C incubator for 3.25 hours to induce the expression of PEEL-1 to kill worms containing extra chromosomal arrays. Several hours after removing the plates from the incubator a quarter-sized chunk was cut out of each plate and flipped upside down onto a new plate. The next day, plates with young moving worms without fluorescent markers were identified by screening on a fluorescence dissecting scope, and individual non-fluorescent worms were singled onto small plates. ∼4 days later these plates were screened to identify plates lacking any Unc worms, suggesting that the parent was homozygous for the insertion. Homozygous insertion was confirmed using PCR that spanned both homology regions and construct expression was confirmed by imaging.

Single-copy insertion of the P*eft-3*:: TIR1^F79G^ transgene was performed using a MosSCI histamine-based negative selection strategy (El Mouridi et al., 2021). MosSCI strains EG6429 (*ttTi5605*, Chr. II) and EG8078 (*oxTi185*, Chr. I) were used for integration. A 2X injection mix containing pSEM233 (P*mlc-1*::TagRFP-T) or pSEM231 (P*mlc-1*::GFP) as a fluorescent co-injection marker (20 ng/µL), pCFJ1532 (P*smu-1*::Mos1 transposase; 40 ng/µL), pSEM238 (P*snt-1*::ce-HisCl1; 40 ng/µL), and stuffer DNA (50 ng/µL) was prepared. 5 µL of the 2X mix was combined with the plasmid pOD5558 (see **Table S1**) (final concentration 25 ng/µL) and nuclease-free water to a final volume of 10 µL and injected into the gonads of 10–15 young adult hermaphrodites. Injected worms were singled onto medium NGM plates and incubated at 23°C. Plates were screened 2–3 days post-injection for viable progeny and incubated for an additional 7 days or until starvation. For negative selection, histamine dihydrochloride was prepared as a 500 mM solution in water and 500 µL was added directly to each plate. After 10 min incubation at room temperature, animals expressing the histamine-gated chloride channel (ce-HisCl1) were paralyzed. Histamine-resistant, non-fluorescent moving larvae were isolated using a fluorescence stereomicroscope and singled onto NGM plates. Correct single-copy integration was confirmed in the progeny by PCR and expression was verified by imaging.

#### Extrachromosomal array strains

Extrachromosomal arrays were generated by preparing mixes combining an operon-containing plasmid marked with *Cb-unc-119* and pRF4, which encodes the dominant roller marker *rol-6(su1006)* (Mello et al., 1991), at equal concentrations (50 ng/µL each). Injection mixes were injected into the MosSCI strain EG6429 (*ttTi5605 II*; *unc-119(ed3)*), which served as a convenient source of *unc-119* mutant animals.

In this background only worms carrying extrachromosomal arrays containing both plasmids exhibited a moving roller phenotype. Approximately 40 adult hermaphrodites were injected and plated (two worms per medium NGM plate). Plates were maintained at room temperature and were allowed to starve over ∼9 days to enrich for third-generation rollers. A quarter-sized agar chunk was cut out of the center of each starved plate and the worms on the chunk were smeared across a new medium-sized plate using the scalpel to discard the chunk after transferring the worms. Plates were incubated at RT for 2 days to allow the small third generation progeny to reach a larger size and 3 roller worms from each plate (60 total) were singled onto individual small plates. After three days, the plates were screened to identify the plates with roller arrays that transmitted at the highest frequency (50-90%). Embryos from the lines with the most robust inheritance were screened for mCherry::H2B expression on the fluorescence confocal microscope, and best lines were selected for mating to degron-tagged AID*::GFP::MAD-1 to test target degradation in the presence of auxin.

#### CRISPR strains

Endogenous *let-502* was tagged with either GFP or AID*::GFP at the N-terminus using CRISPR/Cas9 ribonucleoprotein (RNP)-mediated genome editing, as previously described (Paix et al., 2017). Briefly, repair templates were generated by using PCR to amplify sequences encoding GFP (ttctacgatctcacggacgaaaatcgagctgcagccATGAGTAAAGGAGAAGAACTTTTCA, atccaccagctgatcacgcagctcatcctgctccaTTTTGTATAGTTCATCCATGCCA) or AID*::GFP (ttctacgatctcacggacgaaaatcgagctgcagccATGCCTAAAGATCCAGCCAAA, atccaccagctgatcacgcagctcatcctgctccaTTTTGTATAGTTCATCCATGCCA) from the plasmid pLA023. PCR products were purified using the MinElute PCR Purification Kit (Qiagen, Cat. No. 28004), and at least 3 µg of purified template was dried under vacuum (SpeedVac) per injection mix. A target site near the *let-502* N-terminus was identified using CHOPCHOP (https://chopchop.cbu.uib.no/). Purified Cas9 protein was obtained from the MacroLab (UC Berkeley) and crRNAs and tracrRNA were purchased from IDT and resuspended in IDT duplex buffer to a final concentration of 200 µM each. A co-CRISPR strategy targeting dpy-10 was employed to facilitate the identification of edited animals. A guide RNA mix was prepared by combining 2.5 µL tracrRNA, 0.5 µL *dpy-10* crRNA (5′-GCTACCATAGGCACCACGAG-3′), and 2.0 µL *let-502* crRNA (5′-CAGAGACTATTGATAGATTG-3′), followed by annealing (95 °C for 5 min, then 22 °C for 5 min in a thermocycler). The Cas9-RNP complex was assembled by combining 2 µL of the annealed guide RNA mix with 5 µL purified Cas9 protein (40 µM) and incubating at room temperature (∼22 °C) for 5 min. To prevent Cas9 precipitation, 0.7 µL of 3 M KCl was added and mixed thoroughly. The RNP mix (3 µL) was then used to resuspend the dried repair template DNA to generate the final injection mixes. Injection mixes were centrifuged at maximum speed for 15–30 min at 4 °C to remove particulates and each mix was injected into both germlines of approximately 30 adult N2 hermaphrodites and the injected worms were plated 6 worms/plate. Worms were incubated at RT for 6 hours to allow them to lay embryos produced prior to injection with the CRISPR mix before transferring them to a new set of plates. Worms were allowed to lay embryos for 24 hours to retrieve the majority of the embryos with successful CRISPR edits before removing them from the plates. Plates were kept at RT for about 3 days, until the laid embryos were adult worms. Approximately 120 dpy or roller progeny, indicating successful homozygous or heterozygous *dpy-10* co-CRISPR, were singled on tiny plates and allowed to lay embryos for 24-48 hours before subjecting the parent worm to single worm genotyping PCR across the insertion site followed by Sanger sequencing to confirm correct, in-frame insertion of GFP or AID*::GFP. Homozygous or heterozygous edited strains were outcrossed once to the N2 parental strain to remove the *dpy-10* mutated allele.

To generate the strain expressing AID*::GFP::MAD-1 (PLG28), a sequence encoding AID*::GFP was inserted at the N-terminus of the endogenous *mad-1* locus by using the self-excising cassette method as described (Dickinson et al., 2015). A Cas9-sgRNA plasmid (pPLG118) containing the sgRNA sequence (tgattgcattaaacatatt; (Wang et al., 2017)) was used together with a plasmid (pLA023) containing the AID*::GFP SEC repair template.

#### Generation of auxin-mediated target degradation strains

Strains for auxin-mediated target degradation were generated by generating males from the tagged target (GFP::LET-502, AID*::GFP::LET-502 and AID*::GFP::MAD-1) insertion strains and mating these males with single-copy insertion or array strains expressing the TIR1^F79G^—mCh::H2B operon under control of different tissue-specific promoters. For single-copy operon insertion strains, F2 progeny from these matings were screened to identify strains homozygous for the operon insertion and the GFP or AID::GFP-tagged target gene. For the array strains, roller hermaphrodites carrying the TIR1^F79G^—mCh::H2B operon were mated with males from the target (GFP::LET-502, AID*::GFP::LET-502, and AID*::GFP::MAD-1) insertion strains and the mated hermaphrodites were singled. L4 hermaphrodite roller progeny heterozygous for the target gene insertions were picked from plates that had 50% male progeny and were crossed a second time with target gene insertion males and singled following mating. L4 hermaphrodite roller progeny from this second mating were singled from plates with 50% males and were allowed to lay embryos for 24-28 hours before PCR screening of the parent worms to identify worms homozygous for the target gene insertion (∼50% of the singled worms). Roller worms homozygous for the target gene insertion are maintained by passing rollers at each generation.

### RNA-mediated interference (RNAi)

Templates to generate double stranded RNAs (dsRNAs) targeting *let-502* (taatacgactcactataggGCATGGCGCTGATTCGGA, aattaaccctcactaaagGTCGATTCCTGCTCAGACGA) and the GFP tag (aattaaccctcactaaagGCCAACACTTGTCACTACTTTCTGTTATGG, taatacgactcactataggGTATAGTTCATCCATGCCATGTGTAATCCC) were generated by using the primers in parentheses to amplify the corresponding sequences from N2 genomic DNA and the plasmid pNV4, respectively. DNA templates were gel-purified using a QIAquick PCR Purification Kit (Qiagen). Single stranded RNA (ssRNAs) was generated from DNA templates using MEGAscript T3 and T7 Transcription Kits (Invitrogen) and purified using a MEGAclear Transcription Clean-up Kit (Invitrogen). Equimolar amounts of T3- and T7-generated ssRNA were combined with 3X soaking buffer (32.7 mM Na_2_HPO_4_, 16.5 mM KH_2_PO_4_, 6.3 mM NaCl, and 14.1 mM NH_4_Cl) and annealed by heating at 68°C for 10 min, followed by 37°C for 30 min to form dsRNA. GFP dsRNA and *let-502* dsRNA were injected into L4 animals (OD6238) at concentrations of 1.45 mg/mL and 1.2 mg/mL respectively. For single RNAi experiments, animals were allowed to recover on small plates at 20°C for 20-22 hours before dissecting the gravid adults and imaging the embryos. A more stringent, double RNAi injection protocol was also employed. In double RNAi experiments, L4 worms were injected with dsRNA and allowed to recover for 22 hours at 20 °C before injecting the worms a second time and incubating at 16°C for 20-22 hours prior to dissection and imaging. Embryos were dissected from gravid adults and imaged; embryonic lethality was scored as described below.

### Imaging

For the majority of experiments, imaging of *C. elegans* embryonic development was performed as previously described (Green et al., 2024; Khaliullin et al., 2019; Wang et al., 2019). Briefly, 10-20 gravid adults (selecting only the array-possessing roller worms for array strains) were dissected in ice-cold M9 (42 mM Na_2_HPO_4_, 22 mM KH_2_PO_4_, 86 mM NaCl, and 1mM MgSO_4_) with 0.1 mg/ml of the anesthetic tetramisole hydrochloride (TMHC; Sigma) and with or without the indicated concentrations (between 50 and 300 μM) of the eggshell permeable auxin 5-Ph-IAA-AM diluted from a 50mM stock dissolved in DMSO (BioAcademia; Product code: 30-004). Young (2 to 32-cell stage) embryos from each test condition were transferred by mouth pipet into an individual well of a glass bottom 384-well Sensoplate (Greiner Bio-One) containing 70 μl of the same solution. During dissection and setup, the solution and plate were covered in foil to avoid light exposure. For most experiments, the plate was maintained on ice to slow development and maintain synchrony across conditions until dissections for all conditions were completed. The loaded plate was allowed to settle for 10 minutes or spun for 1 minute at 600 x g to seat the embryos. Embryos were imaged on a CV1000 (Yokogawa Electric Corporation) spinning disk confocal system (bright-field: 90% power, 25 ms, 20% gain and 488nm: 100% power, 200 ms, 60% gain, 568nm: 45% power, 150 ms, 60% gain) with a 512x512 EM-CCD camera (Hamamatsu), in a 16°C temperature-controlled room, which maintained the embryos at ∼22°C on the microscope stage. A 60X 1.35NA U-PlanApo objective was used to acquire 18 x 2 μm z-stacks of 5 fields per well, each field containing 1-5 embryos, at 20-minute intervals for 10.5 hours. To assess embryonic lethality, wells were post-scanned 20-22 h after the start of imaging using bright-field optics and a 10 x 0.4 NA U-PlanApo objective, collecting 18 x 2 μm z-stacks of brightfield and a central plane fluorescent image to determine presence or absence of the mCherry marked array. All image acquisition was performed using CellVoyager software (Yokogawa).

For experiments varying the timepoint when auxin was added, embryos were sequentially staged by dissecting 10-20 gravid adults expressing AID*::GFP::LET-502 and carrying the P*lin-26*::TIR1^F79G^*—* mCh::H2B operon expressing array (OD6343), every hour for 6 consecutive hours. At each timepoint, 2-16 cell stage embryos, were transferred into a well in a 384-well Sensoplate (Greiner Bio-One) containing the anesthetic solution described above, lacking auxin, and held at room temperature. After the last dissection, 5-Ph-IAA-AM was added to all of the wells to a final concentration of 300µM. To assess the effects of auxin concentration on embryonic lethality, gravid adults were dissected into anesthetic solution containing 0, 50, 150 or 300 μM of 5-Ph-IAA-AM at room temperature. For experiments varying auxin concentration or time of addition, the plate was transferred to the CV1000 (Yokogawa Electric Corporation) spinning disk confocal system after auxin addition to image the wells at 10X resolution to determine the initial staging of embryos. The plate was left overnight in the dark (∼22°C) without further 60X imaging, to prevent phototoxicity effects, until the next day (20-22 hours after the last dissection) when the wells were rescanned at 10X and embryonic lethality and abnormal L1 progeny were scored (described below). In addition to comparison between wells, the development of array-containing embryos (red and green markers present) was compared to that of embryos lacking the array (green marker only) in the same well which served as an internal control.

### Image Analysis

All imaging data was analyzed and processed using FIJI/ImageJ (National Institutes of Health). For each experiment filming embryonic development, compared conditions were stage-matched (using comma as a benchmark) and scaled equivalently. Stacks from each compared condition were combined and montages of relevant image frames were generated from the combined stacks into a single montaged tif for direct comparison. Photoshop (Adobe) was used to crop out and generate publication ready images. To ensure comparable scaling for all image processing steps, the combined montage image output from FIJI, containing all conditions to compare, was used for any additional image scaling and processing steps in Photoshop.

### Quantification of degradation in AID*::GFP::MAD-1 strains

Degradation of AID*::GFP::MAD-1 in targeted tissues was monitored by analyzing GFP signal in the target tissue region at each time point, using a custom macro built in Fiji (Schindelin et al., 2012). To do this, a single central focal plane, containing the majority of the target tissue, was chosen for analysis over time. A binary mask was generated based on the red (mCherry-H2B) nuclear signal, which serves as a readout for TIR1^F79G^ expressing cells. Thresholded binary mask edges were dilated two times so that the mask enclosed the AID*::GFP::MAD-1 nuclear envelope signal surrounding each nucleus, the expanded nuclear mask was applied to the AID*::GFP::MAD-1 (green) channel, and the GFP signals in individual nuclei were identified using the Analyze Particles tool. For each timepoint, the GFP intensity data was normalized to the area of the red masked region. Total signal area was chosen for normalization because the intensity of mCherry signal can vary across the set of imaged embryos, due to embryo orientation, changes in the number of nuclei from frame to frame, and variation in the expression from the extrachromosomal arrays in different embryos. The data acquired from the early timepoints, when red signal was not yet expressed, was cut off to eliminate noise in the dataset. The first time point when the signal was detectable by the thresholded mask was used to help define the time window of quantification presented in Figures 1, 2, and to estimate the timepoints when the designated promoter began to effectively express TIR1^F79G^. To generate plots, embryos for each condition were processed to quantify GFP intensity over time and the timepoint most closely matching “comma” stage was noted. Some imaged embryos were not quantified due to suboptimal orientation or positional/focal plane shifts during the overnight imaging timecourse, however a minimum of 5 embryos were quantified for each tested condition. Data sets were time aligned according to the comma benchmark; corresponding frames from onset of particle detection onward were averaged across all datasets to generate average GFP values for each time point. The first 10 timepoints were averaged for controls and degron test conditions, and the control average was used to normalize each point in the timelapse series and the average bar plots.

### Quantification of L1 larval length

To analyze possible elongation defects in hatched larva (**Figures 4E, 5C, and 5G**), progeny length was directly measured tip-to-tail using FIJI freehand line tool for every hatched worm in the 10X post-scan dataset. For each image, length of array containing (red co-marker expressing) worms and worms lacking the array (green only) were measured separately, and the results were plotted accordingly. Worms were counted as “abnormal” if their length values fell below 2 standard deviations for the controls. In some instances, the hatched worms were so short they were difficult to distinguish from arrested embryos. In this case, the presence of an intact eggshell was used to differentiate unhatched progeny from short, hatched worms.

### Automated cropping of embryos

Overnight 60X imaging data contains multiple embryos in each field. To analyze embryos individually, each data set was processed using the automated cropping software previously developed by our group (Khaliullin et al., 2019; Wang et al., 2019). This cropping algorithm detects individual embryos by generating a binary mask from the bright-field images, sequentially cropping each embryo, orienting it, and saving the embryo in a separate file. Before cropping, the software corrects for drift, subtracts background, and performs depth attenuation correction in each fluorescence channel. This software is freely accessible zenodo.org/record/1475442#.W9jvApNKiUl and the source code can be downloaded from github.com/renatkh/embryo_crop.git.

### EMB lethality and abnormal larva scoring

Embryonic lethality was scored based on a 10X scan of the entire well, approximately 20-22 hours after the start of imaging. For initial experiments using 50μM auxin, all progeny were scored as either hatched normal, hatched abnormal, or unhatched (embryonic lethal) based on visual assessment of 10X well scans. Embryos arrested at or before the 2-cell stage were excluded, due to possible osmotic issues prior to eggshell formation at the time of dissection. For array experiments, embryos or worms containing arrays (red co-marker expressing) were counted separately from embryos and worms in the same well that were lacking the array (green only). Plots of embryonic lethality for array containing strains show the data from array containing progeny only. In experiments directly analyzing possible elongation defects, the length of hatched L1 worms were directly measured (see Image analysis) and worms with hatched lengths that were less than two standard deviations from the control mean were deemed to be abnormal.

Because the sample sizes for each experiment vary between experiments, the averages and error bars presented in lethality plots represent weighted averages and weighted standard deviations, which were calculated using standard formulas. This was done to prevent experiments with smaller sample sizes from exerting undue effects on the overall average embryonic lethality values across multiple experiments. For the L1 length data, the individual measurements from independent experiments were pooled and the standard deviation in length was shown for the population.

### Statistical Analysis

Statistical analysis was performed using Excel or GraphPad Prism. Percentages of progeny scored as embryonic lethal or as abnormal larvae were compared by chi-square test on the pooled counts of progeny in each outcome category, with Fisher’s exact test substituted when any expected categorical count was less than 5. Hatched L1 lengths were compared by Kruskal-Wallis test with Dunn’s multiple-comparisons test. Normalized per-embryo AID*::GFP::MAD-1 intensities, averaged across the 10-timepoint window indicated in each panel, were compared between +auxin and −auxin conditions by two-tailed Welch’s t-test. For progeny outcome and larval length comparisons, tests were performed on measurements pooled across independent experiments. The test used is indicated in each figure legend.

## ACKNOWLEDGEMENTS

We thank Pablo Lara-Gonzalez for the TIR1^F79G^-containing plasmid and helpful discussions, the Jin lab for pRF4, and Noura Yassir for initial piloting efforts. The authors used Claude (Anthropic) for editorial assistance, including revising text for length and clarity, copy-editing, and checking internal consistency and references. All text was reviewed and edited by the authors, who take full responsibility for the content of this manuscript.

## AUTHOR CONTRIBUTIONS

Conceptualization: NV, RG, AD, KO; Methodology: NV, RG, AD, KO; Validation: NV, RG; Formal analysis: RG; Investigation: NV, RG, AOD, SM, DG, KO; Resources: NV, RG, KO, SM, AOD, DG; Data curation: NV, RG; Writing – original draft: NV, RG, AD, KO; Writing – review & editing: NV, RG, AD, KO; Visualization: NV, RG, KO; Supervision: AD, KO; Project administration: AD, KO; Funding acquisition: AD, KO.

## COMPETING INTERESTS

The authors declare no competing or financial interests.

## FUNDING

This work was supported by the funding agencies listed below.

National Institutes of Health grant R01GM147265 (KO)

National Institutes of Health grant R35GM163658 (KO)

National Institutes of Health grant R01GM074215 (AD)

National Institutes of Health grant R35GM161808 (AD)

## DATA AND RESOURCE AVAILABILITY

### Strain availability

The recommended toolkit strains containing the TIR1^F79G^—mCh::H2B operon arrays and single-copy insertions will be made available through the Caenorhabditis Genetics Center (CGC). All other strains will be available upon request.

### Plasmid availability

Plasmids driving tissue-specific expression of the TIR1^F79G^—mCh::H2B operon will be made available from Addgene.

## SUPPLEMENTARY FIGURES AND FIGURE LEGENDS

**Figure S1.**
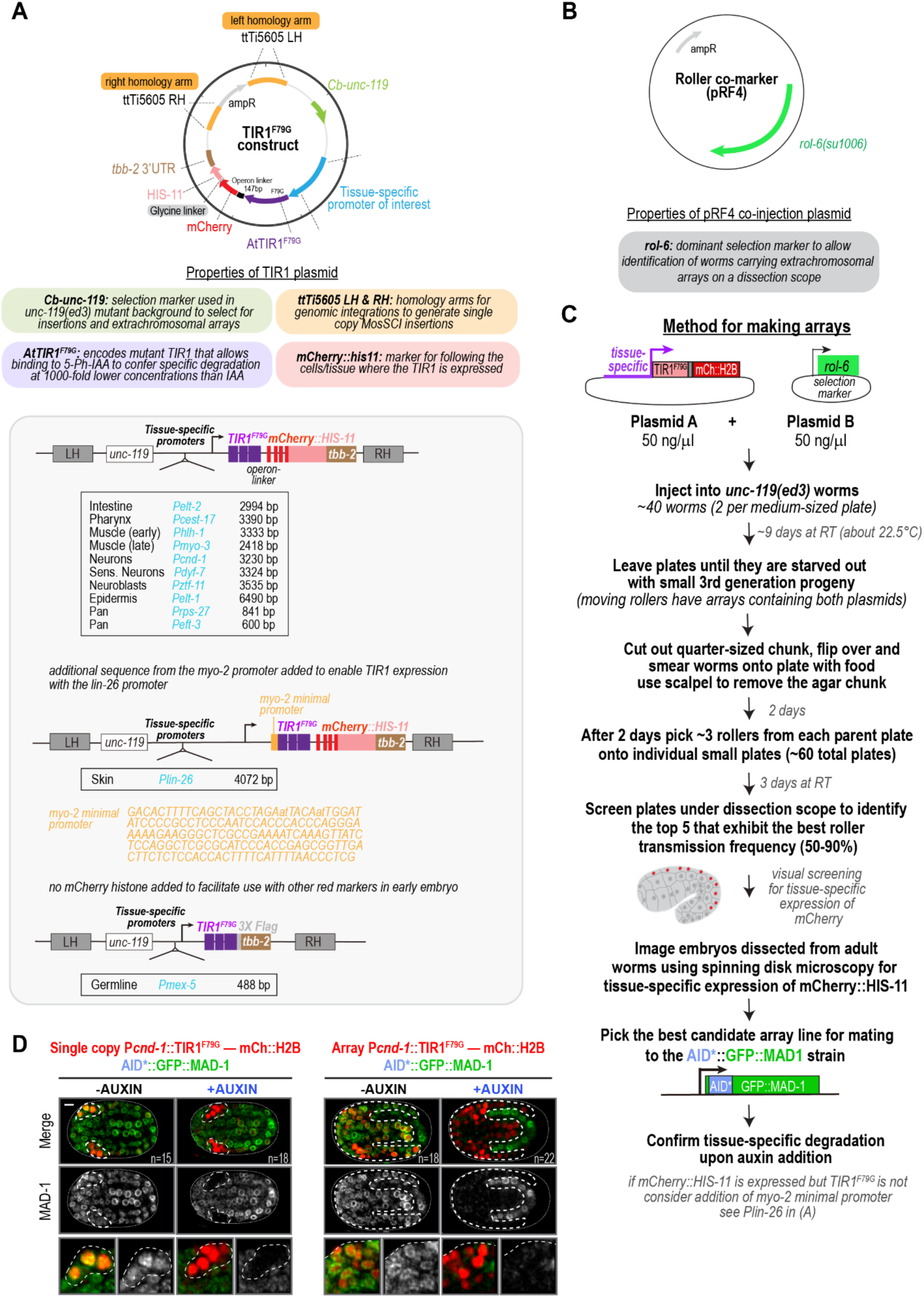
Plasmid design, array construction, and comparison of single-copy versus array expression. **(A)** Design of the TIR1^F79G^—mCh::H2B operon expression construct. (*top*) Circular map of the TIR1^F79G^—mCh::H2B operon plasmid, showing the *Cb-unc-119* selection marker (used in the *unc-119(ed3)* background to select insertions and arrays), the ttTi5605 left and right homology arms (LH, RH) for single-copy MosSCI integration, the tissue-specific promoter of interest, AtTIR1^F79G^ (the mutant TIR1 that binds 5-Ph-IAA derivatives and confers degradation at ∼1000-fold lower concentrations compared to IAA), and the mCherry::HIS-11 (mCh::H2B) reporter that marks nuclei in the tissue of where the TIR1^F79G^ is expressed; element properties are summarized below the map. (*middle*) The linear operon structure with sequences corresponding to the tissue-specific promoter— TIR1^F79G^—the operon linker— mCherry::HIS-11—tbb-2 3’ UTR is shown with a table showing the gene names and sizes of the promoter fragments used for expression in the indicated tissues. For the *lin-26* (epidermis) promoter, additional sequence from the *myo-2* minimal promoter as shown was added to enable TIR1^F79G^ expression. **(B)** Map of the pRF4 co-injection plasmid, which carries an ampicillin-resistance marker (ampR) and the dominant *rol-6(su1006)* marker used to identify worms carrying extrachromosomal arrays on a dissecting scope. **(C)** Protocol for generating extrachromosomal-array strains. The TIR1^F79G^—mCh::H2B operon plasmid (Plasmid A) and the pRF4/*rol-6* selection plasmid (Plasmid B) are co-injected (50 ng/µl each) into *unc-119(ed3)* worms (40 total, two per plate). Plates are grown ∼9 days until they starve out with third-generation progeny; a large quarter-sized chunk is cut out from each plate, and the worms are smeared onto food on a new plate. After 2 days, ∼3 rollers per parent plate (∼60 total) are singled onto individual tiny plates. After 3 days, embryos from the lines with the highest roller transmission (50-90%) are screened by spinning-disk microscopy for tissue-specific mCh::H2B expression, and the best line is mated to the AID*::GFP::MAD1 strain to confirm tissue-specific degradation upon auxin addition. If mCh::H2B is expressed but TIR1^F79G^ is not, adding the *myo-2* minimal promoter fragment upstream of the TIR1 coding region may enable TIR1^F79G^ expression (see the P*lin-26* construct in A). (**D**) Comparison of AID*::GFP::MAD-1 degradation driven by a single-copy insertion of the *Pcnd-1*::TIR1^F79G^::mCh::H2B operon (*left*) and a *Pcnd-1*::TIR1^F79G^::mCh::H2B operon containing array (*right*). Embryos from each strain are shown in the absence (−AUXIN, *left*) and presence (+AUXIN, *right*) of 50 µM auxin (5-Ph-IAA-AM). The top row shows merged images with AID*::GFP::MAD-1 in green and mCh::H2B in red. Dashed lines outline the mCh::H2B expressing cells. The middle row shows AID*::GFP::MAD-1 alone in grayscale and zoomed panels on the bottom show a higher-magnification view (∼2.5X) of part of the highlighted region from the images above. The single-copy insertion drives neuron-specific degradation, whereas the array drives broad, non-specific degradation across the embryo, which is why the single-copy *Pcnd-1* insertion is recommended for degradation experiments and the array is not (see Fig. 2B). *n* values in the lower right corner of the merged panels are the number of embryos imaged. Scale bar, 5 µm.

**Figure S2.**
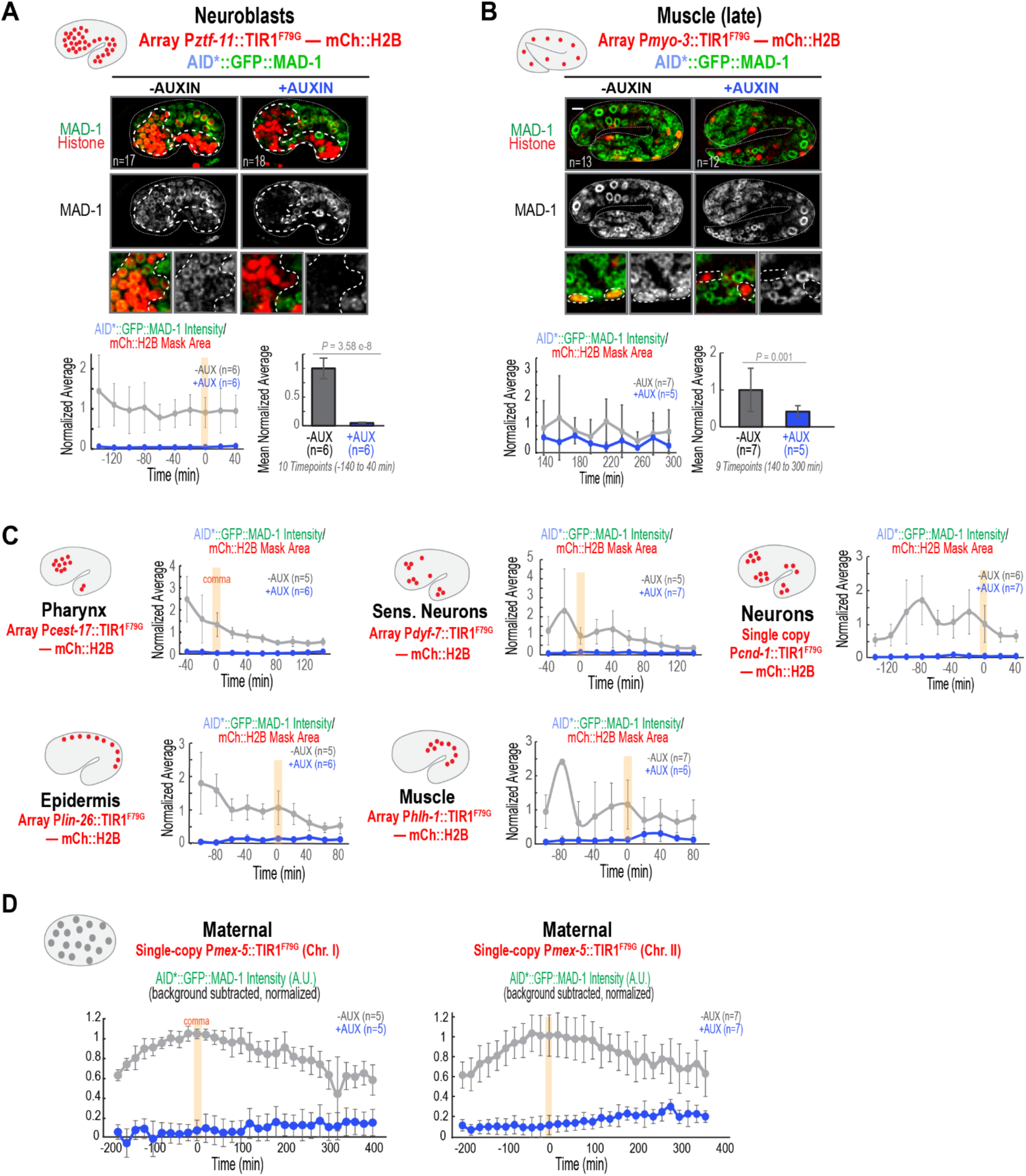
Additional characterization of the tissue-specific auxin-inducible degradation toolkit strains. (A,B) Auxin-mediated degradation of AID*::GFP::MAD-1 in neuroblasts using the P*ztf-11-driven operon* array (A) or in late muscle using the P*myo-3*-driven operon array (B), shown in the absence (*left*) or presence (*right*) of 50 µM auxin (5-Ph-IAA-AM). For each, an embryo schematic indicates the positions of the operon expressing cells. (*top row*) Images show the merge with AID*::GFP::MAD-1 in green and mCh::H2B in red. (*middle row*) Images show AID*::GFP::MAD-1 in grayscale. White dashed lines outline the operon-expressing tissue, defined by the nuclear mCh::H2B reporter. (*bottom row*) Zoomed panels show a higher-magnification view (∼2.5X) of regions from the images above. (*bottom, left*) Graph plots normalized AID*::GFP::MAD-1 intensity (AID*::GFP::MAD-1 intensity / mCh::H2B mask area) versus time showing the mean ± SD at each timepoint relative to the comma stage (timepoint 0; gray, −auxin; blue, +auxin). (*bottom, right*) Bar graph (*right*) showing the mean ± SD of the per-embryo averages across the indicated 9 or 10-timepoint windows; quantified as described for Fig. 1D′, E′). *P* values are two-tailed Welch’s t-tests comparing the + and − auxin conditions. **(C)** Graphs plot normalized AID*::GFP::MAD-1 intensity (AID*::GFP::MAD-1 intensity / mCh::H2B mask area) versus time showing the mean ± SD at each timepoint relative to the comma stage (timepoint 0; gray, −auxin; blue, +auxin) for operon-expressing embryos from the indicated strains. Data was quantified as in *Fig. 1D’,E’* and corresponds to the summary graphs shown in Fig. 2A. **(D)** Quantification of AID*::GFP::MAD-1 degradation for the two maternally expressed single-copy P*mex-5* insertions (Chr. I, *left*; Chr. II, *right*), in the absence or presence of 50 µM auxin (5-Ph-IAA-AM). These strains lack the mCh::H2B operon reporter; an embryo schematic summarizes the location of the expressing cells. Plots show the mean ± SD of the background-subtracted, normalized total AID*::GFP::MAD-1 intensity at each timepoint relative to the comma stage (comma = timepoint 0; gray, −auxin; blue, +auxin); see also Fig. 2A. *n* is the number of embryos analyzed. Scale bars, 5 µm.

**Figure S3.**
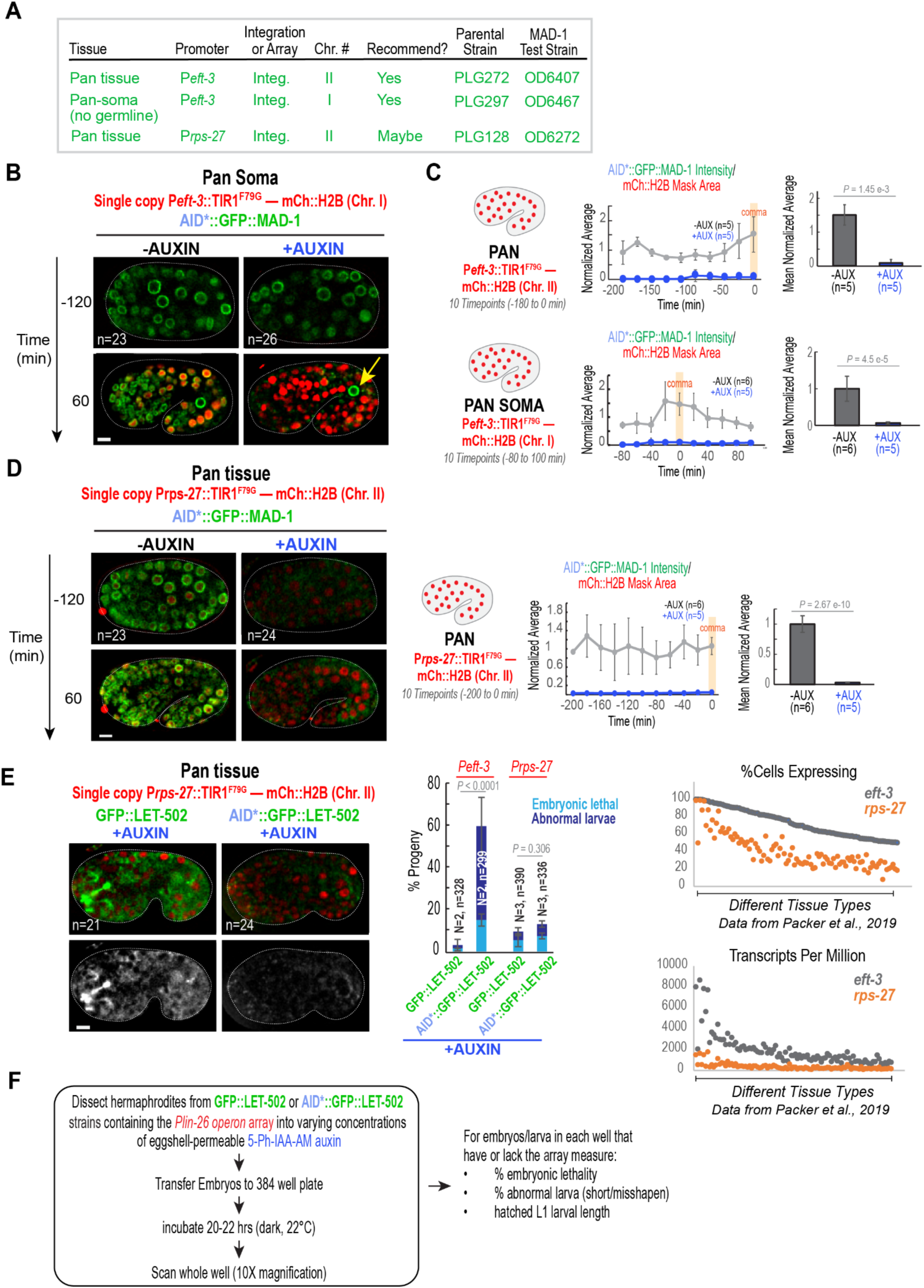
Characterization of pan-tissue degradation strains. **(A)** Table listing the pan-tissue and pan-soma single-copy integration strains, with target tissue, promoter, integration site (Chr.), whether the strain is recommended, and the parental and AID*::GFP::MAD-1 test strain numbers. **(B)** Timelapse images of embryos expressing AID*::GFP::MAD-1 (*green*) and homozygous for the pan-soma single-copy *Peft-3*::TIR1^F79G^—mCh::H2B operon insertion (Chr. I; *red*) in the absence (−AUXIN) or presence (+AUXIN) of 50 µM auxin. AID*::GFP::MAD-1 is degraded in the soma but retained in the germline (*yellow arrow*), consistent with germline silencing of this insertion. **(C)** Quantification of AID*::GFP::MAD-1 degradation for the two P*eft-3* insertions: Chr. II ("PAN," *top*) and Chr. I ("PAN SOMA," *bottom*). For each, a schematic (*left*), a plot of normalized AID*::GFP::MAD-1 intensity (AID*::GFP::MAD-1 intensity/ mCh::H2B mask area) versus time (*middle*; mean ± SD at each timepoint relative to the comma stage, timepoint 0; gray, −auxin; blue, +auxin), and a bar graph of the per-embryo averages across the indicated 10-timepoint window (right; mean ± SD) are shown, in the absence or presence of 50 µM auxin. Graphs show different temporal windows because Chr. II PAN expression initiates earlier than the Chr. I PAN, due to differences in germline expression. Quantified as in Fig. 1D′,E′. *P* values, two-tailed Welch’s t-test. **(D)** (*left*) Timelapse images of embryos expressing AID*::GFP::MAD-1 (*green*) and homozygous for the pan-tissue single-copy P*rps-27*::TIR1^F79G^—mCh::H2B operon insertion (Chr. II; *red*) in the absence or presence of 50 µM auxin; AID*::GFP::MAD-1 is degraded throughout. (*right*) Quantification for this strain, presented as in (C) (10-timepoint window, −200 to 0 min; *P* value, two-tailed Welch’s t-test). **(E)** (*left*) Images of embryos expressing GFP::LET-502 or AID*::GFP::LET-502 (*green*) and homozygous for the pan-tissue single-copy P*rps-27*::TIR1^F79G^—mCh::H2B operon insertion (Chr. II; *red*) in the presence of 50 µM auxin; merge (*top*) and GFP alone (*bottom*). AID*::GFP::LET-502 is degraded, including in the sensory depression region. (*middle*) Mean percent of progeny scored as embryonic-lethal (*sky blue*) or abnormal larvae (*navy*) for embryos expressing GFP::LET-502 versus AID*::GFP::LET-502 and homozygous for the pan-tissue P*eft-3* or P*rps-27* operon insertions, in the presence of 50 µM auxin; a significant increase was seen between AID*::GFP::LET-502 and GFP::LET-502 for degradation in the presence of the P*eft-3* but not the P*rps-27* operon. Note that the bars for the P*eft-3* operon insertion are reproduced from Figure 3H for comparison. Error bars, SD; *P* values, chi-square test. (*right*) Single-cell expression data for *eft-3* (*gray*) and *rps-27* (*orange*) across tissue types: percentage of cells expressing and transcripts per million; *eft-3* is expressed more broadly and at higher levels than *rps-27* (data from Packer et al., 2019). **(F)** Scheme of experiment to assess embryonic lethality, larval abnormality, and hatched L1 length across a range of auxin (5-Ph-IAA-AM) concentrations. *n* is the number of embryos/progeny analyzed; *N* is the number of independent experiments. Scale bars, 5 µm.

## SUPPLEMENTARY MOVIE LEGENDS

**Movie S1. Example movies and quantification of AID*::GFP::MAD-1 in the presence of the P*elt-2* (intestine) and P*elt-1* (epidermal) driven TIR1^F79G^—mCh::H2B operons.** Timelapse images of embryos expressing AID*::GFP::MAD-1 (green), along with arrays expressing the P*elt-2*::TIR1^F79G^—mCh::H2B (intestine, top) or P*elt-1*::TIR1^F79G^—mCh::H2B (epidermis, bottom) operons in the presence or absence of 50µM auxin. Each example shows the masked region generated by the red mCh::H2B signal (right panel) and the region within that mask containing green MAD-1 signal (second from the right) that was used to quantify the green intensity signal at each timepoint.

**Movie S2. Gallery of toolkit strains.** Timelapse images of embryos expressing AID*::GFP::MAD-1 (green), along with the indicated TIR1^F79G^—mCh::H2B operon expressing arrays (P*elt-2*, P*cest-17*, P*hlh-1,* P*elt-1*, P*lin-26*, P*dyf-7*) or single-copy insertions (P*cnd-1*, P*rps-27*, P*eft-3*, P*mex-5*) in the presence or absence of 50µM auxin. Movie pauses briefly to outline the tissues of interest in white for each strain.

**Movie S3. *let-502(RNAi)* and *gfp(RNAi) phenotypes*.** Timelapse images of embryos expressing AID*::GFP::LET-502 (green). A control embryo is shown along with a *let-502(RNAi)* and a *gfp(RNAi)* embryo as described in Figure 3E.

**Movie S4. Epidermal degradation of LET-502.** Movie shows timelapse images of embryos expressing AID*::GFP::LET-502 along with the P*lin-26*-driven TIR1^F79G^—mCh::H2B operon that expresses in the epidermis, in the absence (top) or presence (bottom) of 300 µM auxin. Embryos are the same as those shown in Figure 5D. Fluorescence is overlaid on brightfield images to enable visualization of the changes in embryo morphology throughout development.

## Notes

### Competing Interest Statement

The authors have declared no competing interest.

