## Supplemental Table 1 for "A tissue-targeted auxin-inducible degradation toolkit for *C. elegans* embryogenesis"

**Table S1. Plasmid information & sequences of regulatory elements**

| **Plasmid**  **No.** | **Expressing**  **Tissue (s)** | **Regulatory element** | **Length**  **(bp)** | **5’ end**  Preceded by:  Homology arm | **3’ end**  Preceded by:  Homology arm | **Template used**  **to amplify promoter (Reference)** |
| --- | --- | --- | --- | --- | --- | --- |
| pOD5555 | Intestine | *Pelt-2* | 2994 | 5’- TACATCTTTACCGGCACCAGAAG- 3’ | 5’-TCTATAATCTATTTTCTAGTTTC- 3’ | pOD2044  (PMID: 28619826) |
| pOD5562 | Pharynx | *Pcest-17/ K07C11.4* | 3390 | 5’- AAAGTGCAAAAACGTTAAAATGG-3’ | 5’ CAGCTCCAAAAAAGTGGCTTGAC-3’ | N2 genomic DNA |
| pOD5557 | Muscle (early) | *Phlh-1* | 3333 | 5’- TGGGTTAATGTAGGTGCTGGAAG-3’ | 5’- TTCTGGAAAATTATTGGAAAATT- 3’ | pOD1486  (PMID: 30890570) |
| pOD5556 | Muscle (late) | *Pmyo-3* | 2418 | 5’- GTAATTTATTTTAACCTGTACTC- 3’ | 5’- TTCTAGATGGATCTAGTGGTCGT- 3’ | pOD2046  (PMID: 28619826) |
| pOD5564 | Neurons | *Pcnd-1*  (non-ciliated) | 3230 | 5’- CAGCTATGACACGTGGCTCTAGT- 3’ | 5’- TGTAGAAAATATAACTGGATGAC-3’ | pOD2983  (PMID: 30890570) |
| pOD5565 |  | *Pdyf-7*  (ciliated sensory) | 3324 | 5’- TTCATATACTTTATGTACGGCGT- 3’ | 5’- ACTTGAAGTTTAAATCTGAAATA- 3’ | pOD2766  (PMID: 30827898) |
| pOD5563 | Neuroblasts | *Pztf-11* | 3535 | 5’-AGGTAAAGTATCTGAGAATTGAT- 3’ | 5’- TTTGATATAGATAAAGTCCTAAA- 3’ | N2 genomic DNA |
| pOD5554 | Epidermis | *Pelt-1* | 6490 | 5’- TGACGGTTCAGTTAGGGTCTTGTAG- 3’ | 5’-TATCCTCTTACTATAAAAATTAGAT- 3’ | pOD3678  (unpublished) |
| pOD5559 |  | *Plin-26* | 4072 | 5’- CGAGGTTAATATCTGAGCTCCAT- 3’ | 5’- AAAAGTAAGCGAGGGATGAAGGT- 3’ | pOD2018 (unpublished)  **see footnotes* |
| pOD5566 | Germline | *Pmex-5* | 488 | 5’- AAATATCAGTTTTTAAAAAATTAA- 3’ | 5’- TCTCTGTCTGAAACATTCAATT- 3’ | pLA024  (this work)  *see footnotes |
| pLA095 | Pan | *Prps-27* | 841 | 5’- TTCAATCGGTTTTTCCTTGCTTG- 3’ | 5’- TTTTATTCCACTTGTTGAGCGGG- 3’ | N2 genomic DNA |
| pOD5558 |  | *Peft-3/* *eef-1A.1* | 600 | 5’- GCACCTTTGGTCTTTTATTGTCA- 3’ | 5’- TGTTTCCCAACTGAAAAAAAAAC- 3’ | Addgene plasmid # 71720  (PMID: 26552885) |

***Footnotes:**

1. P*lin-26*:: TIR1^F79G^ —mCh::H2B construct has a minimal *myo-2* promoter sequence introduced in between the *lin-26* promoter and TIR1^F79G^ fragment to enable TIR1^F79G^ expression (see **Fig. S1**).

2. P*mex-5*::TIR1^F79G^::3XFLAG construct has a 3XFLAG tag and no mCh::H2B reporter.

3. For full promoter sequences (see below)

**Summary of promoter expression (from Packer et al., 2019** PMID: 31488706)**:**

- **P*elt-2***: intestine (anterior, middle, posterior and far posterior)
- **P*cest-17/K07C11.4***: pharyngeal muscles (pm1-pm8), supportive cells (marginal cell 1-3), gland lineages (g1A, g2) and pharyngeal neurons (precursors/neuroblasts for M1–M5 and I6)
- ***Phlh-1*:** body wall muscle (anterior, posterior, far posterior and head rows)
- ***Pmyo-3*:** late body wall muscle (anterior, posterior, far posterior, head rows)
- **P*cnd-1*:** non-ciliated neurons (GABAergic neurons (RME, RMED, DD), peptidergic motor neurons (RID), and interneurons (RIV, PVR) as well as a subset of ciliated sensory (ASJ, ADE) and interneurons (AUA, ADA) across the *C. elegans* central and pharyngeal (I5, I3) neurons
- **P*dyf-7*:** ciliated sensory neurons, including amphid neurons and outer sensilla/non-amphid ciliated neurons
- **P*ztf-11*:** neuroblasts, but not in differentiated neurons
- **P*elt-1*:** epidermis (AB ectodermal lineages, including seam cells, dorsal hypodermis (hyp7), GABAergic (DD) motor neurons, cholinergic guidepost neurons (AVG), and T cell lineages)
- **P*lin-26*:** epidermis (epithelial lineages, including dorsal/ventral hypodermal cells (hyp3-7, tail, P) and lateral seam cells, as well as in specialized non-neuronal glial and sensilla supportive cells (ADEsh, Amsh, Amso,Ilso) and a subset of specialized excretory, rectal and pharyngeal tube or valve cells (excretory cell, excretory duct pore, Y, B, K/K', anterior arcade)
- **P*mex-5*:** Germline only
- **P*rps-27*:** pan (most tissue types, including intestine, body wall muscle, hypodermis, pharynx.) but does not express in most neuroblast lineages
- **P*eft-3***/***eef-1A.1*:** pan (expresses broadly across all tissue types)

**Promoter sequences:**

**P*elt-2***: intestine (2994 bp):

TACATCTTTACCGGCACCAGAAGAAATCACCGCTAACCAATGTTCAGTTTTTACCTGAATATCCCTGAAAGTACAAAAAAATCAATGTTCAATCAAGTTTTTGCATGCCTTCGTGTGATGGCTCGCACCAAGTATTCTGACAGTTATAAAATGTTTCCTGTGATCATGAATTTGAAAAAAAAAGAACTGATGGGAAAATGTGATATTCATAAATGAAATGTTTAATAAACAAAAAAGAACCGTCATCACAAATCTACTCTGAAACGTTCACAGTTCCTAATTGACCAATTCAGAATTTTTTGTACTGACTTAATAAATATAAATTTACAGTTTTAAGAAACATAAATTGAATTATATAAATAAAAAAAAAAAAACTAAAATGAAGACGTGATGGAATGTTTCACGTATTAAAAAAGCTCTCGGACCGTTTGACATCCAGAAATTATGTGAAAGTGATTTTCTAACCCTTTGACGTCAAAAAAGTTTTTCTTTTTTGAAAGATACGCTGTTTAGAAATACTTATACAGCTGATTTTAGAATCTCCAAAAAACCTTCAATTACTTAACAAACTTTCTAAATAAAGAATTTTGATTTTGATTTTCTTTGAACTTCAAACCCCCCGTTAAAAAACCAAATTAAAAAAAAGATACCAGATTAATTTTAAAGTGAAAAATATAAATTTATTCCCGACTGTTAAAAACGGTTTATAAACGGAGTCACAACACTCATATACAACTATCGACACCAAATAAATAACTTCGAAAGCTAATTGTGTCATTTCGGATATAGAGAGTAGTGTGAGCAGATGTGTTTTTGCAGAACTAAATTGATGACACTATTATGGAATATAATGACCAAATGTTAAATGTGTTAAGGTTTGATATCAAAACCTGTATTTTCTTTTTATACAGAATTGATAATGTTATCTTCAATTGATTTCTACTTCTGAGCTACGGCGATACGAGGACGCATTCTCAACGATAATGTTGCCATTTTGTCCTGATAATTTTTTTACTGATTGTTTCAGAACACCCATAGTTTTTCTCTATTAAACGTTCATCCTTGACTTCCCCGAGTTTGCTGGCTGAATAGGAAATTTGAAGACAAAAAGGAAAGAATCGGCTCAAACGTCATGCAACTGATAAGGCGACCGTACTTCATTTCAAAAGAAGCTCACTTACTGAGCGCAAACATTGAGAAATGAGAGCAAAAGAAGTGGTTTTACAACATGCTAATGTTTTTAGACCTTGTACCCAATAATATTACTGTAGTATACAGTTCGGAGAGCATATGGTTGAAATCTTGAAATACCAATTTATCACTAGTTTGATTGTGTTATCGATGTATAAAGATATATTTTATCATTTTTGATTATTATCTGATATTGTGGGGTGTGAAGTAATATTATGTGCGTGTGTGGCTGATTATCGAAAAAAACTGAAAATTATCAATTTTTCTACAGGTTATCTTTTTTTGTTTTATTTTTCATTATTGTATTCTTCATACTCCTTATCCTGCCCGAATCACAAAAGTTCCAGACAACTTGAAGGTCGCATCACGTTTTGTTTATAAGAATGTTGAACTGGTCAGCTTTGGAGACAGAACTGTCCTCCAAATGTTGAATATGTTTCCGTAAAACACACTGTGTAACAAAAAGCGTACCCTTTGATCCGGTATTCTTGAAAAAAAACATAGAAGGGCGTACTTTTCATATCTCACAAATATGCGTTTTTTATTTTATAATAATATCATTTATTGATAATTATATGTATTTGTATGTTTTCTTGTCTTTAAAATTATGGTGTGACATCACAGTTTAAAAATTATATGAAAACTAATGAGTGAAAAGTTAAAAATGTATAGTTTGCCGTATTTTCTCCATTAGTATTGTATTCAAAACTTATTTCCAATTGGTGAGCACTATAAACTTTGTAGGGTAATTGAGGTTCAATACTAATTTTTGAATATCTAGCTTTTGACAAATTATTCAAGTTTTGACAATGATGCCTTATCAAATTAAAAAAAATTACCATAAATTCTTGAAAGCTTTTGATTTTGTTTCACTCTGTGAAGGATAGTAATTTATCAACATTTTAACTAGATACATCTGTTCTGGCAATAAATTAAAGCAAGACTATTACTATGTGCAGTATTAATAGCTGCATTGCAACTTGTTTGAAAGTTTCAGTGAAAACGCAAAAACTCAATTTAATTTTCAAGTTTGTTTGCAAAATTGGATATATGTATCCATCAAAAGTCTGATCGCCTGTTTTATCTACTTATTGTCTCCTCAGTTGATTTTTTTGTAGTTATTATAAACGCTACTTTGAAACCAATATTTTCTTCCTCATTCCACTTTTTTCAATTACCAACTACCGTACATGCAATGATGGGCGGACCCGAGTCCCATGTTTGGCTGGAAGTGGGTGGTTGTCTGCGTATATGAGCGACAGAGGTCGGGGCTGAAACTGATAAGAATAGTCGACACTAACGCCATAATCGCTAGCCAGCCATCATGCACACCGAGCTCGGTGTGCACACCATCTTTCTTTTCAAACCAATACGCTTTGTGCCTTCATTGACAATTTTCTTTTTGATAAAATCAGCCTATCTATACTTCCCAATCATTTTTAGTCTTATCGTTGAACAGCTATCGAGGTGCCACTGTTTTCACTGATATCTTCTAAGTTACTATGGCATTAACATCTTTGTGTTCTTATTTTATGGGTTATTTTAATTAATTTTTGCAGTTAATTTTTGGAATGAGCAAGAAAATGTTAATTGTAATATCTTCGTCTGAAAATTGTCTTCAAATAGTTTTAATTTTAAAGGCAGTATTTAAGAAAATACACTTCTCGAAGCATTTTTGAATTTTTGAATTTTTAAACTGCTTGATGTTTTAGGTGCCACTGTTTTCATTGATAAGTTTTGATGTATAAATGCTTGATTTTCTTGGCATTCTAATAAAATAGAAACTAGAAAATAGATTATAGA

**P*cest-17/K07C11.4***: pharynx (3390 bp):

AAAGTGCAAAAACGTTAAAATGGAAAATCCCTTTTCTTGAAAACCGAGAGATGAAGAGAACCGAGACATACAAACAAAACTCTCCGTAGAACGGCCTATGGTGACACTCACAAACAACAACATAATTGGGAGAATTGAGAGAAATAGTGGGAAAATGCGGAAATTTAGGAAAAAGAGGTTCAATGAACATATTTATAGAGACAATTGTTTTTGGACGGGAAAATGTGAACTTATGGGCTATGAAAATTTTTATAAAAGTACGATTACTTCCTTGCCGGAATTTCAATTTCAAATAATCAAAGCTGATAATTTAGTTCAAAATAATATTCCACAATAATCATATAATGTTTAATTTTCAGTAGTTTCATTCGGGATAGTTGTAAAGAACAATTTTGTGAACGATAATTGCAAAAAAATCGTCTTTTATCCAAAAATCTCATGGCAAACAAAATTATATTTCAAGCTTGTTTTCAAGCTTTAGAATCACATTTTTCAAGCTTTAGAATCAAAGAATCAGAAATCATAAGAATTTCATTTTAAGCTTTGATTACCACATAAATCATTTGGTATGGATGAATGATGCTTTTCGTTTTTTGGTTTTGTGACCATATTTTATGAGTCCACCGGAATTTTATAAAAGCTCGGGTCTCTGAAAAAACTGCTATAAGCCAAGCTGAACATTCGCTTGAAACAAGCGTGAAATGAAGTATACTGTCAGTTTGTTCAATTATTAGCACGGCAGTTGGCTATTGTCATTGAAAGTGAATGTTTGAGTGAGTCTCTACGGATTGAAACTTTTTTTTTTGAAGTTTGATGAATGGTTTGTGAAACAATATTTTTGTGTCTCGTTTTTTATTTTATAAAGTCAAATATTCCAAAAGTCCAATAGTGAATCACAAACTTTGGTTTATGTTTTGCAGCATTGTGAAGAAGTTGTTATTAGTCTCGATGCCATGATGAAGCTACAATAGTCTTGGGTTATAATAGGTGAACATTAAATTTGCCTGGGATGCTATAACAGAAATTTATGCAGGCCAAGGAGTTTCCGTGCCTTCGCAAACTTTTTTTTTCTACATATTAATTCAATCAAAACTTATTTCTTATGTACATGCAAATTAAGTATTCAGACTCCAGTTTCCAAAACTATTATTTCCAGAAAAATGTCTGTAAAATAGTGAGTTTGGAATTTCTATCAAAATCTCTTTTCAGTTATCTTTATCATATTTCAAGACGTCATAATAATGATTCATCACTTATTCACCATCTTTCATTTTTTTTTGTCACACTAATGTTATTGTTACACTTCCTGTCTCTCATCGTCTCATATCTCTCGGATCTCTGTACTATATCTACTCTTGACCGATTCAATTCGCTCAACCAATGTCATCTCACGATCAGTCTTGGGTTGAACTATCAACAAAATAACTATTTTCTTTTTTTTTTGTTTGTTTCAAAAACAAAAACTTGAGCCCAAATTTATAGATTCTTGTATTTTTAAAGTTTATGATATGAATTATTTGGCTAGAAAAATGAATTATTTGAGCAAGTTAATGTTATTTTGCCATCAAAAACAATTTTTGTGTAAAAAAATATATTATTACCAAAATTTTCCTGCTTTCTTCATGTTTCATTTTTTTCATTTTCACAATTTTTTTTCACCTCGAATTTATACTAATTCCATTTTCAGATGCAAAATTTATCTTTATCTCTAGTCATTCTATCAGTTCTCATTGCTGTAACTCTTGCTTGCAAGTGTAGAGAGCAATCAACCAAGGAATCATTCTGCAATGCTCACTGGGTTTCTCATGTCAAAGTGAAGGTCCGTGTAGGAAAACAAGGACTTCCAGAAGGATCCGAGAGAAAGGGATTGAATAATTTGAGATACACTGTGCAACATGTCGAGGTTTTTAAGGTAAGCGAGTTTGATTTATTTTACCCCGCAGGTGCTAAAAACTTACTGATAAGCTTATATCAGCGTTCTATACAACTGAAAGTAACATGCATTTCAAACTTACAGAAAAATGTTTCTATGTCAAAGTTTTTTTAATTTCAGAAACCATCCAACATGACAACTCTGCCAGATGAAATTTTCACTCCATCAGAAGCTCCAGCTTGCGGGCTCAAGATTGCCGCCGGTCATGAATATCTACTTGCTGGTCGTGTTGAAGGACCAAATGCTTTGTATACTGTTCTTTGCGGTCAAGTACTGCCAGATGACAGATCTCAAACGTCATTTGAAAACGTTCTTGAATGGAAAAATGTGAGTTGAAAATGCTGTTTGCTAAATAAATTGTTAACTTCCAGGTTCCACAGACATTGCAATCTCAAGTCAAGTCTATCAAATGCTAGAAAGAAATAAAAACGATTTTATTAGTATTTGATAATAAAACTTTAATGAATCTGATTAAATAAACTTACAATTAATGAAAAAAACTTTTAAAGCGAAGCTATGATAAAACAAACGATATTTTCATGTGAGTGAGCAGAAGTTATTGTTTGTAAAAAATTGAAAGCATTATTTTTTCAATAATCGGTATTCGATAATCAGAGGGTGACAGATGTTTTCAGCTGTAAGACTACAAAGGGTGAAAGTCTCACTACATGACTACACGGTCCCTATTGATGGCTACACTGAAATGGTTCAACAGCGCCATTCACATTTTCAGGTGTTCTATATATGTATGTGTTCTCTGCCGGGAATATTTTGTTCAAAACTTATAGCTTTAATCTTTGAATAAATCAAGAAAAAAATTCCTATACCCAACATCTATTAAAGTTATCTTTGGAGAAAAATTTCATGTTTCAGGTTCCTGTTGAGAAAATGTTTCCAAATTTGCTCGGGTGCTCTGAGATCAGTTTTAAAATGAATAACTGTGTAAAACTGTTTGAGTTTTCCAATGTTTGTCGTTGAAAACTGTCGTGGAATATTCATAAAAAAATTTGTCAACAACATTATGAACACGACTTTTTTATTCAAGCACATTTCGCTAAAACCTTCTGTTTTTGAGTTCTACCGATTTGAATAAGTTTGCTAAATTGTCAAAACAAAATTTTGAATTCATCTGTAGCTGGATTATAAATTTTATTCAATTCCCGTGTTCTCCCACCTTCTCATGTTCTTCTTGTCTTTTCACAAAGTAAACATAACACAATTATCACCACACATCTGACCCTTCTTTACCTTCCCCTACCATCTCCATCTTTCTTCATGTATGCCTGGGCTATCTCATATTTGTTTGCAGCACAAATAACTTCTCCACCTTTCGGCTGCGTCTCTTCTATACCGTCAGTCTCTCACCGTTTTGTTGCTCGTTTCCGACGTTACCTCCTTTCGTTCTGCTCATTTGCATATACCTGCCAAAGAGAGGTCAAGCCACTTTTTTGGAGCTG

**P*hlh-1***: body wall muscle (3333 bp):

TGGGTTAATGTAGGTGCTGGAAGGGTACTGTAGGGGTACGGAAGGGTACTGTAGGGGTACGGAAGGGTACTGTAGGGGTACGGAAGGGTACTGTAGGGGTAAATAATCTTAGGTGTTTTTCTATTGTTTAAAAATTAAGTGTTTGTTCCATTTAGTTTCTAAACTATTTTGCCAAAAAATATTCTCCTTTTCCAAATAAATTTTTTCAGCGAGTTCTCGGTCTAACAGTGTCTCCCGTTGAATATTCTTGTAAAATCACACGACGAGGTTTGGATCACAATCAGAATTCTGTGAAATAAGCGTTTTTTGGGATTCTGAATGATTTTTTTCTGTATATATTTTCATTTTCGATTAGTTAGTGTGTAATATGTGTTCTTCAATGAAGATTTTAATGCTTTAATTTTGTTTTGTAAACTTCGAATTTCCAAAATTTGAAATTCACAGTTTGGCAGTGGACTGGCGTAGATTTATATTTTTCTAGGCCATATAATATAGGTCGTGGTGAGACCCAACATTAGATTTTTTGCAATCAAAGTCCTGAAACTTTTCTTTATCTCTTTTCTGTTTAATTTCATTTTATCACCCAATTTTACGTGCTCCTCCGGAGACCGCCAAATTTATTGGAAAAGATACATGTTCTGGTGTCTCGGTCTTCTGCTGGTTTACAACGTGAATGTGCTCCTCCCGAAGAGTCTGCAAAATACCTGAACTCAAGTCTCCTCACATGATTTACGATACCTCCCGCATTCGGAGAGCCTATAGGGGGTGTCTGGCGAATGGGTGAGAAATGAGGGTGAAAAGGGGGTGTAAACTTGGAAAAGGAAAGAAGAAGACATATTTATGACAGTCCATGACTTTTTGACGGTCCGATGGTTTATGGCTATGGGGATGACAATGCGAACAACTGACGACAAAAATAAGGTGAGAAGAGATTTTTGAGAGATTTTTGGATGGGTGAGAAATCTGAGAAGACTTTATTCGAATTGTTCATATTTTTTAGAATTTTGTAAAAAATACTTCTGCTGTTAAAAATGGAAAACCTCTAGAGCTCTGTGGAAGCACAAAAATCAAGAAATCTCGCGTGAAAATAGTGAAAAATTTACTAACGTTGTGAACATATTCACACTTTGAAGATTTTCAGTGAATTGGCTTTAAAGGTTAGCTTTTATAACTTCAAAATGACTAAATATAAAAATACATCACTCCAAAAAATGTTAATTAAAAGATGTTTTAAACGTCTAACTTTAAATTATTTGTTTAATCAGAATTTGCAAAAAAATTAAATTTAGTATTTATTTAAATGTTTCTAAACACAACTGTTCATTTTCACGCTCTTATTTCAGGAAAATTTTTTCAAAACTGTAAAACAAAAACCATTTTTCACAGAATCTAAGGGTATCTGAAAGCTTAAAATAACTTCAGAAAGATATCAATTCCAGCTGTTTAGTACCTGAACTGTCTGTAAACGTTTCTTCTCGAATTATAGAAAATTTTCCACTTTTTCAAGTTCAGGTTTTCAAGAAACCCCACGATTTCCACTCATCGTTTCCAATGTCCAATTTCCCATCCAATTTGCCGCACTCTGACCCAATGACTTGTTCCTTTGCCAATCAATGCTACCTAATAAATTTAAAAGTTTAACACGCATCCAATTGACACACAGGTAACCGCCCTTTCTTCTTTTACATAATTCGGAAACTTCAAAGAGCCGAAGGTGTCGGTTGTAGCAGCAGCGGAGGAACGGATGCCAATTGCGCAACTCTCGGCTCAACTCTTTTAGTGCTCCGAGAGCAGGAAGAAGAATACTGTTGGGTTGTAATAAAGACGGATGTTTTTGTTCAGAGTAGATTAGCTCGTGTTTGATTGGATTTGACCGGATCAAGAGGGGAATGTCCTGGTGGAATTAAATTTTATTAGAATAAATTGTATTTGGTGTTTAAATTCGAATCAATAATATTTATGAGCTTTAATGAATAAGTGTTAGATTATATAATTCTATAATTTTTGAACAAGCAATTCAAAAAGAAAACAAATTTTAGTAACCTCCGAAATCAAGCTGGGTGGCCTCTAGAAGTTTTGAAAAAACTTTTTTTATATTCTGTTGGAGTTTTTTTAAGTTTTATAATTATAGGTTAATCTTTCTAATTTGTAAGCTTTTTCTTAACCAATTTTTTTTGTTAACATTTTTTTGGAATTATGCTATATGACCTATACCTAAAACAGTTTAAAAGTTTAAAAAAATTTTCTATATTTTTCACTTCGTATTGAACCTCCTGGGTACATATATTGACAGCACATATCTTGTTTGTCTCAGATTTTATCAAATAAGTTTAACAAGTAAACCATGCACCAAATATTTTTCTAGGTCTCTGTAGTTAGGAAATATTTAATAAAAATAAAAATAACCGAGATATGAGCCATCAAAGTAGATCAATTAAGGCACAGGAAAAAAGATCTGAATAAAATCGAAGTTCTTAAAAATATAAATCAAACAAAATTTTTTCCAGAATTTCAGCCGAGAATTTCCAGCCGATTTGTTTATATTTTCCACATCACTCCCCACACTTCTCTCACACAAACACGATAAAATCTTGAGAAGCAATTAGCGCCAATCAACTCAACACAAAAACGAAAAGCCAACGAAAAGCTAAAGCTATCATCGTTGTCGCGTCTATGAGCAACTCAATCGTTCATCATCCTCATCTTCAGAGTGCTCAAACCTACCGTAACCCGAATTGGGCGGAGCCAAAGGGTCCGAAACAGTGCACCAGGGCGGGGAGGGACCCTGAGAAACGAGAGGGAAGTGAGCAATTGTTGAAGTGTCAGTTGTGCTCATCGAGGTCCGATGAAGAGACGCGCCTGCTCACCTACACAACTGACTTCCCCCATATACCTTTTTCTAGAATTTCCTTTTTTAGATTTATACGGTCAGGTAAAAAGGTAGAGTTTTACAGTGTAGAAATTAGGAAATTGCTCAAAAAGCCGAGCAGAATGCATATAAGAAGTACCATAGCCCCAAAGATTCGATTTTCCAGGGTTCAATCAATTTTTGTACTTTGACAGCGTATATCTCAGTTTTCTTTGATTTTATCAAAAACTAGTCAACTGACAAAATACTTGAAAAGTATTCCTTTATATTTTGGTAGCTGACCATTGTTTGTTAAAATATAAGGGAATCGAAATGTCGGTTATCAAAGTAGAACCTAACCTAAATCGCTATATATGCTATTTTTCAAAAAAAAAAACACGTTTTACTTTGTCTCAACTTATTAATATTCTTTAATATTTTTTCTATTTCTACCATTTTCCAAATTTTCCAATAATTTTCCAGAA

**P*myo-3***: late body wall muscle (2418 bp):

GTAATTTATTTTAACCTGTACTCTATCACTGCCGGCTATAATAAGTTCTTGAATAAAATAATTTTCCCGACAAAACATGAGTATTTCTTTCGAAAATAAAAGTGCAGGCTAATTAGAGATTATTCTGTAATTAACTGCATAATTTGTCACGTGCCATAGTTTTACATTCCACTACGTCATAGTTCTTAAAATACTAATCTCCTGAAAATAGAAGTAGGTGAAGAAAGTTTAATTATCAGTTCTAAAATGACAATTGATCTTTGGAATATGTTCTGAAACTACCGATCATTGAACAGATGCTATTTGAATGATATAGAATTGTATATTTGCAATTTCTGAAACGCGTTCTTAAAGGCACACAGATTAATTCAAAAGGGTCTGGCCGCAAAAAGGTTTATGGTGGCCGATTTTGAGTTTTGTGTGTGATTGCTTTTTCACAATCAGTGTTTTCAGGATTATGTGATGAACTAGATCTTCAAGTTTCGTTACATTTCATATGTTTTCGGAACTCACGAAGTACATATTGGGTATTGTGCTCAAAAAATTCAGCAATCAGCTTCGCTCCGCTGACTTTAGAACCCAAAAAAATAGTATGGCCAAACTGACTGTGTTACGATCATTTCAATTTTTCAATACATATTTAAGATTTCTAAGAGTAAGAAGGTCAAAAACTGTTCTGGAATACATATATATTTTTCAGGTTACAATTAGTCAAAAAGTGCACTGAAATATACGTTTTAATTTCACGAATAACCCAATTAGTTCAATGTATTTTTGGTCAACCAACGTTAAAGTTTGGCTTCCAACCAATTATCATTTCTGATCAACCACAATGTTTTTTCTTTATCTGCAAGTTAATTTTTTATTTTTATCCAGATGTTTGGCATATTTTTCAATTCTTCACTAGCGCCCACTTCTTGCACTTCCGGCGCCCTGAATCTAATGCATCTGTTGCAAGAATTGAAAGACCAATCAACACATTGTTTTCTTCACGAGATACTGAAGAAAATGAATAAAAACAGAGAAAAAGAGCCATGTGATTAGTGACAACTGTTGCTAACAGATACCATAGCTTGGACTTGGTACGTGATGGCAACGTATGGGTCAACAAAAATGATTGCAGAGGGGGTGCAAAACAGTCAAGTCGAGAAAATATGAAAAACAGAAAACAAAGAACAGAAAAATGGGTTTGAGAGTCAGTATAATTTATAAAAGAAAAATTGTACATAGAAATTAACCATTTTTGTAGAAGAAGTTATTTTTCAAGCATCGTTAAAAATTATTCAAAGCACCTTATTTCATATTTAATTTTAAACATGGTTAAATGAACAACACGGTGCGCAATCAGGAAAACTTGAAATCTGAAACTGTTGTTGTGATCTTCTTCGCAACTGTTCAGATAGCACTAGTGTAATGTTAAGAGTGCGCGAATATAATGGAATATAATGGATCACACCTCCTGCCATCAGGTAAACGTCTCTGTTATCACATATTTCCAACTATTAAATTTTTACCTTTTACAGTTTTACATTTTTTTGAAAAAAGTAACTTTTTGTCTTCAAAATCCCTGACGAAAATATCAAATATTTTAATCGAGACTGCAGAGGAACCGATTGATGATTTGGAAAATCCAGCTTTACCTGTGTAAGAACTGAAAAGTTTCATAACCCTAGGGTATTCCCAGTTACATTCCCCACTGGCTAACAATAGCACCCAGTTTTTCATCACCTTTCTTCAAATTTCTCGGCGATTTGTTAAAAACAAAATTTGTGTCCCTTCTCTGATATCTCTATGTCTCTAAACACAAGTTCATCGGAAAACGAAGGAGGGTAGGTGTTGGTTGGGCTCCCGAAGTGAAAATAGAAGAGCAAGAATAGAATATTAGAGAGAGAGTGCAGAGAGGGCGGGATAGCTCCCGGGATTCCGTTTTCTTCTTCTTTATCTTCAACGATGATGTGTGTGCGTGTTGTATAGATTCTGTTGCTCCCCCACAACTCGCTCCGAAGGCTCAATACAATTCAATTGATATTGGAGGAGAGCCTACCGGAGTGGGAGGATAAGAAGAAACATAAGAAGAAGAAGAAGAAGAAGCATGCTTCTGGTTTTTGATGCTATGAAAACGGCACAAAAAGATGATTGAGGTCCCTTTTCAATACCTTCTCTCATCTTTCAAATCCCATTGAAACCTAAAACTTCTCACCACGCTTTACCATTGTTCTCCAAAAACTTATAGCAATGTCTATAACTTTTTTATCTCTGAAAAGCAGTGTTCCATTTTTCTTTTTCCTATTTTATTTCAATTGTTTCTCACATTTCGTTTGGATTCTTTGCTTGTCAACCAGCTTCTTCTTCCACTTTTACCGTCTAATTTTCAGGGCAGGGAGCCATCAAACCCACGACCACTAGATCCATCTAGAA

**P*cnd-1***: non-ciliated neurons (3230 bp):

CAGCTATGACACGTGGCTCTAGTAATACTTGAAAGTTTTCGCGCATACTTGTAGTGATACACTGTATCCATCCGGTCTGTATTTATTCAGAAGTCATGAAGAGGGCGATGAACACCTGCTGGGAAGAGACAAATATAGGATATCTGTCAATAAAAAAGATATATGGTAACAGGTACATGTGTATTTAATGATTATTACATGCGCGATATATGTGAATGTTTGCGCATTTGGTTGCAATAATATTTGGTTTATTTGACATATTATCACAAAATTGGGAAAAGTTCAGGAGAGGAAACACTTCACAAAAAATTATTCGGAATACCCTTATTTGAAAGGTTTTGCTTGGACAATTCGTCAAAAATTAAACTATGGCAATGTGCAAAAAAAATAATTTGGAATTTTTCCAAAACTATTGGTAGATCTAGAAAAAATCCTCTTTAACGATATTCCTTCATTAATTAATTGAAAAGTTTTTAATAAGTTGAAAAATGTTGAAAATATTTTTTACTATATCTTTCTATATACCCAATTATTGTACCAAATCATCACCTAGGCACCAACCATATTGTTATTCATTAATAACCGCACCGCTTGCTCTCTTCCTCCCACCGATCGGTCTTGAAAGATTAATTACAGACTTGCATCATCATACAAATATACCGCCCGTACTCCTGAAAACGTTCAACATACGGTGCCAAATCCGTTTAGAAATGTGTGACAGTGACAAGGAATTGGTCTAATTAGGTGTGATGCAATGCGGTAACCAAACTGTCAAAGGATGGAATTCAACTAGGGGATTTATGAGTCAAGAGGTGTGAAATTTTGGTATTTGGTTTCTAGCAGAACAAAAGGTATATGAACTCAAATGAGGAAATGTGATAATCAAATTGTGAAATTATTGGTTTTAGAATATCCATGAAACTTAATTTTTCAGTGTCAAAAGAAATAATCTCGAGCCTTTTCAAATTGCTCGAAAAGTGGATTTTTGGACGCTTTCGATAATTTTCAGAGATTCTAGATTCCACGGGGAAAAATCTGCACAAGTACTTGCCTATCTTCCTTCCATTCCGGGCCACTATTTAAGTGTTCTTTCTGTGTTTTTTCCTAGTCTCTTCAAAAGATGAATGACCAACTGTTGGTGGCAGCTGGCTAGATAGCACATGAGTATAGGAGACTAAGACATTTGAAGGACTTGAATTAATTATTGCCAATTTAGGTTAGCTTGAGGATGGACTGAGGCCATTACCGGAAGAAGGGTGTTATTAGAAAAAAATAGAGGAAAGAAAAGAAAACAGTTATGATTTAAATGGAAAGTTAATTGGCATTAAGTGAGATTCTCATTAACTCATTATTTGCGCGAAAGTGGTACATTTCTCTTGAACTAAAACAAACAAACTGAAATGTATTCAGTAAATTTTTTCGCTGAAGTTTTTTCCAAAACTAGTCTTGGTTTTAGGATGTTTTTTTGTAAGATTAGGTTTTTATTATATTATTTTTTGATAAAATATGTTAACTTTATAACCAACCTTTTCATGTTTTAAAACAAAAAGTGACTAATAAAGTGCTCTAAAAGGCTTACTGAATGCTTACCAACTAATAACAGAATCTGCAGATTAGTGAAGTCTCAAAAACATGCAATGAAGAAAAAAACTTGTTTTTCAAAACCATCGTTTTTGTGAAAAGACCTTATCAACTTCCAATCAGTTCCCGCGTCAACAAATATCTGTGATCTATCTAGAACATTTCTCAGAACATCCCGTGCACTTATCTTGAAGAGTATGTCAGTTACAGAATTGTGTGACCCTTATAAAACCTTTCTTGGTAATTAGATGATATCTGCATTTACTGATGCACATCATTGCATAATTGGAGAAATATGACATAACAACTTTTACTGGATGTATACTACTCTATTTCGATAACGGTCGTATCGATAGATCCTATAGAACAAAAATTCAGACTATATGTTTTCACTCTTGGACCTTCACTCTCGGAATCGAAAAAACGATGATATTTGAATTTGAAAGAATAAGACAATCTATGATCTCAATTTTCAAATCAGAATTTGATTTTCAGAATTTTTTGTGGAATATTTCATATTGATGTATTGTTTATATTGACCCCTGGCAAAAGTCTCACAGTACAATGAGATCCCAGAAGAACACTTGGATTATGTTCAGGAAGGAAGAAAATATTACAGTTGACACCTCTGGTATGTAGGATTACAGAGATTTTCATTGTTTCATGAAACTTTGACACCTGCTCATTGACATTTGTGATTTGTAAAAGGTCAAATATTTTGAACTGACAAGTTTCTAGTAATGGGCACTCTACACCTTTGTTAGGCGGGAATTAATGGCAATTCCAGAAATTCACGGCAATTTTCAAATCTATTTATATTTCGAAATTTAAAATTACAAATTGCAATTGCGTGCATATCTGACATGCTCTCCGGATCTAACAGAAGATCAACGAATGAGATAGTGCTACTGTTTTAGGCAAATAACATGTTAATCGGTGACACGCATGTGTTTTGGTTGAGATAAAATCATACTGTAATTAATATGGAGTAGTTGAACTACCGTGGACGATTTCTAGCATGTTTCAAGATTTTTAAAATTACAAGCACGAGCGTTTTTGTTAACTGGATCGTTTTCAGAACAAATTTTTTATCTATTGGAAGGGATTTATTGAAATGTGGAACACCATTCCACACATGTTATAAAGCTTCGTATTTTAATATTTATAGTCAATAGTGTCGATCATGTTACAAGTTTGGCACATACATTCAAAAATATTAACTTTTTGGAATGCAGTTTTTTAGTTGGAACATAATTAAGAGTCCTTTGCCTTGTTGGTATACTTAGAACATTTTGATATTTTGTTTTCTAAAAAGTAGATAAAGTAGAGCTAAAAGTTTTTTTTAAAAGTATTTTTCTATGCAAGGTTCCTATTTTAAGCAAGTTTTCTAAAATCTGATTGTAAGACACATCAACATTAAACTTTTTCGAAAGTTTCATCTGTAAATTTATCTTTTTAGAACTTTTTAATGCTATCATTGGAAACCATTCCATCCTTCCCAACGAACATCCCGCCCCTTTTGTACACCACGGGCTCTCTGTTCACCCAGCTCCACCAGCCCCGCCCCTCCTTCTGTGAACCATCCACCAGATCAGTAGCTCATTCTCTTTCACTATACACTTCCCTGTCATCCAGTTATATTTTCTACA

**P*dyf-7***: ciliated sensory neurons (3324 bp):

TTCATATACTTTATGTACGGCGTACACGCACTACCATATATCAATTCTGCATCCAACTTTATTCTCTATGGACTTTTAAATCGTCAGGTATGTTGAATTTATAAGTACGGGTTCAAGAAAACATCTGGAAATGGAAATCGAATGTTTTTTGTACTTTTAAAAATTCTCACTGGGGATCATACTGTTTATTTAGGTAAATATTTTAAATTTTAGAAACTTCAGTAAATTCAGATTTACTTAACAAAGGCATGAAAGCGCAAGGGTATAACTTTTAGACATGTTTCAAAAATTCGATCTCATGATAAAGGCATTCCATGCATTTTCCAGACTGATTCTCCACTCATTTCTAAAACAATTCCTCAGCCGAACCCTAATTTTTGGACAAATAAATATTTTTCAGCTGCACAATGCTCCAGAGCGAAAATACACTAGAAATGGAGTGGGAGGCCGCCAAATGTCACATGCATTGACGACGAACACCCGACCTGAATATAGCGAGCTAATCGCAATACGTAAGTTGTTATCATTCAAGTTCAAAAAACTCGAATTCGGTTTTTTTCAGCATCCAGCAGTTGTCGCCCGGATTCTCGAGTGTCGGCAATGATCCATAACAACAATAACAACACGGAATCTTTACCTCTGGCACAGAACATCTCGAATATGACTAACAAAGTTTGAATTTGTTCCAATTTCTATCATGTCTCTTGTCAATTGGTCTCATCACGTTTTTGTGTTTTTGCTCTAACTTTCAAAATTTATTTCTAAAACTGCTTTTTCCTACACACCAGCACGTGACTGTCTATTACCTGTTGCTTGTCTTTTCACCTTGTCTCGCTTCTGCAACTGTCTCGATTATCTCCATGCATATTTTGTAACTTTATCAGGATTCTGCAACTATTGTTCCAATGCCAATGTCTGCCAACGTCGACGGGAATGAGATCTACAATTGGATCACTCCAGATACGGAGTCTGTAATTCTGTAAATTATATTGATACCCATCATCGTGTACAACCTGCTGATTTGCATTCTTTTTCTGATTTTTCATGCTGTTGTTTTCTGCCACCAGTTGACTTTCAGTCAAAACTGATACAATACTTTCTTTGAAGTGTACCCTTGCTTGACTAGATTCCCTCTTTATGCTTCAAATAGCTGCAACTAGCTAAACGCTCTGCTTGTCTTACAATTGGTTTCGGGACTTTAACAATTCCAGTGCTAAATTATTTTCCAGGTTTAGTAAATTCATGTGCTTCTTATTGATATTATGATCTCGTTTGTAAATTCTCATTGATCTTTTTATTTCTTCCAATTTCTAATGCTTTTTGTTTAGTTTTTACAATTTCTCCATTAATAAATGATTTTAACGTTAAGATAGAATCTGCAAACCAATAACCGTTGCAAAAAAAGTCGAAACAGTAAATGAAAGACAAGAGCCGTAAATATGCGTTATTATAATTTGTAAAACGCAACGTTGCATATTTTTTTTAACTGCAAAAAAAATTAAGTTTCGAAATACTTATTCGACAAACATCATTTTTCAACAGTATTATATCGAAAACACCAGTGAGCACGTAATGTATGCAGTGACCGTACTGATCATTTTAAAAACAAATTTGCAAATGTTTCACAAATCCTAGCTTGACACATAAAACTTGCAGAGTTCGTCTTAACATACATAATTCAAATAGTTCTTAATTCAAATAGTTCTCATAATTCAAATAGTTCCCAAAGGTGTTCCCAAAATTATTGGTGCATAGAATTTTTTTTTAATTTCTGAAAAAAGCAAACATAAACCCCTGTTATCTGAAAAATGTAGTAGCGAATGAAACTTGCACACACTGTAAAATAAATCAATTCATTTGCATTTCCCACGTGGAAACAGTTTTCTAAGAATATATTTCTTGCAACAAAACCTCTACTTTATTGCCACCTGGTTTCTTCGTATTAATGAACGTTCACTGTGCTAATTAAGGTATCGAGATGATCCGGTTAGAGCGGTATGCCTTCTTTCACCCAGGTGGTGCTCATTATCGGTGCGGTTGTATTCATGAAATACAACTTTGCTTTGACATTGCCGAATTTACAATTGTGTGTCCAAAAGGGCCACCTGAATATGCAATATCAGTCTCATGCTAATTAAGTGAATGTATATGATGTATTGCTGGTGATAAACACTTATCGTGTATGTTCTCTCACTAATTATTTATATCCTTTGCAATGTCGGTTGCCTAGTTGAGCCCGCCCATTTGCGTGTCATAGAAACTTGTCCTCCACATTGAAAGGTAAGTACGACATGCCGGTTGGTCCCAGCAAGTCTCGACGATCTATCATTTCTGTCCACATATTGGCTCTCGTTTTGCTCCACTTTCCGCTTGTTATGTTTTTCAATAATGGGCAAGCGAACACCTCACTTATTCATTTTCTTGGCCCTCATTCACCGTCTTCTCTTTTTCTCGCCCACCATACCGACCATCTGTTGGCATACGCCACTTCCGAGGCAGGTGCCTTTGTCCGGGAGATGAGGGCGAAACGTAGGTGTTGCTCACCGTTCTCTCTCTCCGCCCCCTGAAAACAAGGACTACTAGTGCAAACGTGAAGCTCTAACCTAATTTATTTGCCATTTTGTTTGATAAAAAACTATGCTTTAGTTTTTAATGCAAAAAGATGCCGTTAATTTTTTCCGAAAGTCCGTGTAGCTACGTGAAAAACTAAATTTGTAACATATTGTAACGTATCCATACAATTTATGTTTTGATCCGATCAGTATGTTCAAGTTTTTAGCTTGTTGGATTTGTCTATATTTGTTGGTTTTTTTTTCTTATAGCTCAGAATGTGAGCTGGCAATAAACATGGTCCTATAATTTTCTGTACTTGAATCTAACCTGTATCTCTGTATCTACCCTGAAAACCATAATTTTCAACAACAATGTGCCCACTTTTTAATCAAAGCTTGTGGATAATTGATAAATTCAGTTGAAAGTTTTCTTTAAGCAAAATTTTTACACGTCAGAAACGAAAAAAAAAGTTTTAAATTCTTGTTATTCTTCAATCATTCAAAAAAAAATTTTATTTTTATTTTTCATAATCCGACATACAGCTTTTTCTATTTTTACCCCAATCATTGAGACAATAGCTTCTCCACGATTAATTGGGCGGTTGGCAAGAACTGGGTGCACCTTTTCACCCACCTAAAAGCTTAACGAGCTCACCCCATTTTCAAGATACCATAGCAACCTTTGTGACTACCCAACCTAGTCTCCTCATTTGTTTTAGGTAGACCTTTGTTTCAAACAACACTGTTTCTATTTCAGATTTAAACTTCAAGT

**P*ztf-11***: neuroblasts (3535 bp):

AGGTAAAGTATCTGAGAATTGATTTTGTGACAGTGAGACATTTCAGCTGGTCAAGGCGTGACGGGGGGGGGGGGCTCTCTATTCGCCCTACTCTCGCCCGCTCCACTTTTCAGAAGAGACTAGCGAGGGTTTGCCCATTGCCATTTTCCGGAAAACTTTTCATTTAAGGAAAAAAATCTTGAAGTTCATTCTTGTCTTGAAAAGTTCACTTATTTCTAGGTACATTGCATCTATATTAATTATTCATCGCTAATTGAAACTTTGAGAAAATTATATTTGAGGTCTTCCAATTTCCAAAAAAAAGTTTCCTCCTGACAATGAGAAATATAATTATTCGAAAATTGTTGTGCATGCGAAAAAGTGGGCAATTTTTCCAAGTTGTGATTCATAGTTTTTATATCGTTTTATTTGTTTCAACTGGGCATTTGGAATCAAAAATAAATCACATGATTCAAGTTTTTAACTGTCAATCATCAATGATTATTCTTCAGTTTTTGTTTTCAATTCAGAAGCTTCACACATCAAAATTTAATTTTTAGTGGAATTTTTGAGTTTTTAGAAGTTTAGATATTCTACTCGGTAATCAAAAAACTTTCCTACGTTTTTACACCGATTTGTTTGATTAAATTTAAAGTTTGTCATTTCAAAAAATCAACTTTCATAAGTTTTGCTATAACTCAAAATTTTGAAATAAGTCCAAATTTTCGGCAAAATTGTTCAATTTTTCAAACTGAAAATTTCTGGAGTGTTCAATACTGAGATTCGGTCATGTTCCAGGTTATTAGAAAGTCCCACCGGTTCAATTACACCGTTAACTATGCGGTAGGGTACAACAAGCTACCTTTGAAGCTACATAAAACCAACAAATATGCTTATTTTAGAATATATTTCTTCTAAAGTAAGTTTGAGGACCGCATTGGACTTCCTGTAACTTGAAATTTATATGAAATTTGATCTACCCCTCTTCACTCAACAAGCCATGGTCTACTTACCTTTTATTCAAACTCATTATACTCTCGAGTTTCTTCAAAACTTCTTCCATTTTCCCTTATTCATTCTACTTCAACTTCTTTCCAATTTCGGAGCGAAAAAAAGTTGGTCTTTCTCATTCATCTCGTCATTCATCTTGGCGCTTGTGTATGTGCTCGGGAGGCCTTCCTTCTTCACCATATTAATTCAATTTTCATAAATATATCGCATTCTACAACTAACACTCACTTTTCTATTCCAATATACTCTTTGTTTTATTTCTACTTTTACACGCGTCTTCAAAACCCATAAGCGACCGCCTTGGACCAACTTTTTATTTTTCGAATGATGAGTACAACATCTGTCCCTATTTTCTGTACTATTAACTTTATTATTCTGCACTTTTGAATATGAAACTTTTAACATACTTTGATTAATCTCAAAGTACTGAACTCCAAATCCCAATAGTTGAAAGTATGCCTCTATAACTCCGCCCTCCGTCACTTCTTCTTTCGACTCCGCCTCCAATTCTCAACTATTGGCTAGAGGAAGCCGCCGTGCACAACTCAGTTGTCTTCTACTCTTTAAGTTTTTACATCGTCTCCCTCTCCAACCATCATTAATCGTTTTGTTCTTTTCAGTTACTACTAGGGGAGTCGTGTTTCGGATTGATTTTTTTAAAAATTTAGTAGGAAATGACACATGATACATCACATCATTAAAGTTCAACGATCGTTTCGAAACAAATTGGCCATGCTTGTAGTTTGACAACACTCTTTGCTCACTTTGTAACTTCCATTATTTTCTTTCATGGTATTTTGTAGATATGTTTTTTTTTCTAATAATGTAAATTAAAACTGATATCTAGAGTTATCTAGAGTTGCGAATTCACATATAAAGTGTTAAGTTGGTTGATAAGCATGATGCGCTTTTTTCTATTAGTATTTCTCTCTTCAAGTTGGCATTCTATATTTCACTACATTTTAAAAATACAAAAAGTAGTTAATTACTAAATTGTAGAGATAGAAATGTTACATTTGTGCTAGTTGATAATTTCTGATGAAAAAAAACATCTGTCAAAATTAAGCATTAGGGTTCAAGTCTGATAGAGTGGCTAGTTTTCCAGATGGTTACTAAACAAAAGTTGGTAGCATACACTTCCAAACTTGGTAGATCTTCCCCATTAAAAGTCCTATGATTCAAGAACCTGCTCTTTCCGTAGTAGGAAAATACGGCTTATACTTGTGGGATTTCAGTTTTAATGTTTTTCTCGATCATATAAGTAAATTTAAAAACATTTCGTTAAACGGTCTAAAATTAAAAACTGTGTACCCTTGTAAGATCTCTATCTAATAATAAGTTTAGTTTGCACTCTATTAGGTCTGCCTGTTGTCAATACAGTGACCTACATAACTTCCCAATAGTATCTATTTCTGTGTGCATAACCAATATACATTGCCACGTCATAAACGAGATCCCTCACCTTTTATTAGGTTCTCCTCTTGACCGCCTCAGAGTTTGGCTTATCTTCTGTTACATTCAAAACAAACAAAATGTAGGATAATCATTCAAATTTCTGTAATCTTCCGGTTCACACTTGAATGTGAAACTGTTACTATAGTGACAGTGAAGATATGTTTTAGGATTATGATTGATTTTGTTCAAGAAGAAGATCTACCAAACTTGGAAGTGTACACTACCTACTTTCTGGGGTAATCTAGAAATTCAGACTTTATTTGTTTTGCTTCTAAATTTTTCAACTTTGCTAGGTTGTATTCTGCTATATCAACAGATTATCAATTGGCAAGAAAATGTTTATTCATTTTTCTACCGTTTTGTAGTTAGTTACTTTTTGATATCTGTTAAAAAACTATTTGTTAGGTGAACTTGTGTAAAGTTTCAAAATCCAAAAATTATCAGTGAATCCTGAAAAATTACAATTGTGTATCGTCCATTGACAACTTTTCACATTTTCAAAATTATTATACATACAATTAGCTTCAAAAGTTTGAAAAACTCCATATCTAGACTATACTATTCATAGCTCTTTATTCAAAAGTGCCATAGACAGGTGTCGCATTACAAGTGTCTTGTCTGTACCATCTAATCTGATGGATTTGATCTCCCATACACGCGCAACGTAAACTATTATTATTATTTTCCATCCTCCCATCAAGTGTTCTCCACTGAATACCATGCACATCTAACTTTACCTAATCCCTGCTGTCCCATCTCCAATAATTCATTTCCTGCTCTTCCACACCTGCCCCCGTAAGCATTATCGGGGGCCATTGGGGTTTAACTTGGTACCCGAACGCAACCCCATTACTGTCCAAAAGAAGGCGGAGTCTCTGTATGTGCGCGGGCAATGCATGATGCCAGGTGCACCACGCCGCTAAGCAGATGGTGTCCTCCTCCAGACCGCCCTCCTTACTTTGAGTGTCTATGTGTGGTCATTCTGTGGTCCTTCTACACTTTCTTGCTACTTTTCTTGTTTGCTGAAAACTTTTTCTCTGTACCTTCTTACTATTTAAAAAATATGTTTTAGGACTTTATCTATATCAAA

**P*elt-1***: epidermis (6490 bp):

TGACGGTTCAGTTAGGGTCTTGTAGCCTGCTCATATTTTTAAATGGAATTATTACTAGAATCTGATAGTTTCCCAACATTGACTGCGGTTATTGGCAAATTTCCCCAACTCGCTCATTTTGGTTTACGACTGACATTGCCCGTCTGAATGACCCAAGGTGGTTGACCCTTCCCTTCTAATCAGGTTTGAATTTTCCTGATCTGTTTGAACATAAATTATGTTGCTAAAAATAGAGTTTCTTAGATCCGTCTATCTTTGAACGTTTAGTTAAAACCCAAGCACGAAAACCATATCTTCAATTCAAACATTTTTAAATGTTTCAAGGTAAATTTGAAACTTTGGCGAATGATCACACTAGCCTGAATTAGTTGCACTCAATCAACTCAAACAACTTGTTATATCTAATAAATATTGATAAGCCAGTCCCTAAACACCATATTCTTATTTCTCCACTTATTCCACCTCATCCATCCATCCATCCATCCATTCATTCATTTGTTTCATGGTCTTCAAAATGTCCGCGACGACCCGAAGTCGTCGTCGGCGTGAAAAGAAGAAGAAAACACATGAAGACGAGGGAAAGGAATAAGGTACTGTAGAGGAATAAGGCAAAGAGGGATACTGTTATCGACTCGGTGAATCTTGTGCGAGTGACACCTACAAACACGGCTTGTGAAAGAGAGAGAGAGACAGAAATGAGCGGCCGCCTAGAAAAGAAGACGAACTTTGGTGACCTCATGTTCTGAGGAATCTGGTTGACCCAAATTCATTCCGGTTTCTGAAATTTGAAGCTTGAAAAATTGATATTGAAGTAGGAACTGGGCTCAAAATTACAACAAATTACAGTTAAGAAAAGATAACCATTTGCGAGATATTGATGGAAAAATAAAAATAATTCAATCAGTTCTAATTGAATTTTTGTAGGAGTTTATTGAATTGGAAAAAAATTAAGTAAAAAAAAAAAGTTCAACTTTTTCCCATTCATTTCTGTGCACTATGCAAGTTTATTTTGGATTAATAAAAAAAAACTACACGGCAGTACCTTTTCCAAAAACTGAAATCGTTCCTTAATCATTTTTTGTTAGGAAATGTTTTATATGGCTTAATTTTTCATTAGTTTGAGAACCAGAGAGCATCCTGTCAATTTCCTTTTTTGAGCTCATCCTCCTTGTTTTCCTGCTCTTCTTTTTTTCTATTTAGCAAATAAATTATATGATGATAAATCAGTGTGAAGGAGCCGCCCATGGTTGTCCCGAAAGAACAGCCGGGTGTCTACCCCTCTCTTCATTGGACAGATTTTATGCAAATAGTATCATCAGCTTGTTCTCTCCTTCTTCTTCTCCAAGGCATGGTCGTAACCGAATGTGAGGAGACGATTGATGAGTTATCGAGAGTCTGACACGTTCATAACAAAAGAGAAGCAAAATGGTTGAAGATGGGCGGTGAGGAGCAATTCATTCACAATAACTTAATCCGTCGACAGATTTTGAGGGGAGGAGCTGATGGGTTCGAGAGGAAGTTGAGAAGGAAATGAAATTTAGATGGCTTCTGAGCCATCGTGAGCAATTATTTTGATCGTTTCATTTTTTTTGTTGCTTTTGCTTTTTTTCACCTTTCATGTTCCTTTCATGTTCCCTTTCATGTTCATATAACTTAATTTCGGGTGGCTTTAATGATACTTCCTTTATTGATCGATTCTAAAAATTAAAATATAACGTTATTATTTAATGATAAGCCATTTTCAGTAATTGCATTTCACAATTATCCTTGTCAAACTAACGCGATGTCTGTTTCTGTCGATTTTTTAAAATTACATTTTTTGAAACGACACCAATTACTGTGGGGTATTTTAGGTAAATTACTGGTTTTATCAATTTTTTCTTGACCTTAACTTTTTTAACTGTATTTCCCATCTTATAAGAGACTATAGTCTTGCACTCCTTCCGATTAAAAATTGTTTTTTGCATTCTCTGATAGTAGTTGCCGTATTTTTTGCAGATGTCAAGTCCCTCACTTTTTAGTCAAATAGACAAGCTGTATCGAATCAATACTTTGGAAAACTAAGAGACATTCTAGAATTAATTTTGCACCAACAATTCATACATGACTAATAAACATTTTTTGTGAATACCTTTCCATTTTGAACATTTAGTAGAGTTGGAAAGCTTGTTGATTTGATTTGAAAAAGAAAACAATAAAATTGACTTTTCCCATCTTTGCCCACCTGTGATGGGACCTCATCATCATCATGACTTTATGATATACAGTGTTGCCCCTAAACATTTTACTACCCACCACAAAACTTAACGATGTGTGGCAACAGACACGTTTATGATTGGTGCAAGTGTCGATAAAACTGTACGCGTAACATCATCAGTACTTGCAAAAAATATACAGCAACTCACTTCATCAATCTCTATTCTTGACATATTTGGTGTTGTATTGGTTCCACCGTTCCTAGTTCTTGCCTTTTGAATGGAAGATGGTTATCTTGAAGACATGTACACAAGACTGATGACGTTGCAAGACGGGAATACGTGGCAAAGCGGCTTCTCTCTTCTTACAACACCTGAATTTTTAATCGTAGTTTTGATAGTATTTCTACATCTATCTAAATTTTTTTGGGGTATAAATTTTTTTCCTATCAAAAAAATTATATAAAAATTATTTTAGAAGTAAAATTAACAATAAAACATAGAAAAGTTATTTTTTCAATACTCCAACTCAAAATTTATACCTACGTTAAAGTTTCCTAAATATTCGAAAGAAATTATTTATTGAAATTATAGTTTTTAACTCAATTAACAAAAAAAACGAACGTATTTGTGCTGATAGAAAATAAAAGTTTTCCCTAAATAGTTCGTTAACATATGTTGTTCTTTTTTTTTAAACATTTTAGTAATCTGAAAAATTTCAAAGTCCCCTGAATTTGAACCCTGCCTGATGTCCCGTTACCTATCAATTCAAAACCCCATGGGGTATTTACTTTCACCCACACTTCATGCTAAAACTTCAATTATTATGATGTGCTTCACCCGTTTAGTTTTGAGTTGATCGACACCCTTCACACCGATAACCCCGAGGTCAATTCTCTAAAAGTCCCGGGGAGCCGCCAAACGTCTGTCACCTATAATTGTGATGTCCGGAAGCTATGAATCTCGATTAAGAGTCGGCTATTCTGTTGTTGTGGTTTAGGGATTGAAAAGAGGAGATAGTCCCCGGGGAACTATCTACTCTTCGTTTATTTTCCATTTCTCTTTAGCTCTTTTGATTCTATTCGAGACGAATCAAAAACGAGAGGGGGAGGAGGATCATTGAAGAAAGTTAGAATGACTAATCGTGCATTTATTGAAATAATGAAGATAACTTTTACCTATAACTTTATGATTTTTAACATGACCCATAATTTGAAATCTTAGTGTTTGCATACACTTGTAAACTTTTCTGTTTTATTTGAAGGTAATCTACCTTCTTTCTCCCTTTTGCAGTTTTTTTCGTCGCAATTCCAATTAGTCAATTCTAGAAACATCGTATCTTTTTTCCAATGGCATGCCTGACATAGTGTGAACACTTTTTTTTCAAATTCCGCTCACTGACACCTTTCAAAGCGTGCGGTGCCTGCCTGACAGTCAAACCTGGAGATGGCCTCTTTAATTTCTTTGAACTAAGCGCTCAGAAGATGTGTTACCGGAAAGGTGACAGAGCAACGGAAGGAAGATACATAATATATGTGGACGAAACCGGTGGTCGTCTTCTTTTATTGATTTTTTTAGACGGTTGCCGTTTGAATTTCAATAAAGGTGTATGACAAACGAACTTTAAACTATTTGGCATTCTGAAACTAAAAGAAAATTTGAATTTGAATAATTTCAATAGTAAACTGACATTAAAATTCAAGTAATTATCAACTAGTTTCATCAATTAGAAATTTCTAGTAGATTTTTTTCGCTGTTCTTCAATAAATGTTCCAATAATATTTTACTAATACCATCTCCACTTGTTCTGCTCACACGTGTTCTTCTATCACTTGTTTCTTTATTCAAATTACATCTTCTCTTATAAACAAAGAATTACACCCGATTTCGCACCTCGAGTGAGACTGGTATGAGAGCATAGAGACACGCGCAAAAACCAAAAATTTTACGAGCAACAAACAAATTTTTCAAGACTTAGGTGAACTCTTGCAATTTGAGAAGTATCAGCTGGCATTTTAGGATTACGCGTTCCGCATGCTCACAACTATTTTGCTGTTTCAAAAATGCAAAATAAAGATGAACGTAGATGAATATACCGTGTTTAATTTTTGTTCTTCTCTCTTGTGGACATACAAAATTAAGGCAGTGCTAATAGAAAAAACGAAGATACGAAAAAACATCATCGATACCTTTTTACCGTTTTACCACCTTTAACCGACATTATGGAGATGCCAGGCAATAAAAACTTTAACAATGACGATGTAACATTAGAAAATATATCAGAATATATACTTTCGAAAACAAACATCCATAACATCCAATGTCACGCATTCATCATAAAACATTGGCAAAAAGGGCACTGAAATGAGTGGCATCAGATGGAAATCTCAAGAAGATTGCCGAGTCGCGATACAGTGCAGTATGTCATTTGCGGTAAAAAGCGGCTACCTCTCTGACTTCCTGTCACACCAATCACCACCACAACCACCACCTAAAGAGCAATAAATCAGTGGAGGGGGAGATAGTGCGAGCTCGACGCGTGGGGTGTCAGGGTTCCTGAGGAATTTAAAGTGGATCGAGGAGACGGGGACAGGCGCCAGTGCTTTCATCTCTCCGCCCTTTTCTCTTCATCCGAAGTATGGAAATGGATTAGTGTGATTTTGATGGCATACTGACTGCGCTAATTGATTTTGCGGTAGACATTGTGAGCATCTATTTTGAGTCTCTCTAACTTTGTCAAAATCACCTTATATTCGGAATAGATTATTAGATTATTAAAGCTAACGATTCTACCCTTCTTCCATACTAGTTTCTGAGCTCCAACTGTTTGTAGAAATTTCTCTAGAATATCATGTTGTCTAGTCAAATACTTACTAATTTATATCCCCCCATTCAATTCATCTCGTATTTGTTTCTTCGAAACATTGAATGTAGCCGAGCTCATTTCTTCGTTGTGCTGCTAAACACAAAAAAGAAGCTCCTTCTTCTCGTTTGCCTCTCACTCCTGCCATGATTTAATGGGGTGTTCAATGAAGCGAGGAATGAGCTCAGAGGATAGGCTACGGTAGTCAAGAAATAAGAAGAAGCTTGTCTTTTTGAACATCCGGTGGGAATTCGAGACGATGAAACTGAACACAGACAAAATGAATCCGAAGCAACGTGTATTCACATCATTTATTCTGGCGAATTCTTGGAGAAGAACCTAAACGTATTTTATAAGAAAAGGATTACGGTATTCGATTTGCCTTTTTGGCAGAGTTCAAAAATTAAGCTTCAAAAGAATCTAAAACTCAAAATAAGACTGCCAAAAAAATGACCCAATCAGGCATTTTTTGAACCACAACATGTAACACTTCCGCTATGATCTTTATATTGTGATAAGATCCCAATCTGTGATCTGATAATGTCCAATGTGATAAAGTCCGCTCGATAGCGTTTCGGTTTTTAATCTGTCACACCTATCCTCATGTTCACACTTTGTTTTCTTATCCGACCTACTGATCTAGATTCAGTTTTACCACTGAAAGTTCTAACAAATTACCACCATCTGGTGGAGTAATTTCAATACTTTCACTGTTTTATTCACTGTATTCACTGTGTATTCACTGTATTCACTGTGATTTCACTGTATTCACTGTTTTATTTATTAGAGTTATTACCTTGGTTAAAAAGTATATAACATAAGATCTAGAATATTCAGAGTTTTGATTTCTTGTTTTATTCTTAGAAGACATTCCACACCTAAAAATTTTTTGACCTATCTTTCCCGCCCAAACTCCAATTACTGCTTCGTTCGTTTTCAGGAATTATTTTTAATCTTTTTGTGTAATTTTTCATTATTTTGAGTGCCAACTGTGTTGTCAGATGCTCAGTTATAATTATCGCTTGGCTCAAAATATTGCTCTCACCATTTTATCGCTTGAATTCGAGTTTTTTGACATAATAGTCACTTTGCTCGCATTTTCCAATGAAAGATGAATAATTTTTTTTATTTTTTATTTTTTTCATTTTTTAAGTTTTTTTAATTAGAAAATCAATATACCACGCTTCGATAACTTTTAAACATTTTTTTCACGGACCACGTGGAGTTTCAGCAAAAAGATTATTTCTTATTAAAAAATGAACTTCTCATGAACTTTCGTCATTAGTTCCACAATTTCCTTCGACAAGTCTCCACGAAATTAAACAATTTGTTGTGATCGACAGTCTAAATCATCTTGAACTGTTTGAAGACGGTAAGCCCTCGTAGCCTAGATGTCGAGGCCAATTAATCTAATTTTTATAGTAAGAGGATA

**P*lin-26***: epidermis (4072 bp):

CGAGGTTAATATCTGAGCTCCATTGAGCAATATAAAGACTATTATGTAAGGTATCCCACATATCCCATTGGTGAGCATAGCTATTCTATCCGTCGATAATACCTCGTGCATATCGCTAAGACCCGTTTGCTTCCGTGTTTTTCATCTCCTCACACATTCTTCGTTGCAACTTTTCATCTGTGTGCGCGAAGGCGCATCAGATGATCGTCTTCTTCGGCTTCTTCTCCCTTTCTGCTACTTCCATTCAATTCGGCGCACACAAAACAGAAGCACTCTCATCCTCAAACCGGCTTGCCAGTCACCCACGAGCGTACCATCTCTCGGGGGCCACCGCTCCTTCGCCGCTCGTGTCCTTATTACACTCCGACGCTCACAGTTCTTCACATTTTTCCTCTTCTTTTACCACTTCTTCTTCTTCTGAGAAGAAGACGTTGCCAATTTGAATGCGCGACTTCTTGTGTGTATGAGCGCGCTCTGTAACGAAGCGATGCAGAAGCATGAGTCTCTGCTGTGGCCCAAATTGCTTGCCATAACAATTTCGGGCCGTCGCCGACAGCGGTAGCAACGACGCCATTTGCTTTTTGCTTATCACGAAGTGAAGCAAGAACCGGTTGGCACTTGTCTCAGACACCACCATCAAGTGCTTGCTCGTCTTCTCTTTTCGACTGTCAATGTCTTATTCTTTCTTCTTCCTGCTACTTATGTAGTACAGTATATTGGTTCTACTTCCTCTAAAAACTTACTGTTGAGATTTTGGTTAAACATTTGTTCCATCCGTGTAATCATATTTAATTTCAAGAAATCCCAAACAACGTTCACCTTTTTCTGGTCTCCATTTCTCTTTACCCGAGCATTTTCTTATCAGACGAGTTGATCTACTTTCTAGCGGTTGCATGCCCTTGACTCTTGATGGTGTTCGGTAATTCTAACCGTAAACTTGATTTAGGATAATTGCTCAATATCCTTCAACTTATTGTTAGACATCTATGTTATTGCTGAAACTGTGTAGTATTTGTCTGACGGTTGTCGCTCCTTCGTTGAGCGAAGAATCATTTGCCTCAGTCGATCTAAAACTCGTTGTGTAGGGGGTGACAGGCTCCTAAAAGAAAGAAGGAACCGCCCAACTTGTCTCTTGTGGCGTCGTCTCGCACACTATTTCTGTGCCCCGTCGACGATGCGGACGTACAGAAGAAGAGCCAAAAACGAGTTTTCTCTAGATGGACGCGGTGGGGGAGAAGGAGTAGAAGATGGAGGCAACATAGTCTCTATCGGAAGACGAAGAAGACGTTTCTCATTCCTCGACGTGTGATGTCTCTCTATTCGCCACAATCTCTACTGTCCTTCAGTGGCGTACGACCACGATGTAAACAGCACAGCGCCCCCTAGAAGACAGCGGGATACTGTAGAAAGAGAGTGCTCGGCGAGTAGTGCGCAGAGAACTTTGATTGGGTAATTTATCGAATGATAACCGCACGAAGTCTGTTAGTTTTTGAGAAAAAAGAAGGAAAATAGCGTCTTCCTTGGTTTGTCGATTGCGATAACTATCTAGATTTGAACACATTCACAAACAGATACTGATCAATTTCATTTTTTTAATCATTTTGGCACGAAAAACGTCACTCTCCTGTCTCGAGAGTTAGTTGACACATTCCTTCATTTATTTTTCACTTCTTGAAATGCAAATTGTTATCCCTCAACCAATCCCATCAATCGTCGCCATTTTGTCGTCCGTTTCTGGGTCACCGGAGCGGCGGTAGTTCCTGATAATTGATATGTGCGCAGTTTACGACACACACTCTTTGCAAAGACCTTGAAGAGCTCTGTCATTATCAATCAATCGCCCGTCTTATATCCCACCCAGATATTTTTTTCTGCAAATTTCTCGTTCATTTCCATTAAAACATTTGAGCCGGCTCCGCCTCTCTTTTGTAGAATGAGTTTCGCTCTTCCACGGAACGCCACGCCCATCATCTCAACGTAGGTTCCATAGCGTTCAGTCGTCACTATTTCGAATCATGAGATGGTTACTGTCGTGACCCTGTCGGAGATTGGAATCTCGGAGCTCGCTGAGTTGGCAATGTGTCTACTCGTCCATCTATTCTTTCTTATATGATTACTGCCTCTTTCTCTATAGAATCTCTTCATTTGTCCTTGAGTCGCAACAGCCATCTCACATGATATTCGGGTGAATCATCGTTTAATTTCTTGTATTTCTTGCTCTTTTTGTTTGTGAAAAATTGTTTTGACAAATTTCAGTAGTATGATAATATCAATTCAAATTTATTTGGAAACCCCTCATAAAAACATCACGTTGTAGATAGTTCATAATATCAATTAGTTATGAATGTATTATCTCCAATTTTTTTTCTAAATTTAATCACTGCCAAGTTCAAATATCCTCAAGATATTTTAACACCTCATCCTTATTTTCTAATATCTTTTATCCGACTCACCAGTATTTTCTCTTTCTTTCTCTCGTATTCGCTCACCTTGATCATCTACAGCTCTTACCGTAACTCCGCCCCTACTCCGCTCCACCCCTATACGCGCTCACGCACAAACCCGCATTCAGTGGCCTTCAGCCTCGGAGCGCTCTACCTGCGCTCTCATGTGAACCAACATTCTTGTTTGAGCTCCTTCTTTTTCTTGTATTTTTTTTAATATTCTGAATTTCAGGTGAAAAAGAATTTCGAACGTGTCTCGCTGCTCACACGTCTCGAAGATCACCCGTATCTTCGTGATGATCAAGATGTCGTTCAAGACGATACACAGTCTCAAGGAACCGATGGAATGCTTCTTCAGCTCAATGATGAGGAACGACAAAGGCTAATGAAGATGATAAGCACACCTGGACCGCAGACATTGAGCATGGAACGGATACAGATGGCTCCAGGACAAGGTCAGCATATTCTCCAACAACATCAGATTCCATCAACATCGTCATACACAAACGTAAGTGGTCACTTCTTTGACATTTTAAATATTTTATTTACAGTTAATTTTATAGTGAAAACTGATAAAAACAAGAATGATAATTTTTAGACCTTTAAATTAAAAATATAATGTTTTCTAAAAAAATAAATAATATAACAAAAAAATAAACTGTTTCAGAATTCGTTTATCTACATCAAAAGTGATCCACAATACAAAGATGAACCACAACAACAATCCAATCAATTCGATGCCACTGTGAACTCTGTTGCTCATGGACAAGCTGGACTTTCAACACTAGAAATGTTCGCTTGTAATGGATCTCCGAACGGTGGTGATGTACTTCGCAGAAAGACGCCGTCTCGGAAAGCTGAAGATTCAATAAAAGCTACGTGTAAAGTATGTGGACATGATGTAATGTATTCTCCTCAAAGGACTTGGAACTTGATGCGTCACGTATGGATTATGCACCAATCATCAAAACCGAATCAGTGTTCAATATGCGGATATTCTCACATCAAGCCATATGTTAGGAAACATATCGAGTCTCAGCATAAGCATGATTCAAATGCAACGATAATTGACTTAAAGTGAGTTTTACATTTCTCAGTTTGATAAAATTGTCAATTTGATTTTCAGATCACCAGAACTAGAAGCTGAATGGAGTCAATTGTTAGATCAATGCTTCGGAGTCACATATCGAAAGTGGAAGCATCATAAGGAGGATACGCAGCACACCGGTGGGTTACTGTTCGATGACGACGAGTTGAGAGGGTGAGCGAGAGCGTCACCGACATCAACTGAAGACCGCCAGTGTAGTCTTTGCAGCGGGTGATGAAGATTACGGTCACGAGCATCATTTTTCACCGGAGGAAGATGTAATGACAACTTTATGATTCTTCATGGCTTTCACTTGTCGACGTCTTCCCATTGTCTTCCATTCATCTTTCGTCATCTCTTCATATTTCACTCAATCACTGCCTTGTGCGATCATGTAATTTCAAACTATTGTAATTCGTTGAAATTATCAAATAATAAATTTAATCAGGTTATTATTGAATGTTTATGTAACCTATTCCATCTCACAATGAGTTAATACAACTCTTGAATACTTGAATACCTTCATCCCTCGCTTACTTTT

**Pmex5**: germline (488 bp):

AAATATCAGTTTTTAAAAAATTAAACCATAAAACAAATAATATAACCCAATTTTTACATCAAACCACAAGAAAAAAATACATTTGGGCCCACGGATAAAGAAATTAAAAAAATACATTTTTTAAAGGCGCACCGAATTAAAATTCATTTGGGTCTTACCGCGTATACCGTACTCCGTTTGTTTGATCATTTTTGTCAGCGCTGGCGGTTGTTTTTTCATTTCATTTCTGCTTCAAAGACGTTTTCTCGAATAATTTTTCGTTTATTCTCTTTTTTAAAATTAATTTCTAGCCGTAAATGTTATAAATTCACCCATTTAACGCAAATTTCATGGTAATCTCATGGAAAAATGCAGTTTCTTTGTTAAAGAAAGCTTAAATAGCAAAAATTCCCCGACTTTCCCCAAAATCCTGCTCGATTTTCCGTTTTCTCATTGTATTCTCTCTTAATTAATTTTATCGATAATCAATTGAATGTTTCAGACAGAGA

**P*rps-27***: pan (841 bp):

TTCAATCGGTTTTTCCTTGCTTGCGCCCAATTCTGATGGTTCTTCCATTTCTAAATTTTAGTTTAATAAAAAAATTTTAAAATAATACGAGTTTTGAGACTAACTTTCAATAAAAACCAAAAAAAATTGCTTCAAAAAAGAGTTCGCGCCGATTTACGATAAACACAGCGCGCTACGACACTCCGAAAACCGCGCCGCACCATTACATTCTCCGCCCCACCGCACGGTTTGCGGAAGGCGATGAAATTGCTGGGCCTAAAATTTTAACCCTTTTTTTCCCGTTTTTCAACTGTTTTCCCGATTTTTTCGATATTTTTTTCAGGTTTTTAGTTAGAAAAGTATATAAAAATTGCATTATTTGTAGTTTTCGCTCGTAATTTGTGATAGAAAACGGCCAATTTGCCTAGATTTGTGATTCTACCAAGTGGAATTTTTGGGAAAATATCAATATTCCACGATTTTTATTCAAATCATAACGTTTCGAAATTATCAGCTTAAATAAAACTAAAATAAATAATAAAATACTAAAAACCCTGAATTTCTCGGTTCTCGGTTCAGCTTGAGTTCGTTCTATACACGTTATCGATTTTTGTCATTTCGTTTAATCATGGGTTTTGATTCATTTTAAATATGAAATTTGATTTATTCCTCGTTTCGAAGTTGGTTTGTCTTGTCCGGCCGGTAAACAAATGTTATTAAAAAAATAATAAATTTCGAAATTTTTCGTAATGTTCCGCAATCGTGTTTTTCCGTTTATCTCGTCTTCCCTTCTTTCAAATATACTATTTATTTTCTCAATTCACATATTTTCAGCCAGCCCCGCTCAACAAGTGGAATAAAA

**P*eft-3***: pan (600 bp):

GCACCTTTGGTCTTTTATTGTCAACTTCCATTGGTTCTTCCATTGTTTCTGTTAAATTAATGAATTTTTCATAAAATAAAGACATTATACAATATAAAAATGAAGAATTTATTGAAAATAAACTGCCAGAGAGAAAAAGTATGCAACACTCCCGCCGAGAGTGTTTGAAATGGTGTACGGTACATTTTCGTGCTAGGAGTTAGATGTGCAGGCAGCAACGAGAGGGGGAGAGATTTTTTTGGGCCTTGTGAAATTAACGTGAGTTTTCTGGTCATCTGACTAATCATGTTGGTTTTTTGTTGGTTTATTTTGTTTTTATCTTTGTTTTTATCCAGATTAGGAAATTTAAATTTTATGAATTTATAATGAGGTCAAACATTCAGTCCCAGCGTTTTTCCTGTTCTCACTGTTTAGTCGAATTTTTATTTTAGGCTTTCAACAAATGTTCTAACTGTCTTATTTGTGACCTCACTTTTTATATTTTTTTAATTTTTAAAAATATTAGAAGTTTCTAGGATAATTTTTTCGACTTTTATTCTCTCTACCGTCCGCACTCTTCTTACTTTTAAATTAAATTGTTTTTTTTTCAGTTGGGAAACA
