## Supplemental Table 2 for "A tissue-targeted auxin-inducible degradation toolkit for *C. elegans* embryogenesis"

**Table S2. List of strains used in this study**

***** *The strains recommended in Figure 2 are available from the Caenorhabditis Genetics Center*

| **Tissue(s)** | **Promoter** | | **Strain** | **Genotype** |
| --- | --- | --- | --- | --- |
| **Strains carrying the TIR1F79G—mCherry H2B operon as an extrachromosomal array*** | | | | |
| Intestine | | *Pelt-2* | OD5648 | *ttTi5605 II; unc-119(ed3) III; ltEx19[* *pOD5555/pNV36; Pelt-2::AtTIR1[F79G]::*operon linker::mCherry::linker::*his-11::tbb-2 3'UTR; cb-unc-119(+); pRF4 (rol-6(su1006)]* |
| Pharynx | | *Pcest-17* | OD5981 | *ttTi5605 II; unc-119(ed3) III; ltEx33[ pOD5562/pNV29; Pcest-17::AtTIR1[F79G]::* operon linker::mCherry::linker::*his-11::tbb-2 3'UTR; cb-unc-119(+); pRF4 (rol-6(su1006)]* |
| Muscle | | *Phlh-1* | OD6094 | *ttTi5605 II; unc-119(ed3) III; ltEx42[ pOD5557/pNV39; Phlh-1::AtTIR1[F79G]::* operon linker::mCherry::linker::*his-11::tbb-2 3'UTR; cb-unc-119(+); pRF4 (rol-6(su1006)]* |
|  |  | *Pmyo-3* | OD5645 | *ttTi5605 II; unc-119(ed3) III;* ltEx21[pOD5556/pNV38; Pmyo-3::AtTIR1[F79G]:: operon linker::mCherry::linker::his-11::tbb-2 3'UTR; cb-unc-119(+); pRF4 (rol-6(su1006)] |
| Neuroblast | | *Pztf-11* | OD5978 | *ttTi5605 II; unc-119(ed3) III; ltEx36[pOD5563/pNV54; Pztf-11::AtTIR1[F79G]*::operon linker::mCherry::linker*::his-11::tbb-2 3'UTR; cb-unc-119(+); pRF4 (rol-6(su1006)]* |
| Neurons | | *Pcnd-1* | OD6317 | *ttTi5605 II; unc-119(ed3) III; ltEx58[* pOD5564/pNV73*; Pcnd-1::AtTIR1[F79G]::* operon linker::mCherry::linker*::his-11::tbb-2 3'UTR; cb-unc-119(+); pRF4 (rol-6(su1006)]* |
|  |  | *Pdyf-7* | OD6326 | *ttTi5605 II; unc-119(ed3) III*; ltEx67[ pOD5565/pNV66; Pdyf-7::AtTIR1[F79G]:: operon linker::mCherry::linker::his-11::tbb-2 3'UTR; cb-unc-119(+); pRF4 (rol-6(su1006)] |
| Epidermis | | *Pelt-1* | OD5681 | *ttTi5605 II; unc-119(ed3) III*; ltEx24[ pOD5554/pNV16; Pelt-1::AtTIR1[F79G]:: operon linker::mCherry::linker::his-11::tbb-2 3'UTR; cb-unc-119(+); pRF4 (rol-6(su1006)] |
|  |  | *Plin-26* | OD6320 | *ttTi5605 II; unc-119(ed3) III*; *ltEx61[* *pOD5559/pNV74; Plin-26::AtTIR1[F79G]::*operon linker::mCherry::linker:*:his-11::tbb-2 3'UTR; cb-unc-119(+);* pRF4 (rol-6(su1006)] |
| **Strains carrying the TIR1F79G—mCherry H2B operon as an integrated single-copy transgene*** | | | | |
| Intestine (Mos II) | | *Pelt-2* | OD5691 | ltSi1933[*pOD5555/*pNV36; Pelt-2::AtTIR1[F79G]:: operon linker::mCherry::linker::his-11::tbb-2 3'UTR; cb-unc-119(+)]II; unc-119(ed3?)III |
| Pharynx (Mos II) | | *Pcest-17* | OD5774 | ltSi1952[pOD5562/pNV29; Pcest-17::AtTIR1[F79G]:: operon linker::mCherry::linker::his-11::tbb-2 3'UTR; cb-unc-119(+)]II; unc-119(ed3?)III |
| Muscle (Mos II) | | *Phlh-1* | OD5800 | ltSi1956[pOD5557/pNV39; Phlh-1::AtTIR1[F79G]:: operon linker::mCherry::linker::his-11::tbb-2 3'UTR; cb-unc-119(+)]II; unc-119(ed3?)III |
| Neurons (Uni I) | | *Pcnd-1* | OD6313 | ltSi2067[oxTi185; pOD5564/pNV73; Pcnd-1::AtTIR1[F79G]:: operon linker::mCherry::linker::his-11::tbb-2 3'UTR; cb-unc-119(+)]I; unc-119(ed3?)III |
| Epidermis (Mos II) | | *Pelt-1* | OD5694 | ltSi1935[pOD5554/pNV16; Pelt-1::AtTIR1[F79G]:: operon linker::mCherry::linker::his-11::tbb-2 3'UTR; cb-unc-119(+)]II; unc-119(ed3?)III |
| Germline (Uni I) | | Pmex-5 | OD6102 | ltSi2035[oxTi185; pOD5566/pNV43; Pmex-5::AtTIR1[F79G]::3xFlag::tbb-2 3'UTR; cb-unc-119(+)]I |
| Germline (Mos II) | |  | OD5834 | ltSi1961[ pOD5566/pNV43; Pmex-5::AtTIR1[F79G]::3xFlag::tbb-2 3'UTR; cb-unc-119(+)]II; unc-119(ed3?)III |
| Pan (Mos II) | | Prps-27 | PLG128 | ccpSi49[pLA095; Prps-27::AtTIR1[F79G]::3Xflag::operon linker::mCherry::linker::his-11::tbb-2 3'UTR; cb-unc-119(+)]II; unc-119(ed3?)III |
| Pan (Uni I) | | Peft-3 | PLG297 | ccpSi100[oxTi185; *pOD5558/pNV44/pLA179*; Peft-3:: AtTIR1[F79G]::operon linker::mCherry::linker::his-11::tbb-2 3'UTR; cb-unc-119(+)]I; unc-119(ed3?)III |
| Pan (Mos II) | |  | PLG272 | ccpSi90[ *pOD5558/pNV44/pLA179*; Peft3::AtTIR1[F79G]:: operon linker::mCherry::linker::his-11::tbb-2 3'UTR; cb-unc-119(+)]II; unc-119(ed3)III |
| **Strains carrying *in situ*-tagged CRISPR alleles** | | | | |
| - | | - | PLG28 | mdf-1(ccp8[AID*::GFP::3xFlag::LoxP::mad-1]V |
| - | | - | OD6264 | let-502 (lt273[GFP::let-502])I |
| - | | - | OD6238 | let-502 (lt268[AID*::GFP::let-502])I |
| **Strains carrying the TIR1F79G—mCherry H2B operon as an extrachromosomal array along with *in situ*-tagged AID::GFP::MAD-1** | | | | |
| Intestine | | *Pelt-2* | OD5660 | *ttTi5605? II; unc-119(ed3?)III; mdf-1(ccp8[AID*::GFP::3xFlag::LoxP::mad-1])V; ltEx19[ pOD5555/pNV36; Pelt-2::AtTIR1[F79G]*::operon linker::mCherry::linker::*his-11::tbb-2 3'UTR; cb-unc-119(+), pRF4 (rol-6(su1006)]* |
| Pharynx | | *Pcest-17* | OD6274 | *ttTi5605? II; unc-119(ed3?)III; mdf-1(ccp8[AID*::GFP::3xFlag::LoxP::mad-1])V; ltEx33[ pOD5562/pNV29; Pcest-17::AtTIR1[F79G]::* operon linker::mCherry::linker::*his-11::tbb-2 3'UTR; cb-unc-119(+); pRF4 (rol-6(su1006)]* |
| Muscle | | *Phlh-1* | OD6151 | *ttTi5605? II; unc-119(ed3?)III; mdf-1(ccp8[AID*::GFP::3xFlag::LoxP::mad-1])V; ltEx42[ pOD5557/pNV39; Phlh-1::AtTIR1[F79G]::* operon linker::mCherry::linker::*his-11::tbb-2 3'UTR ; cb-unc-119(+); pRF4 (rol-6(su1006)]* |
|  |  | *Pmyo-3* | OD5661 | *ttTi5605? II; unc-119(ed3?)III; mdf-1(ccp8[AID*::GFP::3xFlag::LoxP::mad-1])V; ltEx21[ pOD5556/pNV38; Pmyo-3::AtTIR1[F79G]::* operon linker::mCherry::linker::*his-11::tbb-2 3'UTR ; cb-unc-119(+), pRF4 (rol-6(su1006)]* |
| Neuroblast | | *Pztf-11* | OD6150 | *ttTi5605? II; unc-119(ed3?)III; mdf-1(ccp8[AID*::GFP::3xFlag::LoxP::mad-1])V; ltEx36[* *pOD5563/pNV54; Pztf-11::AtTIR1[F79G]::* operon linker::mCherry::linker::*his-11::tbb-2 3'UTR ; cb-unc-119(+); pRF4 (rol-6(su1006)]* |
| Neurons | | *Pcnd-1* | OD6357 | *ttTi5605? II; unc-119(ed3?)III; mdf-1(ccp8[AID*::GFP::3xFlag::LoxP::mad-1])V; ltEx58[* *pOD5564/pNV73; Pcnd-1::AtTIR1[F79G]::* operon linker::mCherry::linker::*his-11::tbb-2 3'UTR; cb-unc-119(+); pRF4 (rol-6(su1006)]* |
|  |  | *Pdyf-7* | OD6362 | *ttTi5605? II; unc-119(ed3?)III; mdf-1(ccp8[AID*::GFP::3xFlag::LoxP::mad-1])V; ltEx67[* *pOD5565/pNV66; Pdyf-7::AtTIR1[F79G]::* operon linker::mCherry::linker::*his-11::tbb-2 3'UTR; cb-unc-119(+); pRF4 (rol-6(su1006)]* |
| Epidermis | | *Pelt-1* | OD5686 | *ttTi5605? II; unc-119(ed3?)III; mdf-1(ccp8[AID*::GFP::3xFlag::LoxP::mad-1])V; ltEx24[ pOD5554/pNV16; Pelt-1::AtTIR1[F79G]::* operon linker::mCherry::linker::*his-11::tbb-2 3'UTR ; cb-unc-119(+), pRF4 (rol-6(su1006)]* |
|  |  | *Plin-26* | OD6333 | *ttTi5605? II; unc-119(ed3?)III; mdf-1(ccp8[AID*::GFP::3xFlag::LoxP::mad-1])V; ltEx61[* *pOD5559/pNV74; Plin-26::AtTIR1[F79G]::* operon linker::mCherry::linker::*his-11::tbb-2 3'UTR ; cb-unc-119(+); pRF4 (rol-6(su1006)]* |
| **Strains carrying the TIR1F79G—mCherry H2B operon as an integrated single-copy transgene along with *in situ*-tagged AID::GFP::MAD-1** | | | | |
| Intestine | | *Pelt-2* | OD5693 | ltSi1933[ *pOD5555/*pNV36; Pelt-2::AtTIR1[F79G]:: operon linker::mCherry::linker::*his-11::tbb-2 3'UTR*; cb-unc-119(+)]II; unc-119(ed3)III?; mdf-1(ccp8[AID*::GFP::3xFlag::LoxP::mad-1])V |
| Pharynx | | *Pcest-17* | OD5798 | ltSi1952[pOD5562/pNV29; Pcest-17::AtTIR1[F79G]:: operon linker::mCherry::linker::*his-11::tbb-2 3'UTR*; cb-unc-119(+)]II; unc-119(ed3)III?; mdf-1(ccp8[AID*::GFP::3xFlag::LoxP::mad-1])V |
| Muscle | | *Phlh-1* | OD5814 | ltSi1956[pOD5557/pNV39; Phlh-1::AtTIR1[F79G]:: operon linker::mCherry::linker::*his-11::tbb-2 3'UTR*; cb-unc-119(+)]II; unc-119(ed3)?III; mdf-1(ccp8[AID*::GFP::3xFlag::LoxP::mad-1])V |
| Neurons | | *Pcnd-1* | OD6359 | ltSi2067[oxTi185; pOD5564/pNV73; Pcnd-1::AtTIR1[F79G]:: operon linker::mCherry::linker::*his-11::tbb-2 3'UTR*; cb-unc-119(+)]I; unc-119(ed3)III?; mdf-1(ccp8[AID*::GFP::3xFlag::LoxP::mad-1])V |
| Epidermis | | *Pelt-1* | OD5696 | ltSi1935[pOD5554/pNV16; Pelt-1::AtTIR1[F79G]:: operon linker::mCherry::linker::*his-11::tbb-2 3'UTR*; cb-unc-119(+)]II; unc-119(ed3)III?; mdf-1(ccp8[AID*::GFP::3xFlag::LoxP::mad-1])V |
| Germline | | *Pmex-5 (Uni I)* | OD6147 | ltSi2035[oxTi185; pOD5566/pNV43; Pmex-5::AtTIR1[F79G]::3xFlag::tbb-2 3'UTR; cb-unc-119(+)]I; unc-119(ed3?)III; mdf-1(ccp8[AID*::GFP::3xFlag::LoxP::mad-1])V |
|  |  | *Pmex-5 (Mos II)* | OD5876 | ltSi1961[ pOD5566/pNV43; Pmex-5::AtTIR1[F79G]::3xFlag::tbb-2 3'UTR; cb-unc-119(+)]II; unc-119(ed3)III?; mdf-1(ccp8[AID*::GFP::3xFlag::LoxP::mad-1])V |
| Pan | | *Prps-27*  *(Mos II)* | OD6272 | ccpSi49[pLA095; Prps-27::*AtTIR1[F79G]::* 3Xflag::operon linker::mCherry::linker::*his-11::tbb-2 3'UTR*; cb-unc-119(+)]II; unc-119(ed3?)III; mdf-1(ccp8[AID*::GFP::3xFlag::LoxP::mad-1])V |
|  |  | *Peft-3 (Uni I)* | OD6467 | *ccpSi100[oxTi185;* *pOD5558/pNV44/pLA179; Peft-3::AtTIR1[F79G]::operon linker::mCherry::linker::his-11::tbb-2 3'UTR; cb-unc-119(+)]I; unc-119(ed3?)III?; mdf-1(ccp8[AID*::GFP::3xFlag::LoxP::mad-1])V* |
|  |  | *Peft-3*  *(Mos II)* | OD6407 | *ccpSi90[pOD5558/pNV44/pLA179; Peft3::AtTIR1[F79G]::*operon linker::mCherry::linker::*his-11::tbb-2 3'UTR; cb-unc-119(+)]II; unc-119(ed3)III?; mdf-1(ccp8[AID*::GFP::3xFlag::LoxP::mad-1])V* |
| **Strains carrying the TIR1F79G—mCherry H2B operon as arrays or integrated single-copy transgenes along with *in situ*-tagged GFP::LET-502** | | | | |
| Intestine | | *Pelt-2* | OD6299 | let-502(lt273[GFP::let-502])I*; ttTi5605? II; unc-119(ed3?)III;* ltEx19[*pOD5555/*pNV36; Pelt-2::AtTIR1[F79G]::operon linker::mCherry::linker::*his-11::tbb-2 3'UTR*; cb-unc-119(+), pRF4 (rol-6(su1006)] |
| Pharynx | | *Pcest-17* | OD6308 | let-502(lt273[GFP::let-502])I*; ttTi5605? II; unc-119(ed3?)III;* ltEx33 [pOD5562/pNV29; Pcest-17::AtTIR1[F79G]:: operon linker::mCherry::linker::*his-11::tbb-2 3'UTR*; cb-unc-119(+); pRF4 (rol-6(su1006)] |
| Muscle | | *Phlh-1* | OD6296 | let-502(lt273[GFP::let-502])I*; ttTi5605? II; unc-119(ed3?)III;* ltEx42[pOD5557/pNV39; Phlh-1::AtTIR1[F79G]:: operon linker::mCherry::linker::*his-11::tbb-2 3'UTR*; cb-unc-119(+); pRF4 (rol-6(su1006)] |
| Neuroblasts | | *Pztf-11* | OD6310 | let-502(lt273[GFP::let-502])I*; ttTi5605? II; unc-119(ed3?)III; ltEx36[pOD5563/pNV54; Pztf-11::AtTIR1[F79G]::* operon linker::mCherry::linker::*his-11::tbb-2 3'UTR; cb-unc-119(+); pRF4 (rol-6(su1006)]* |
| Neurons | | *Pdyf-7* | OD6365 | let-502(lt273[GFP::let-502])I*; ttTi5605? II; unc-119(ed3?)III; ltEx67 [pOD5565/pNV66; Pdyf-7::AtTIR1[F79G]::* operon linker::mCherry::linker::*his-11::tbb-2 3'UTR; cb-unc-119(+); pRF4 (rol-6(su1006)]* |
| Epidermis | | *Pelt-1* | OD6294 | let-502(lt273[GFP::let-502])I*; ttTi5605? II; unc-119(ed3?)III;* ltEx24[pOD5554/pNV16; Pelt-1::AtTIR1[F79G]:: operon linker::mCherry::linker::*his-11::tbb-2 3'UTR*; cb-unc-119(+), pRF4 (rol-6(su1006)] |
|  |  | *Plin-26* | OD6337 | let-502(lt273[GFP::let-502])I*; ttTi5605? II; unc-119(ed3?)III; ltEx61[pOD5559/pNV74; Plin-26::AtTIR1[F79G]::* operon linker::mCherry::linker::*his-11::tbb-2 3'UTR; cb-unc-119(+); pRF4 (rol-6(su1006*)] |
| Germline | | *Pmex-5* | OD6307 | let-502(lt273[GFP::let-502])I; ltSi1961[ pOD5566/pNV43; Pmex-5::AtTIR1[F79G]::3xFlag::tbb-2 3'UTR; cb-unc-119(+)]II; unc-119(ed3)III? |
| Pan | | *Prps-27* | OD6300 | let-502(lt273[GFP::let-502])I; ccpSi49[pLA095; Prps-27::AtTIR1[F79G]::3Xflag::operon linker::mCherry::linker::his-11::tbb-2 3'UTR; cb-unc-119(+)]II; unc-119(ed3?)III |
|  |  | *Peft-3* | OD6406 | let-502(lt273[GFP::let-502])I; ccpSi90[*pOD5558/pNV44/pLA179*; Peft3::AtTIR1[F79G]::operon linker::mCherry::linker::*his-11::tbb-2 3'UTR*; cb-unc-119(+)]II; unc-119(ed3)III? |
| **Strains carrying the TIR1F79G—mCherry H2B operon as arrays or integrated single-copy transgenes along with *in situ*-tagged AID::GFP::LET-502** | | | | |
| Intestine | | *Pelt-2* | OD6305 | let-502(lt268[AID*::GFP::let-502])I; *ttTi5605? II; unc-119(ed3?)III;* ltEx19 [*pOD5555/*pNV36; Pelt-2::AtTIR1[F79G]:: operon linker::mCherry::linker::*his-11::tbb-2 3'UTR*; cb-unc-119(+), pRF4 (rol-6(su1006)] |
| Pharynx | | *Pcest-17* | OD6309 | let-502(lt268[AID*::GFP::let-502])I; *ttTi5605? II; unc-119(ed3?)III;* ltEx33[pOD5562/pNV29; Pcest-17::AtTIR1[F79G]:: operon linker::mCherry::linker::*his-11::tbb-2 3'UTR*; cb-unc-119(+); pRF4 (rol-6(su1006)] |
| Muscle | | *Phlh-1* | OD6303 | let-502(lt268[AID*::GFP::let-502])I; *ttTi5605? II; unc-119(ed3?)III;* ltEx42[pOD5557/pNV39; Phlh-1::AtTIR1[F79G]:: operon linker::mCherry::linker::*his-11::tbb-2 3'UTR*; cb-unc-119(+); pRF4 (rol-6(su1006)] |
| Neuroblasts | | *Pztf-11* | OD6311 | let-502(lt268[AID*::GFP::let-502])I; *ttTi5605? II; unc-119(ed3?)III;* ltEx36[pOD5563/pNV54; Pztf-11::AtTIR1[F79G]:: operon linker::mCherry::linker::*his-11::tbb-2 3'UTR*; cb-unc-119(+); pRF4 (rol-6(su1006)] |
| Neurons | | Pdyf-7 | OD6366 | let-502(lt268[AID*::GFP::let-502])I; *ttTi5605? II; unc-119(ed3?)III;* ltEx67[pOD5565/pNV66; Pdyf-7::AtTIR1[F79G]:: operon linker::mCherry::linker::*his-11::tbb-2 3'UTR*; cb-unc-119(+); pRF4 (rol-6(su1006)] |
| Epidermis | | *Pelt-1* | OD6302 | let-502(lt268[AID*::GFP::let-502])I; *ttTi5605? II; unc-119(ed3?)III;* ltEx24[pOD5554/pNV16; Pelt-1::AtTIR1[F79G]:: operon linker::mCherry::linker::*his-11::tbb-2 3'UTR*; cb-unc-119(+), pRF4 (rol-6(su1006)] |
|  |  | *Plin-26* | OD6343 | let-502(lt268[AID*::GFP::let-502])I; *ttTi5605? II; unc-119(ed3?)III; ltEx61[pOD5559/pNV74; Plin-26::AtTIR1[F79G]::* operon linker::mCherry::linker::*his-11::tbb-2 3'UTR; cb-unc-119(+); pRF4 (rol-6(su1006)]* |
| Germline | | *Pmex-5* | OD6306 | let-502(lt268[AID*::GFP::let-502])I; ltSi1961[ pOD5566/pNV43; Pmex-5::AtTIR1[F79G]::3xFlag::tbb-2 3'UTR; cb-unc-119(+)]II; unc-119(ed3)III? |
| Pan | | *Prps-27* | OD6292 | *let-502(lt268[AID*::GFP::let-502])I; ccpSi49[pLA095; Prps-27::* AtTIR1[F79G]::*3Xflag::operon linker::mCherry::linker::his-11::tbb-2 3'UTR; cb-unc-119(+)]II; unc-119(ed3?)III?* |
|  |  | *Peft-3* | OD6404 | *let-502(lt268[AID*::GFP::let-502])I;*ccpSi90[ *pOD5558/pNV44/pLA179*; Peft3::AtTIR1[F79G]:: operon linker::mCherry::linker::*his-11::tbb-2 3'UTR*; cb-unc-119(+)]II; unc-119(ed3)III? |
